# Mapping information-rich genotype-phenotype landscapes with genome-scale Perturb-seq

**DOI:** 10.1101/2021.12.16.473013

**Authors:** Joseph M. Replogle, Reuben A. Saunders, Angela N. Pogson, Jeffrey A. Hussmann, Alexander Lenail, Alina Guna, Lauren Mascibroda, Eric J. Wagner, Karen Adelman, Gila Lithwick-Yanai, Nika Iremadze, Florian Oberstrass, Doron Lipson, Jessica L. Bonnar, Marco Jost, Thomas M. Norman, Jonathan S. Weissman

## Abstract

A central goal of genetics is to define the relationships between genotypes and phenotypes. High-content phenotypic screens such as Perturb-seq (pooled CRISPR-based screens with single-cell RNA-sequencing readouts) enable massively parallel functional genomic mapping but, to date, have been used at limited scales. Here, we perform genome-scale Perturb-seq targeting all expressed genes with CRISPR interference (CRISPRi) across >2.5 million human cells and present a framework to power biological discovery with the resulting genotype-phenotype map. We use transcriptional phenotypes to predict the function of poorly-characterized genes, uncovering new regulators of ribosome biogenesis (including *CCDC86*, *ZNF236*, and *SPATA5L1*), transcription (*C7orf26*), and mitochondrial respiration (*TMEM242*). In addition to assigning gene function, single-cell transcriptional phenotypes allow for in-depth dissection of complex cellular phenomena – from RNA processing to differentiation. We leverage this ability to systematically identify the genetic drivers and consequences of aneuploidy and to discover an unanticipated layer of stress-specific regulation of the mitochondrial genome. Our information-rich genotype-phenotype map reveals a multidimensional portrait of gene function and cellular behavior.

## Main Text

Mapping the relationship between genetic changes and their phenotypic consequence is critical to understanding gene and cellular function. This mapping is traditionally carried out in either of two ways. A phenotype-centric, “forward genetic” approach reveals the genetic changes that drive a phenotype of interest. Conversely, a gene-centric, “reverse genetic” approach catalogs the diverse phenotypes caused by a defined genetic change.

Recent technological developments have advanced both forward and reverse genetic efforts (*1*). CRISPR-Cas tools now enable the deletion, mutation, repression, or activation of genes with ease (*2*). In forward genetic screens, CRISPR-Cas systems can be used to generate cells with diverse genetic perturbations. This pool of perturbed cells can then be subjected to a selective pressure, with phenotypes assigned to genetic perturbations by sequencing. Forward genetic screens provide powerful tools for the identification of cancer dependencies, essential cellular machinery, differentiation factors, and suppressors of genetic diseases (*3–6*). In parallel, dramatic improvements in molecular phenotyping now allow for single-cell readouts of epigenetic, transcriptomic, proteomic, and imaging information (*7*). Applied to reverse genetics, single-cell profiling can refine the understanding of how select genetic perturbations affect cell types and cell states.

However, both phenotype-centric and gene-centric approaches suffer conceptual and technical limitations. Pooled forward genetic screens typically use low-dimensional phenotypes (e.g., growth, marker gene expression, drug resistance) for selection. The use of simple phenotypes can conflate genes acting via different mechanisms, requiring extensive follow-up studies to disentangle genetic pathways (*8*). Additionally, in forward genetics, serendipitous discovery is constrained by the prerequisite of selecting phenotypes prior to screening. On the other hand, while reverse genetic approaches enable the study of multidimensional and complex phenotypes, they have typically been restricted in scale to rationally chosen targets, limiting the ability to make systematic comparisons.

Single-cell CRISPR screens present a solution to these problems. These screens simultaneously read out the genetic perturbation and high-dimensional phenotype of individual cells in a pooled screening format, thus combining the throughput of forward genetic screens with the rich phenotypes of reverse genetics. While these approaches initially focused on transcriptomic phenotypes (e.g., Perturb-seq, CROP-seq) (*9–13*), technical advances have enabled their application to epigenetic (*14*), imaging (*15*), or multimodal phenotypes (*16–18*). From these rich data, it is possible to identify genetic perturbations that cause a specific behavior as well as to catalog the spectrum of phenotypes associated with each genetic perturbation. Despite the promise of single-cell CRISPR screens, their use has generally been limited to studying at most a few hundred genetic perturbations, typically chosen with a bias towards predefined biological questions.

We reasoned that there would be unique value to genome-scale single-cell CRISPR screens. For example, while the number of perturbations scales linearly with experimental cost, the number of pairwise comparisons in a screen—and thus its utility for unsupervised classification of gene function—scales quadratically. Similarly, in large-scale screens, the diversity of perturbations allows one to explore the range of cell states that can be revealed by rich phenotypes. Additionally, as many human genes are well-characterized, these genes serve as natural controls to anchor the interpretation of observations in comprehensive datasets. Finally, genome-scale experiments could help address fundamental biological questions, such as what fraction of genetic changes elicit global transcriptional phenotypes and how transcriptional programs are rewired between cell types, with implications for understanding the organizing principles of cellular systems (*19*).

Here we perform the first genome-scale Perturb-seq screens. We use a compact, multiplexed CRISPR interference (CRISPRi) library to assay thousands of loss-of-function genetic perturbations with single-cell RNA-sequencing (scRNA-seq) in chronic myeloid leukemia (K562) and retinal pigment epithelial (RPE1) cell lines. Leveraging the scale and diversity of these perturbations across >2.5 million cells, we show that Perturb-seq can be used to study numerous complex cellular phenotypes—from RNA splicing to differentiation to chromosomal instability— in a single screen. We demonstrate how the interpretability of scRNA-seq phenotypes enables the discovery of gene function and extensively validate our findings with orthogonal experiments. Finally, we invert our analysis to focus on regulatory networks rather than genetic perturbations and uncover unanticipated stress-specific regulation of the mitochondrial genome. In sum, we use Perturb-seq to reveal a multidimensional portrait of cellular behavior, gene function, and regulatory networks that advances the goal of creating comprehensive genotype-phenotype maps.

## Results

### A multiplexed CRISPRi strategy for genome-scale Perturb-seq

Perturb-seq uses scRNA-seq to concurrently read out the CRISPR single-guide RNAs (sgRNA) (i.e., genetic perturbation) and transcriptome (i.e., high-dimensional phenotype) of single cells in a pooled format (Fig. 1A). To enable genome-scale Perturb-seq, we considered key parameters that would increase scalability and data quality, such as the genetic perturbation modality and sgRNA library.

**Figure 1:**
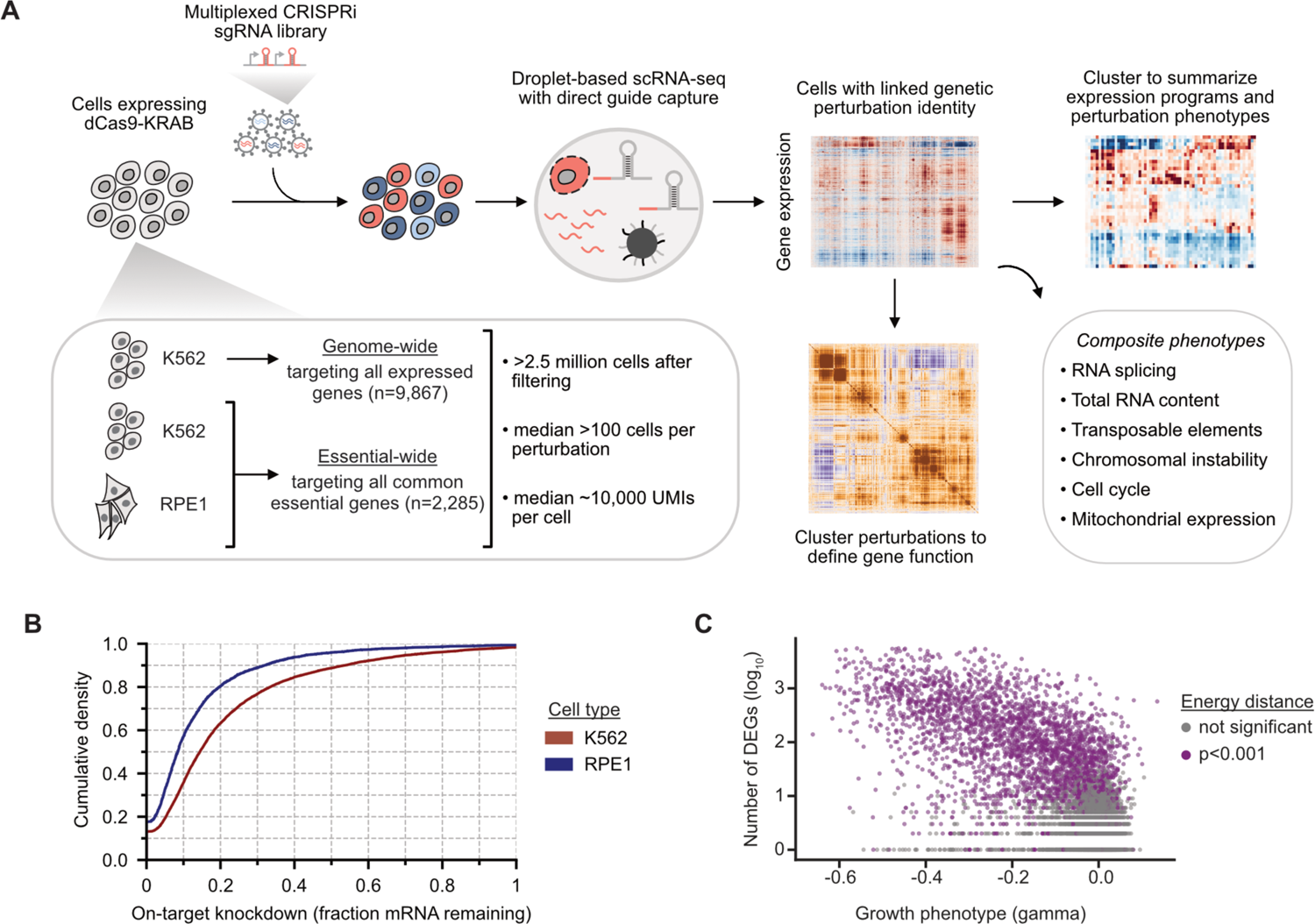
Genome-scale Perturb-seq via multiplexed CRISPRi. **(A)** Schematic experimental strategy. A multiplexed CRISPRi sgRNA library was used to knock down all expressed genes (in K562 cells) or all common essential genes (in RPE1 and K562 cells). Cells were transcriptionally profiled using droplet-based single-cell RNA-sequencing, with genetic perturbations assigned to cells by direct capture and sequencing of sgRNAs. **(B)** On-target knockdown statistics. Cumulative density plot of on-target knockdown, for n=9,464 target genes in K562 cells (red) and n=2,333 target genes in RPE1 cells (blue). **(C)** Comparing growth phenotype versus the number of differentially expressed genes (DEGs) for each multiplexed guide pair in K562 cells. Growth phenotypes are reported as the log_2_ guide enrichment per cell doubling (gamma). DEGs were determined using a two-sample Anderson-Darling test compared against non-targeting guides, and a pseudocount of 1 was added to the number of DEGs before log_10_ transformation. Dots are colored by Energy distance as either permutation significant (purple) or not significant (grey). The growth phenotype and number of DEGs are anticorrelated (Spearman’s rho=-0.51).

Although Perturb-seq is compatible with a range of CRISPR-based perturbations including knockout (*10–12*), knockdown (CRISPRi) (*9*), or activation (CRISPRa) (*20*), we elected to use CRISPRi for several reasons. First, CRISPRi allows the efficacy of the genetic perturbation, knockdown, to be directly measured from scRNA-seq. Exploiting this feature allowed us to target each gene in our library with a single element and empirically exclude unperturbed genes from downstream analysis. Second, CRISPRi tends to yield more homogeneous genetic perturbation than nuclease-based CRISPR knockout, which can generate a subset of cells bearing active in-frame indels (*21*). The relative homogeneity of CRISPRi limits selection for unperturbed cells, especially when studying essential genes. Third, unlike nuclease-based gene knockout, CRISPRi does not lead to activation of the DNA damage response which can alter cell state and transcriptional signatures (*22*).

To improve scalability, we optimized our CRISPRi sgRNA libraries. To maximize CRISPRi efficacy, we used multiplexed CRISPRi libraries in which each construct contains two distinct sgRNAs targeting the same gene (table S1-S3; see *Methods*) (*13*). To avoid low representation of sgRNAs targeting essential genes, we performed preliminary growth screens and, during library synthesis, overrepresented constructs that caused strong growth defects (fig. S1A-D).

Next, we devised a three-pronged Perturb-seq screening approach encompassing multiple timepoints and cell types (Fig. 1A). As a primary cell line, we studied chronic myeloid leukemia (CML) K562 cells engineered to express the CRISPRi effector protein dCas9-KRAB (*23*). In this cell line, we performed two Perturb-seq screens: one targeting all expressed genes sampled at day 8 after transduction (n=9,866 genes; n=10,673 total perturbations; some genes have multiple independent transcripts) and another targeting common essential genes, which was sampled at day 6 after transduction (n=2,057 genes; n=2,176 total perturbations). As a secondary cell line, we used RPE1 cells engineered to express dCas9 fused to a KRAB domain derived from the gene *ZIM3*, which was recently shown to yield improved transcriptional repression compared to the *KOX1*-derived KRAB domain used in previous CRISPRi experiments (*24*). In contrast to K562 cells, RPE1 cells are a non-cancerous retinal pigment epithelial cell line that are hTERT-immortalized, near-euploid, adherent, and p53-positive. In RPE1 cells, we performed a screen targeting common essential genes plus a subset of nonessential genes that produced phenotypes in K562 cells sampled at day 7 after transduction (n=2,393 genes; n=2,549 total perturbations).

We conducted these three screens with 10x Genomics droplet-based 3’ scRNA-seq and direct sgRNA capture (*13*). After sequencing and read alignment, we performed sgRNA identification and removed any cells bearing sgRNAs targeting different genes, which are an expected byproduct of lentiviral recombination between sgRNA cassette or doublet encapsulation during scRNA-seq. In total, we obtained >2.5 million high-quality cells with a median coverage of >100 cells per perturbation (fig. S1 E-G; table S4-S6). We observed a median target knockdown of 85.5% in K562 cells and 91.6% in RPE1 cells (Fig. 1B), confirming both the efficacy of our CRISPRi libraries and the fidelity of sgRNA assignment (*13*). The difference in performance between these cell lines was likely due to the use of the optimized *ZIM3*-derived KRAB domain in the RPE1 cells, suggesting that future efforts would benefit from improved CRISPRi efficacy.

### A robust computational framework to detect transcriptional phenotypes

The scale of our experiment provided a unique opportunity to ask what fraction of genetic perturbations cause a transcriptional phenotype, a preliminary requirement for inferring gene function. Significant transcriptional phenotypes can take many forms, ranging from altered occupancy of a given cell state to focused changes in the expression level of a small number of target genes. To contend with this diversity, we created a robust framework capable of detecting transcriptional changes between groups of cells in our data. Our experimental design included many control cells bearing diverse non-targeting sgRNAs. These allow for internal *z*-normalization of expression measurements, and we found that this procedure corrected for batch effects that resulted from parallelized scRNA-seq and sequencing (fig. S2). As Perturb-seq captures single-cell genetic perturbation identities in a pooled format, we can use statistical approaches that treat each cell as an independent experimental sample. In general, we chose to use conservative, non-parametric statistical tests to detect transcriptional changes rather than making specific assumptions about the underlying distribution of gene expression levels.

First, we examined global transcriptional changes using a permuted energy distance test (see *Methods*). We compared cells bearing each genetic perturbation to non-targeting control cells at the level of principal components (approximating global transcriptional features like cell state and gene expression programs). Relative to a permuted null distribution, this test asks whether cells carrying a given genetic perturbation could have been drawn from the control population. By this metric, we found that 2,987 of 9,608 genetic perturbations targeting a primary transcript (31.1%) compared to 11 of 585 non-targeting controls (1.9%) caused a significant transcriptional phenotype in K562 cells.

While sensitive, the energy distance test assays global shifts in expression without providing insight into which specific transcripts are altered. To detect individual differentially expressed genes, we applied the Anderson-Darling (AD) test to compare the distribution of expression levels for each gene in cells bearing each genetic perturbation against control cells. Importantly, the AD test is sensitive to transcriptional changes in a subset of cells, enabling us to find differences even when phenotypes have incomplete penetrance. With the AD test, we found 2,935 of 9,608 genetic perturbations targeting a primary transcript (30.5%) compared to 12 of 585 non-targeting controls (2.1%) caused >10 differentially expressed genes in K562 cells. These results were well-correlated between time points and cell types (fig. S3A,B; tables S4-S6) and concordant with the energy distance test (78.7% concordance by Jaccard index).

We then explored features of genetic perturbations that predict a transcriptional phenotype. We found that the strength of the transcriptional response was correlated with the strength of the growth defect (Spearman’s rho = –0.51) with 86.6% of essential genetic perturbations (gamma < –0.1) leading to a significant transcriptional response in K562 cells (Fig. 1C; fig. S3C,D). A substantial number of genetic perturbations that cause a transcriptional phenotype nonetheless have a negligible growth phenotype (n=771; fig. S3E), indicating that many genetic perturbations influence cell state but not growth or survival. We also found that highly expressed genes were more likely to produce transcriptional phenotypes (Spearman’s rho = 0.42) (fig. S3C).

Considering that some of our genetic perturbations did not yield strong on-target knockdown, our estimate of the fraction of genetic perturbations that cause a transcriptional phenotype is likely to be a lower bound. While a fraction of phenotypes may result from off-target effects, an advantage of Perturb-seq is the ability to directly detect potential off-target activities such as the knockdown of neighboring genes. Consistent with earlier studies (*25*), we found that ∼7.5% of perturbations caused significant knockdown of a neighboring gene in K562 cells, but neighbor gene knockdown was not enriched in genetic perturbations with a negligible growth defect that produced a transcriptional phenotype (fig. S4). Taken together, these results present a coherent picture where knockdown of a significant fraction of expressed genes causes a transcriptional response.

### Annotating gene function from transcriptional phenotypes

Previous Perturb-seq screens have focused on targeted sets of genetic perturbations that are often related biologically, such as genes identified in forward genetic screens. Our large-scale screen targeting all expressed genes in K562 cells presented a unique opportunity to assess how well transcriptional phenotypes can resolve gene function when used in an unbiased manner.

We focused on a subset of 1,973 perturbations that had strong transcriptional phenotypes (>50 differentially expressed genes by AD test) (Fig. 2A). Because related perturbations could have different magnitudes of effect, we used the correlation between mean expression profiles as a scale-invariant metric of similarity.

**Figure 2:**
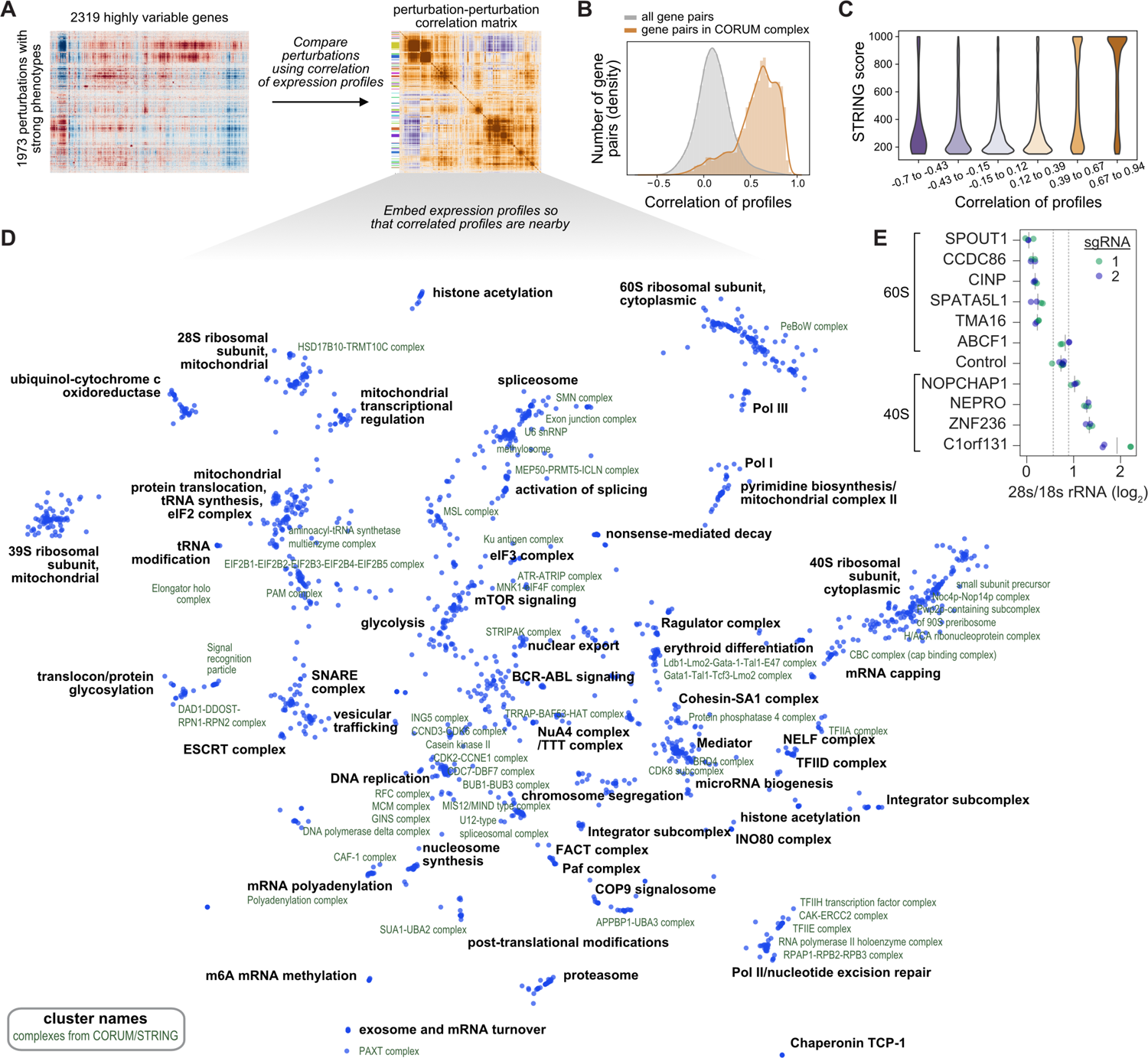
Data-driven inference of gene function from transcriptional phenotypes. **(A)** Schematic of analysis. To examine the ability of transcriptional phenotypes to assign gene function, we analyzed 1973 genetic perturbations that elicited strong responses. Perturbations were compared and clustered using the correlation of gene expression across 2319 highly variable genes. **(B)** Expression profile correlations among genes in curated complexes. 327 protein complexes from the CORUM3.0 database have at least two thirds of complex subunits within the dataset. Plot compares the distribution of pairwise expression profile correlations among genes in complexes vs. all possible gene-gene pairs. **(C)** Comparing expression profile correlations to predicted protein-protein interactions from STRING. 243,558 gene-gene relationships within the dataset are scored within STRING. The relationships were sorted into 6 equally spaced bins based on expression profile correlation. Plot shows kernel density estimates of STRING scores within each bin. **(D)** Minimum distortion embedding of dataset. Each dot represents a genetic perturbation, arranged so that perturbations with correlated expression profiles are nearby in the two dimensional embedding. Manual annotations (black labels) of cluster function are placed near the median location of genes within the cluster. CORUM complexes or STRING clusters (green labels) are annotated when involved genes are nearby within the embedding. **(E)** Quantification of 28s to 18s rRNA ratio. Poorly characterized genes with Perturb-seq predicted roles in ribosome biogenesis were targeted by CRISPRi. The 28s/18s rRNA ratio was measured by Bioanalyzer electrophoresis in biological duplicate with two distinct sgRNAs per gene (green and blue; solid grey lines represent mean). Dotted grey lines represent two standard deviations above and below the mean of non-targeting controls.

To assess the extent to which correlated mean expression profiles between genetic perturbations indicated common function, we compared our results to two curated sources of biological relationships. First, among the 1,973 targeted genes, there were 327 protein complexes from the CORUM 3.0 database with at least two thirds of the complex members present, representing 14,165 confirmed protein-protein interactions (*26*). The corresponding expression profile correlations were markedly stronger (median correlation 0.61) than the background distribution of all possible gene pairs (median correlation 0.10) (Fig. 2B). Second, we compared the correlation between genetic perturbations to the STRING database of known and predicted protein-protein interactions, which had scores for 243,558 of the possible gene-gene relationships within our dataset (*27*). High STRING scores, reflecting high-confidence interacting proteins, were also strongly associated with high expression correlations (Fig. 2C).

We next performed an unbiased search for global structure to group similar perturbations within the dataset. We identified 64 discrete clusters based on strong intra-cluster correlations and annotated their function using CORUM, STRING, and manual searches. To visualize the dataset, we constructed a minimum distortion embedding that places genes with correlated expression profiles close to each other in the plane and labeled the location of gene clusters (Fig. 2D).

Both the clusters and the embedding showed clear organization by biological function spanning a diverse array of different processes including: chromatin modification; transcription; mRNA splicing, capping, polyadenylation, and turnover; nonsense-mediated decay; translation; post-translational modification, trafficking, and degradation of proteins; central metabolism; mitochondrial transcription and translation; DNA replication; cell division; microRNA biogenesis; and major signaling pathways active in K562 cells such as BCR-ABL and mTOR (table S7). We further annotated the embedding visualization by labeling CORUM complexes and STRING clusters whose members were placed in nearby positions, revealing structure at finer resolution such as identifying the SMN complex, exon junction complex, U6 snRNP, and methylosome within the spliceosome and the association of ribosome biogenesis factors with the 40S ribosomal subunit.

In our dataset, we identified many poorly annotated genes whose perturbation led to similar transcriptional responses to genes of known function, naturally predicting a role for the uncharacterized genes. To orthogonally test a subset of these predictions, we selected ten poorly annotated genes whose perturbation response correlated (*r*>0.6) with subunits and biogenesis factors of either the large or small subunit of the cytosolic ribosome, which formed distinct clusters in our data (fig. S5A). This included genes that had no previous association with ribosome biogenesis (*CCDC86*, *CINP*, *SPATA5L1*, *ZNF236*, *C1orf131*) as well as genes that had not been associated with functional defects in a particular subunit (*SPOUT1*, *TMA16*, *NOPCHAP1*, *ABCF1*, and *NEPRO*). We used CRISPRi to target these genes in K562 cells and looked for evidence of ribosome biogenesis defects by assessing the ratio of 28S to 18S rRNA by Bioanalyzer electrophoresis. Knockdown of nine of the ten candidate factors led to substantial defects in ribosome biogenesis, with the exception of *ABCF1* (Fig. 2E). In every case, the affected ribosomal subunit corresponded to the Perturb-seq clustering across two independent sgRNAs. While this study was in progress, another group used cryo-EM to identify C1orf131 as a core structural component of the pre-A1 small subunit processome, complementing our functional evidence (*28*). This validation suggests that many poorly characterized genes can be assigned functional roles through Perturb-seq, although a subset of these relationships might be explained by indirect or off-target effects (fig. S5B,C).

In total, these results show that transcriptional phenotypes revealed by Perturb-seq have utility beyond studying gene regulation or transcriptional programs, and can serve as valuable tools for resolving and interrogating many central processes in cell biology.

### Delineating functional modules of the Integrator complex

In general, perturbations to members of known protein complexes produced similar transcriptional phenotypes in our dataset. Therefore, we were surprised by the wide spectrum of transcriptional responses to knockdown of subunits of Integrator, a metazoan-specific essential nuclear complex with roles in small nuclear RNA (snRNA) biogenesis and transcription termination at paused RNA polymerase II (Fig. 3A) (*29*). Each of the fourteen core subunits of Integrator was targeted in our experiment, allowing us to systematically compare their transcriptional phenotypes in K562 and RPE1 cells (Fig. 3B; fig. S6A). *INTS1*, *INTS2*, *INTS5*, *INTS7*, and *INTS8* formed a tight cluster which weakly correlated with *INTS6* and *INTS12*. Separately, *INTS3*, *INTS4*, *INTS9*, and *INTS11* clustered together alongside splicing regulators involved in snRNP assembly and the tri-snRNP. Finally, *INTS10*, *INTS13*, and *INTS14* formed another discrete cluster together with *C7orf26*, an uncharacterized gene.

**Figure 3:**
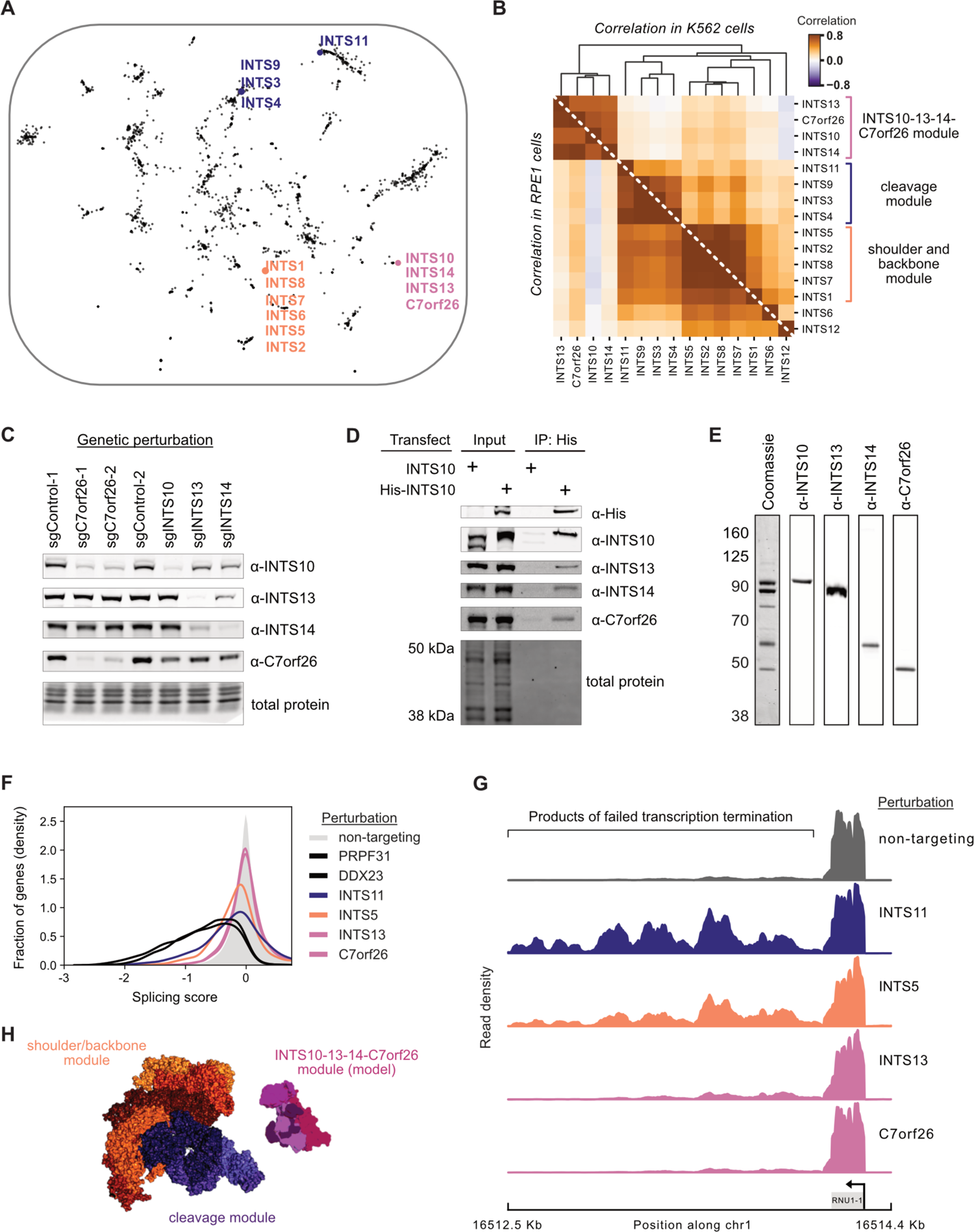
Perturb-seq discovers a novel gene member and functional submodules of the Integrator complex. **(A)** Location of known Integrator complex members in the minimum distortion embedding. **(B)** Relationship between Integrator complex members and *C7orf26* in K562 cells (top) and RPE1 cells (bottom). The heatmap displays the Pearson correlation between pseudobulk *z*-normalized gene expression profiles of Integrator complex members. Genetic perturbations are ordered by average linkage hierarchical clustering based on correlation in K562 cells. Functional modules suggested by the clustering are highlighted. **(C)** Co-depletion of Integrator complex members. Individual Integrator complex members were depleted in CRISPRi K562 cells. Lysates were then probed for other module members by western blot. **(D)** Co-immunoprecipitation of endogenous C7orf26 with His-INTS10. HEK293T were transfected with His-INTS10 or INTS10. Cell lysates were affinity purified and select Integrator proteins were probed by western blot. **(E)** Purification of a INTS10-INTS13-INTS14-C7orf26 complex. His-INTS10, INTS13, INTS14, and C7orf26 were overexpressed in Expi293 cells, affinity purified, and separated via SEC. The INTS10-INTS13-INTS14-C7orf26 proteins co-fractionated as visualized by Western blotting. **(F)** Effects of Integrator modules on splicing from Perturb-seq data. Histogram (kernel density estimate) compares gene-level splicing scores. Splicing scores represent the change in the log_2_ ratio of total to unspliced reads for each gene relative to non-targeting control guides. Representative genetic perturbations from Integrator modules as well as the spliceosome are shown colored by module. **(G)** Density of PRO-seq reads at the snRNA *RNU1-1* locus mapping actively engaged RNA polymerase II. For each perturbation, densities are shown relative to the maximum read count in the locus. **(H)** Structure of the Integrator complex colored by functional modules revealed by Perturb-seq. The endonuclease (blue) and shoulder/backbone (orange) modules were obtained from the cryo-EM structure (*31*). The model of the newly discovered 10-13-14-C7orf26 module was built by docking the crystal structure of INTS13-INTS14 (*33*) with an AlphaFold multimeric model of INTS10 and C7orf26.

These distinct functional modules mirror the architecture of the Integrator complex observed in recent structures (*30, 31*). The INTS1-2-5-7-8 functional module contained the subunits identified as the structural shoulder and backbone of Integrator. The INTS3-4-9-11 functional module contained the subunits identified as the structural cleavage module (as well as INTS3 which was not resolved). While INTS10, INTS13, and INTS14 were not resolved in the recent cryo-EM Integrator structures, these subunits have been identified as a stable biochemical subcomplex (*32, 33*).

Integrator is an essential and well-studied complex, so we were intrigued by the robust clustering of the uncharacterized gene *C7orf26* with Integrator subunits 10, 13, and 14. To explore this, we tested whether loss of C7orf26 impacted the abundance of Integrator subunits. CRISPRi-based depletion of C7orf26 destabilized INTS10 in K562 cells, confirming either a regulatory or protein-level relationship (Fig. 3C). Next, we checked for a biochemical interaction between these proteins. Pulldown of His-INTS10 from cell lysates recovered endogenous C7orf26 alongside INTS13 and INTS14 (Fig. 3D). Additionally, overexpression of C7orf26 with INTS10, INTS13, and INTS14 enabled the purification of a stable INTS10-13-14-C7orf26 complex by size-exclusion chromatography (Fig. 3E; fig. S6B and fig. S7). We also detected a physical interaction between the Drosophila C7orf26 orthologue and fly Integrator in S2 cells and observed co-essentiality between C7orf26 and INTS10, INTS13, INTS14 in the Cancer Dependency Map, suggesting that this relationship is conserved across species and cell types (fig. S8). Together, these results suggest that C7orf26 is a core subunit of a novel INTS10-13-14-C7orf26 Integrator module. We sought to better understand the distinct transcriptional phenotypes induced by knockdown of INTS10-13-14-C7orf26 compared to the shoulder/backbone and cleavage modules. Comparison of the genes differentially expressed between modules did not reveal function in an obvious way (fig. S6C,D), perhaps owing to the late time point assayed in our experiment. We next explored the canonical role of Integrator in snRNA biogenesis. As mature snRNAs are not captured in 3’ scRNAs-seq, we monitored changes in global splicing as a proxy for snRNA biogenesis defects. In our Perturb-seq data, we quantified changes in splicing by comparing the ratio of intronic (unspliced) to exonic (spliced) reads for each gene. Validating our approach, depletion of known splicing factors as well as subunits of the cleavage and shoulder/backbone modules led to gross splicing defects (Fig. 3F). By contrast, depletion of subunits of the INTS10-13-14-C7orf26 module did not cause a substantial splicing defect. To directly test the effect of the INTS10-13-14-C7orf26 module on snRNA biogenesis, we used PRO-seq to probe the positioning of active RNA-polymerase. These data confirmed that extended knockdown of the cleavage and backbone/shoulder modules, but not *INTS10*, *INTS13*, or *C7orf26*, caused a dramatic increase in transcriptional readthrough past the 3’ cleavage site of snRNAs (Fig. 3G). In addition, the PRO-seq data confirmed that loss of the INTS10-13-14-C7orf26 module causes a transcriptional phenotype distinct from other modules (fig. S6E).

In sum, our results show that INTS10-13-14-C7orf26 represents a functionally and biochemically distinct module of the Integrator complex, and we propose that *C7orf26* be renamed *INTS15* for future studies (Fig. 3H). Although Integrator has been subjected to extensive structural analyses, it has been difficult to resolve the INTS10-13-14 components in relation to the rest of the complex. Inclusion of C7orf26 may facilitate future structural efforts. Broadly, this example highlights the utility of high-dimensional functional phenotypes for the unsupervised classification of protein complex subunits into functional modules.

### Data-driven definition of transcriptional programs

While clustering can organize genetic perturbations into pathways or complexes, it does not reveal the functional consequences of perturbations. To globally summarize the genotype-phenotype relationships in our dataset, we: (i) clustered genes into expression programs based on their co-regulation across perturbations; (ii) clustered perturbations with strong phenotypes based on their transcriptional profiles (as described above); and (iii) computed the average activity of each gene expression program within each perturbation cluster (Fig. 4A,B; fig. S9A; table S7,S8; see *Methods*). This map uncovered many known gene expression programs associated with genetic perturbations, including upregulation of proteasomal subunits due to proteosome dysfunction (*34*), activation of NFkB signaling upon loss of ESCRT proteins (*35*), downregulation of growth-related genes in response to essential genetic perturbations, and upregulation of the cholesterol biosynthesis pathway in response to defects in vesicular trafficking (*36*). Beyond these large-scale relationships, we could also score the effects of individual genetic perturbations on different expression programs. For example, our analysis delineated the canonical branches of the cellular stress response into the independently regulated Unfolded Protein Response (UPR) and Integrated Stress Response (ISR) (Fig. 4C) (*9*). The ISR was highly activated by loss of mitochondrial proteins, aminoacyl-tRNA synthetases, and translation initiation factors, whereas the UPR was activated by loss of ER-resident chaperones and translocation machinery. Collectively, this analysis establishes the ability of Perturb-seq to learn regulatory circuits by leveraging the variability of responses across perturbations.

**Figure 4:**
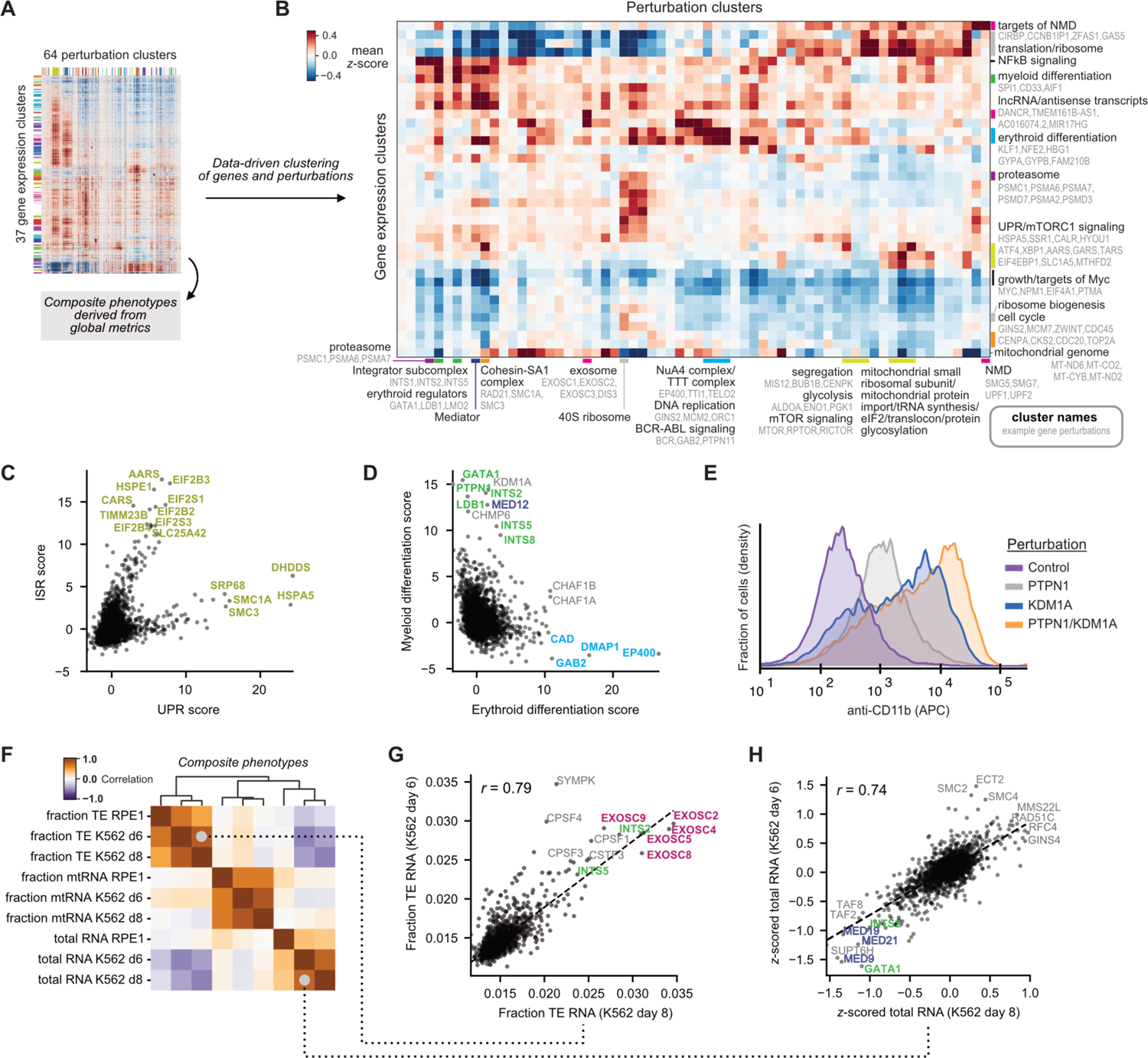
Summarizing genotype-phenotype relationships with Perturb-seq. **(A)** Schematic of analysis. To produce a high-level summary of genotype-phenotype relationships in K562 cells, 1973 genetic perturbations that elicited strong responses and 2319 highly variable genes were clustered using HDBSCAN after nonlinear embedding. Alternatively, composite phenotypes were derived from global metrics in a hypothesis-driven manner. **(B)** Heatmap of the high-level genotype-phenotype map. The heatmap represents the mean z-scored expression for gene expression and perturbation clusters. For a subset of clusters, clustered are labelled with manual annotations (black labels) of cluster function along with example genes within the cluster (light gray labels). **(C)** Comparison of ISR and UPR scores for genetic perturbations. Scores were recovered from unbiased clustering of genes by genetic dependency, and manually annotated. **(D)** Comparison of erythroid and myeloid differentiation scores for genetic perturbations. Scores were recovered from unbiased clustering of genes by genetic dependency, and manually annotated. Genetic perturbations are colored to reflect cluster identity. **(E)** Expression of CD11b/ITGAM in K562 cells upon knockdown of *PTPN1* or *KDM1A*. CD11b was labelled by cell surface staining with anti-CD11b antibody and measured by flow cytometry. **(F)** Correlation of composite phenotypes across time points and cell types. Composite phenotypes were defined in a hypothesis-driven manner. Fraction TE (repetitive and transposable element) represents the number of non-intronic reads mapped to TEs over total, averaged over all cells bearing each perturbation (both collapsed on UMIs). Fraction mtRNA represents the mean number of reads mapped to mitochondrial genome protein-coding genes over total. Total RNA represents the mean total RNA content (number of UMIs). **(G)** Comparison of TE expression across time points. The mean fraction TE reads per perturbation is highly correlated across time points in K562 cells (*r*=0.79). Genetic perturbations are colored to reflect cluster identity. **(H)** Comparison of total RNA content across time points. Total RNA content per perturbation is highly correlated across time points in K562 cells (*r*=0.74). Genetic perturbations are colored to reflect cluster identity.

Interestingly, our unbiased clustering uncovered many perturbations that drove the expression of markers of erythroid or myeloid differentiation, consistent with the known multilineage potential of K562 cells (Fig. 4D) (*37*). The scale of our experiment allowed us to comprehensively search for genes whose modulation promotes cellular differentiation, an application of major interest in both developmental and cancer biology. As expected, loss of central regulators of erythropoiesis (*GATA1*, *LDB1*, *LMO2*, and *KDM1A*) caused myeloid differentiation, whereas knockdown of *BCR-ABL* and its downstream adaptor *GAB2* induced erythroid differentiation (*38*). Surprisingly, loss of a number of common essential genes (i.e., essential across cell lines in the Cancer Dependency Map) also caused expression of either myeloid (e.g., Integrator subcomplex) or erythroid (e.g., NuRD complex, DNA replication machinery) markers. Next, we investigated the differentiation effect of selectively essential genes, which could be promising targets for differentiation therapy, analogous to ongoing efforts for KDM1A (*39, 40*). We observed that loss of *PTPN1*, a tyrosine phosphatase selectively essential in K562 cells, drove myeloid differentiation. While inhibitors of PTPN1 have been developed for use in diabetes and certain cancers (*41*), to our knowledge they have not been tested as a differentiation therapy. Remarkably, in targeted experiments, we found that knockdown of *PTPN1* and *KDM1A* in combination caused a substantial increase in differentiation and growth defect compared to either single genetic perturbation, suggesting that these targets act via different cellular mechanisms (Fig. 4E; fig. S9B). These results highlight the utility of rich phenotypes for understanding differentiation as well as nominating promising therapeutic targets or combinations.

### Hypothesis-driven study of composite phenotypes

We next recognized that our scRNA-seq readout could be used to study phenotypes that integrate data from across the transcriptome and, therefore, would be difficult to study in traditional forward genetic screens. Examples of these “composite phenotypes” include total cellular RNA content and the fraction of RNA derived from transposable elements (TE). We found numerous composite phenotypes under strong genetic control, with highly reproducible effects across screen replicates and cell types (Fig. 4F). In the specific case of TE regulation, two major classes of perturbations increased the fraction of TE RNA by affecting broad classes of elements including Alu, L1, and MIR (Fig. 4G; fig. S9C). First, loss of subunits of the exosome led to a substantial increase in the fraction of TE RNA, suggesting that transcripts deriving from TEs might be preferentially degraded. Second, loss of the CPSF cleavage and polyadenylation complex and parts of the Integrator complex produced a similar phenotype, suggesting that many of the TE RNAs observed in K562 cells may be derived from failure of normal transcription termination.

Turning to total RNA content (Fig. 4H), we found that loss of many essential regulators of S-phase and mitosis increased the RNA content of cells. This is consistent with the observation that cells tend to increase their size, and thus their RNA content, as they progress through the cell cycle (fig. S9D), so perturbations that arrest cells in later cell cycle stages yield increased total RNA content on average. By contrast, loss of essential transcriptional machinery, including general transcription factors, the Mediator complex, and transcription elongation factors, decreased total RNA content. In sum, these analyses show that genome-scale Perturb-seq enables hypothesis-driven exploration of complex cellular features that are challenging to study through other means.

### Exploring genetic drivers and consequences of aneuploidy in single-cells

As Perturb-seq is a single-cell assay, it enables the study of cell-to-cell heterogeneity in response to genetic perturbations. We reasoned that systematically exploring sources of heterogeneity could reveal insights into phenotypes that are missed in bulk or averaged measurements.

To assess the penetrance of perturbation-induced phenotypes, we first applied SVD-based leverage scores as a metric of single-cell phenotypic magnitude (see *Methods*). In this formulation, leverage scores quantify how outlying each perturbed cell’s transcriptome is relative to non-targeting control cells without assuming that perturbations drive a single axis of variation. Supporting this approach, we found that mean leverage scores for each genetic perturbation were correlated with the number of differentially expressed genes (fig. S10A, Spearman’s rho = 0.71), and reproducible across the day 6 and day 8 K562 experiments (fig. S10B, Spearman’s rho = 0.79). To quantify the degree of heterogeneity in response to genetic perturbations, we then scored perturbations by the variation in single-cell leverage scores (Fig. 5A; see *Methods*). Comparing leverage scores across subunits of large essential complexes, we observed evidence for both biological (e.g., subcomplex function or dosage imbalance) and technical (e.g., selection to escape toxic perturbations) sources of phenotypic variation in response to genetic perturbations (fig. S10C-F).

**Figure 5:**
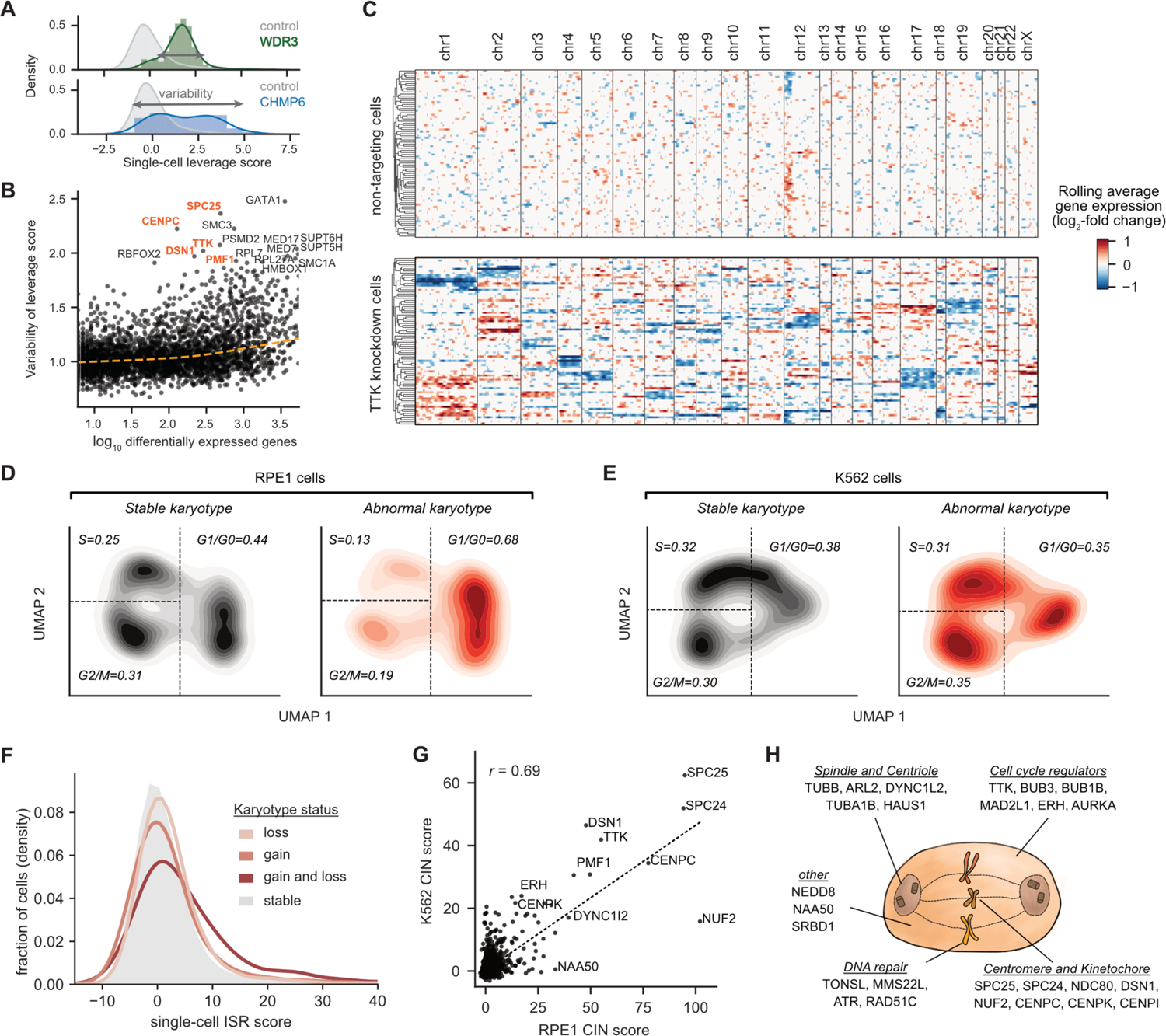
Exploring acute consequences and genetic drivers of aneuploidy in single-cells. **(A)** Schematic of heterogeneity statistic. Single-cell leverage scores quantify how outlying each cell is relative to non-targeting control cells by PCA. For each perturbation, heterogeneity of single-cell phenotypes is quantified as the standard deviation of leverage scores. **(B)** Identifying heterogeneous perturbations. Known regulators of chromosome segregation were among the perturbations with the highest single-cell heterogeneity (high variability of leverage scores), especially compared to their number of differentially expressed genes (based on Anderson-Darling test). **(C)** Heatmap of chromosomal copy number inference from Perturb-seq data. For all genes (expressed >0.05 UMI per cell), the log-fold change in expression is calculated with respect to the average of non-targeting control cells, and genes are ordered along the genome. A weighted moving average of 100 genes is used infer copy number changes (columns) in single-cells (rows) with noise and median filtering. 80 *TTK* knockdown RPE1 cells and 80 randomly sampled non-targeting control RPE1 cells are shown. Cells are ordered by average linkage hierarchical clustering based on correlation of chromosomal copy number profiles. **(D, E)** Comparison of cell cycle occupancy upon acute karyotypic changes. Abnormal karyotypic cells were defined as having ≥1 chromosome with evidence of changes in chromosomal copy number for >80% of the chromosomal length. For single-cells, cell-cycle positioning was inferred by UMAP dimension reduction on differential expression profiles of 199 selected cell-cycle regulated genes. Cell cycle occupancy is shown as a 2D kernel density estimate of a random subset of 1000 cells per karyotypic status. Approximate gates between cell cycle phases (G1 or G0; S; G2 or M) are shown as dotted lines, and the fraction of cells in each cell cycle phase are indicated. **(F)** Effect of chromosomal instability (CIN) on activation of the Integrated Stress Response (ISR). Histogram (kernel density estimate) compares the ISR score versus CIN status in RPE1 cells. CIN status is defined as evidence of gain or loss of chromosomal copy number for >80% of the chromosomal length, with 240,768 stable cells, 5,522 cells bearing chromosomal loss, 1987 cells bearing chromosomal gain, and 904 cells bearing gain and loss of chromosomes. ISR score is defined as the sum of z-normalized expression of ISR marker genes where increased values indicate stronger ISR activation. **(G)** Comparison of the effect of genetic perturbations on the CIN score across cell types. For each genetic perturbation, the CIN score is calculated as the mean single-cell sum of squared CIN values, *z*-normalized relative to non-targeting control perturbations. The CIN score is correlated across cell types (*r*=0.69). **(H)** Schematic of a subset of genetic perturbations that drive CIN. CIN drivers play diverse roles in mitosis, cell cycle regulation, and DNA repair.

Intriguingly, many genes implicated in chromosome segregation were among the top drivers of heterogeneity, including *TTK*, *SPC25*, and *DSN1* (Fig. 5B) (*42*). We hypothesized that the extreme transcriptional variability caused by these genetic perturbations might result from acute changes in the copy number of individual chromosomes due to mitotic mis-segregation. To explore this, we used inferCNV (*43*) to estimate single-cell DNA copy number along the genome by quantifying the change in moving average gene expression compared to control cells. Consistent with our hypothesis, knockdown of *TTK*, a core component of the spindle assembly checkpoint (*44*), led to dramatic changes in estimated DNA copy number in both intrinsically aneuploid K562 and near euploid RPE1 cells (Fig. 5C; fig. S11A). Specifically, in RPE1 cells, we found that 61/80 (76%) of *TTK* knockdown cells had evidence of karyotypic changes compared to 274/13140 (2%) of unperturbed cells. Notably, *TTK* knockdown cells bore highly variable karyotypes due to the stochastic gain or loss of chromosomes, accounting for the phenotypic heterogeneity observed in these cells (Fig. 5C).

An important advantage of the rich data provided by Perturb-seq is the ability to dissect not only perturbation-phenotype associations but also relationships between cellular phenotypes. We were curious how chromosomal instability (CIN) would affect cell cycle progression in euploid, p53-positive RPE1 cells versus constitutively aneuploid, p53-deficient K562 cells. Expanding our analysis to all cells in our experiment independent of genetic perturbation, we found that RPE1 cells with abnormal karyotypes tended to arrest in G1 or G0 of the cell cycle (G1 or G0 fraction 0.68 for abnormal karyotype vs. 0.44 for stable karyotype), while K562 cells with altered karyotypes had less significant shifts in cell cycle occupancy (Fig. 5D,E). Within the population of RPE1 cells bearing a chromosomal loss, the likelihood of cell cycle arrest directly depended on the magnitude of karyotypic abnormality (fig. S11B). Additionally, we observed that cells with the most severe karyotypic changes—those bearing both chromosomal gains and losses—had marked upregulation of the ISR (Fig. 5F and fig. S11C). These results are consistent with models in which cell cycle checkpoints are activated by the secondary consequences of aneuploidy (e.g., DNA damage or proteostatic stress) rather than changes in chromosome number *per se* (*45, 46*).

Finally, we looked across all perturbations to systematically identify genetic drivers of CIN. We assigned a score to each perturbation based on the average magnitude of induced karyotypic abnormalities. Validating our approach, we found that CIN scores were strongly correlated across K562 and RPE1 cell lines (*r*=0.69) and identified many known regulators of chromosomal segregation, including components of the spindle assembly checkpoint, centromere, and NDC80 complex (Fig. 5G). Remarkably, we uncovered CIN regulators with diverse cellular roles, from cytoskeletal components to DNA repair machinery (Fig. 5H; table S4-S6). While many of these genes have previously been associated with chromosomal instability through targeted studies, the scale and single-cell resolution of Perturb-seq allowed us to identify numerous genetic drivers of CIN in a single experiment. This analysis also shows the potential of single-cell CRISPR screens to dissect phenotypes that were not predefined endpoints of the experiment.

### Discovery of stress-specific regulation of the mitochondrial genome

Mitochondria arose from the engulfment and endosymbiotic evolution of an ancestral alphaproteobacterium by the precursor to eukaryotic cells (*47*). While the vast majority (∼99%) of mitochondrially-localized proteins are encoded in the nuclear genome, mitochondria contain a small (∼16.6 kilobase) remnant of their ancestral genome encoding 2 rRNAs, 22 tRNAs, and 13 protein-coding genes in humans. An open question is how the nuclear and mitochondrial genomes coordinate their expression to cope with mitochondrial stress (*48*). The scale of our experiment provided a unique opportunity to investigate this question.

We began by comparing the nuclear transcriptional responses to CRISPRi-based depletion of nuclear-encoded mitochondrial genes (i.e., mitochondrial perturbations). We found that mitochondrial perturbations elicited relatively homogeneous nuclear transcriptional responses, illustrated by well-correlated transcriptional phenotypes across mitochondrial perturbations (Fig. 6A and fig. S12A). While there was some variation in the magnitude of transcriptional responses (e.g., proteostatic injury drove an especially strong ISR activation), nuclear transcriptional responses generally failed to discriminate genetic perturbations by function. Although this result was broadly consistent with recent literature that has highlighted the role of the ISR as response to mitochondrial stress (*49–53*), the lack of functional specificity of the transcriptional response was puzzling in light of: (i) the multifaceted roles of mitochondria in diverse processes such as respiration, intermediary metabolism, iron-sulfur cluster biogenesis, and apoptosis and (ii) the high-resolution separation of cytosolic perturbations by transcriptional response in our data described above.

**Figure 6:**
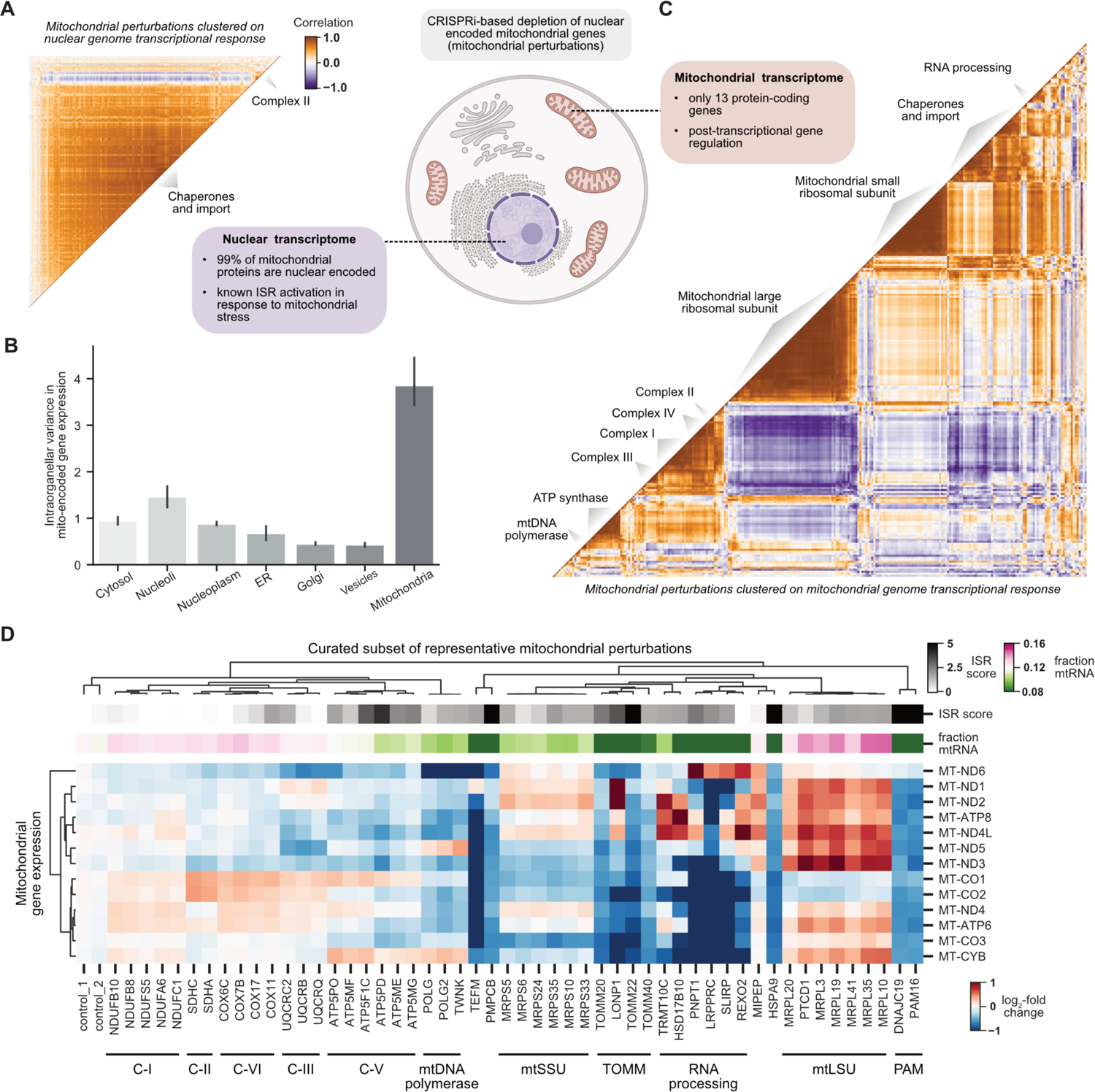
Global organization of the transcriptional response to mitochondrial stress. **(A)** Clustering mitochondrial perturbations by nuclear transcriptional response. CRISPRi enables knockdown of nuclear-encoded genes whose protein products are targeted to mitochondria (mitochondrial perturbations). Mitochondrial perturbations were annotated by MitoCarta3.0 and subset to those with a strong transcriptional phenotype (n=268 mitochondrial perturbations). Gene expression profiles were restricted to nuclear encoded genes which includes 99% of mitochondrial proteins. The heatmap displays the Pearson correlation between pseudobulk *z*-normalized gene expression profiles of mitochondrial perturbations in K562 cells. Genetic perturbations were clustered by HDBSCAN with a correlation metric. **(B)** Comparing variability in the mitochondrial transcriptome by perturbation localization. The mitochondrial genome encodes 13 protein-coding genes. Genetic perturbations were grouped based on localization of their protein products as determined by the Human Protein Atlas. For each of these 13 mitochondrially encoded genes, the variance in pseudobulk z-normalized expression profiles was calculated between all perturbations with the same localization. Barplots represent the average across genes with 95% confidence interval obtained by bootstrapping. **(C)** Clustering mitochondrial perturbations by mitochondrial transcriptional response. Mitochondrial perturbations were annotated by MitoCarta3.0 and subset to those with a strong transcriptional phenotype as above (n=268 mitochondrial perturbations). Gene expression profiles were restricted to the 13 mitochondrial-encoded genes. The heatmap displays the Pearson correlation between pseudobulk *z*-normalized gene expression profiles of mitochondrial perturbations in K562 cells. Genetic perturbations are clustered by HDBSCAN with a correlation metric. Clusters were manually annotated. **(D)** Heatmap visualizing the mitochondrial genome transcriptional response to diverse mitochondrial stressors. The expression (log_2_ fold-change relative to non-targeting controls) of the 13 mitochondrially encoded genes is shown for a subset of perturbations representative of different mitochondrial complexes or function. Neither the ISR score nor mean fraction of mitochondrial RNA (mtRNA) would allow for high-resolution clustering by function as provided by the mitochondrial genome response. Genetic perturbations and genes are ordered by average linkage hierarchical clustering with a correlation metric.

In contrast to the nuclear transcriptional response, we observed that the expression of mitochondrially encoded genes was highly variable between different mitochondrial perturbations (Fig. 6B; fig. S12B,C,D). When we clustered mitochondrial perturbations based solely on expression levels of the 13 mitochondrially encoded genes, a remarkably intricate and coherent pattern emerged: the clustering separated perturbations to Complex I, Complex IV, Complex III, Complex V (ATP synthase), the mitochondrial large ribosomal subunit, the mitochondrial small ribosomal subunit, chaperones/import machinery, and RNA processing factors (Fig. 6C; fig. S12E). To quantitatively support this observation, we trained a random forest classifier to distinguish cells with perturbations to different mitochondrial complexes and found that the mitochondrial transcriptome was far more predictive than the nuclear transcriptome (mitochondrial accuracy 0.64; nuclear accuracy: 0.25) (fig. S12F). We then visualized the gene expression signatures of a subset of representative perturbations (Fig. 6D). The coregulation of mitochondrial genes tended to reflect function, with the exception of the bicistronic mRNAs *ND4L/ND4* and *ATP8/ATP6* (*54*). However, we did not identify a simple logic to explain the connection between genetic perturbations to their observed transcriptional consequences. While previous studies have described distinct regulation of the mitochondrial genome in response to specific perturbations [notably, related to loss of complex III and complex IV assembly factors (*55, 56*)], our data generalize this phenomenon to a comprehensive set of stressors.

Next, we wanted to shed light on the mechanistic basis for this unappreciated complexity of mitochondrial genome responses. Given its singular origin, the mitochondrial genome is expressed by unique processes (Fig. 7A) (*57*). Mitochondrially encoded genes are transcribed as part of three polycistronic transcripts punctuated by tRNAs. These transcripts are then processed into rRNAs and mRNAs by tRNA excision, and individual mRNAs can be polyadenylated, expressed, or degraded. This complex system limits the potential for distinct transcriptional control but presents multiple opportunities for post-transcriptional regulation. To identify modes of perturbation-elicited differential expression, we examined the distribution of Perturb-seq reads along the mitochondrial genome (Fig. 7B). As our scRNA-seq used poly-A selection, most reads aligned to the 3’ ends of mRNAs. To validate the utility of this position-based analysis, we confirmed that knockdown of known regulators of mitochondrial transcription (*TEFM*) and RNA degradation (*PNPT1*) led to major shifts in the position of reads along the mitochondrial genome. By contrast, many of the perturbation-specific responses discovered in the present study appeared to cause shifts in the relative abundance of mRNAs rather than gross shifts in positional alignments. To determine whether the observed mitochondrial genome responses reflected regulation of the total level of mitochondrial mRNAs or specific regulation of mRNA polyadenylation, we performed a bulk RNA-sequencing experiment with no poly-A selection. We observed perturbation-specific changes in the level of total RNA similar to those measured by scRNA-seq (cophenetic correlation r=0.79; Fig. 7C). Given the complexity of the observed responses, we propose that there are likely to be multiple mechanisms that impact the levels of the various mitochondrially encoded transcripts in response to different stressors.

**Figure 7:**
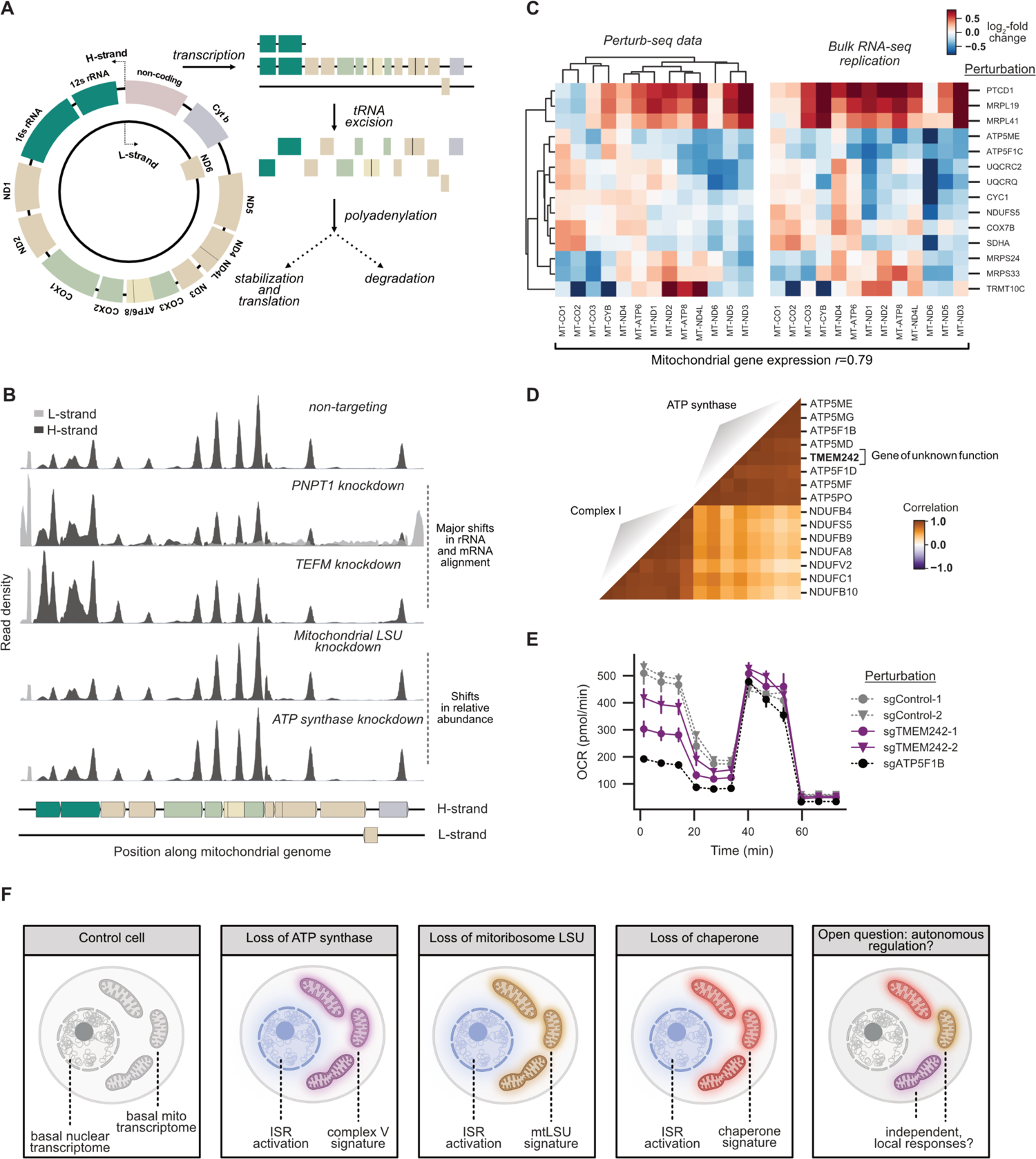
Investigating regulation of the mitochondrial genome in stress. **(A)** Schematic of the mitochondrial transcriptome. Each human cell contains many copies of the circular 16.6 kb mitochondrial genome distributed throughout the mitochondrial network. The human mitochondrial genome encodes 2 rRNAs, 22 tRNAs, and 13 protein-coding genes. Both the heavy (H) and light (L) strand of the genome are transcribed as polycistronic transcripts punctuated by tRNAs. Excision of tRNAs from transcripts generates nascent mRNA precursors (colored by complex membership). mRNA precursors can then be polyadenylated, stabilized, or degraded. **(B)** Density of Perturb-seq reads along the mitochondrial genome from select genetic perturbations. Reads are aligned to both the H-strand (dark grey) and L-strand (light grey). For each perturbation, densities are shown relative to the maximum read count in the locus. **(C)** Comparison of mitochondrial gene expression profiles between Perturb-seq and bulk RNA-seq. Heatmap displays log_2_-fold changes in expression of the 13 mitochondrial encoded genes (columns) for genetic perturbations (rows) in Perturb-seq and bulk RNA-seq data collected from K562 cells. Bulk RNA-seq was conducted to analyze total RNA (including non-polyadenylated RNA), with data representing the average of biological replicates. Genetic perturbations and genes are ordered by average linkage hierarchical clustering with a Euclidean distance metric. The profiles are strongly correlated (r=0.79, *p*<10^-39^). **(D)** Clustering of *TMEM242* genetic perturbation based on the mitochondrial transcriptome. Genetic perturbations to members of ATP synthase and Complex I of the respiratory chain were compared to knockdown of *TMEM242*, a mitochondrial gene of unknown function. Gene expression profiles were restricted to the 13 mitochondrially encoded genes. The heatmap displays the Pearson correlation between pseudobulk *z*-normalized gene expression profiles of mitochondrial perturbations in K562 cells. Genetic perturbations are ordered by HDBSCAN with a correlation metric. **(E)** Effect of *TMEM242* knockdown on mitochondrial respiration. A Seahorse analyzer was used to monitor oxygen consumption rate (OCR). The Mito Stress Test consists of sequential addition of oligomycin (an ATP synthase inhibitor that enables measurement of ATP-productive respiration), FCCP (an uncoupling agent that enables measurement of maximal respiratory capacity), and a mixture of rotenone and antimycin A (inhibitors of Complex I and Complex III, respectively, that enable measurement of non-mitochondrial respiration). Data is presented as average ± SEM, n=6. **(F)** Schematic diagram of mitochondrial stress response.

Finally, we asked whether we could use the detailed clustering produced by the mitochondrial genome to predict gene function. Knockdown of an unannotated gene, *TMEM242*, produced a transcriptional signature resembling loss of ATP synthase in both K562 and RPE1 cells (Fig. 7D; fig. S12G). Supporting this relationship, the top five co-essential genes with *TMEM242* were components of ATP synthase in the Cancer Dependency Map. Using a Seahorse assay, we further confirmed that basal respiration was decreased in *TMEM242* knockdown cells (Fig. 7E). While this work was in progress, another group used a biochemical approach to show that *TMEM242* physically interacts with ATP synthase subunits and regulates ATP synthase complex assembly in cells (*58*). Together, these experiments discover a novel factor required for ATP synthase activity and point to the precision of mitochondrial genome regulation.

## Discussion

Single-cell CRISPR screens represent an emerging tool to generate rich genotype-phenotype maps. However, to date, their use has been limited to the study of preselected genes focused on discrete, predefined biological questions. Here, we perform genome-scale single-cell CRISPR screens using Perturb-seq and demonstrate how these screens enable data-driven dissection of a breadth of complex biological phenomena. Reflecting on this study, we highlight key biological insights and derive principles to guide future discoveries from rich genotype-phenotype maps.

A primary aim of large-scale functional screens is to organize genes into pathways or complexes. To this end, we used Perturb-seq to perform high-resolution clustering of genetic perturbations. From a single assay, we recapitulated thousands of known relationships while also assigning new, experimentally validated roles to genes involved in ribosome biogenesis or translation (*CCDC86*, *CINP*, *SPATA5L1*, *ZNF236*, *C1orf131*, *SPOUT1*, *TMA16*, *NOPCHAP1*, *NEPRO*), transcription (*C7orf26*), and respiration (*TMEM242*). However, other large-scale experimental techniques, such as protein-protein interaction mapping, genetic interaction mapping, and co-essentiality analysis, similarly group genes or proteins by function. How then are single-cell CRISPR screens distinct?

We argue that these screens are particularly powerful because of the intrinsic interpretability of comprehensive genotype-phenotype maps, enabling in-depth dissection of the functional consequences of genetic perturbations that impinge on many distinct aspects of cell biology. Of particular note is the ability to use the information-rich readouts to study complex, composite phenotypes, which are difficult to measure by other modalities. These composite phenotypes can be created in a data-driven (e.g., deriving transcriptional programs) or hypothesis-driven manner (e.g., measuring intron/exon ratios to study splicing), resulting in an enormous breadth of measured phenotypes. In the case of scRNA-seq, we show that it measures not only features such as differential gene expression and the activity of critical transcriptional programs, but also RNA splicing and processing, expression of transposable elements, differentiation, transcriptional heterogeneity, cell cycle progression, and chromosomal instability. Once a phenotype is defined, the genotype-phenotype map can be used to explore its genetic underpinnings, in a manner analogous to a forward genetic screen, as well as its relationship to other cellular phenotypes.

An illustrative example of this process is our study of chromosomal instability. Based on an initial observation of heterogeneous responses to specific perturbations, we suspected that some cells carried genetically-induced chromosomal gains or losses. In a hypothesis-driven manner, we then used our rich phenotypic data to discover a large collection of perturbations—which were only loosely connected by clustering on average transcriptional phenotypes—that promote chromosomal instability. Importantly, the single-cell nature of our Perturb-seq data also allowed us to explore the relationship between karyotypic changes and other phenotypes, including cell cycle progression and stress induction. While aneuploidy is an important hallmark of most cancers, it has not been easy to study with traditional genetic screens as it requires both a single-cell and multimodal readout. In future work, this platform could be used to investigate interactions between genetic perturbations and specific karyotypes, karyotype-dependent stress responses, or the temporal evolution of karyotypes (*59*).

Genetic perturbations can push cells into extreme states that are not observed in unperturbed cells. Because composite phenotypes can be generated and explored without being preregistered at the time of data collection, rich genotype-phenotype maps provide a powerful resource for the discovery of new cellular behaviors. Using this ability, we discovered a remarkable degree of stress-specific changes in the expression of mitochondrially encoded transcripts. It was only possible to appreciate the functional specificity of this regulation by pairing a defined set of mitochondrial perturbations with a high-dimensional readout. This discovery suggests a framework to explain how cells cope with diverse insults to mitochondria: a general nuclear response is layered over perturbation-specific changes in the expression level of mitochondrially encoded genes (Fig. 7F). Building on this observation, we can ask new questions about the mitochondrial stress response. The transcriptional changes we observed may reflect adaptive responses or, alternatively, complex patterns of dysfunction owing to disruption of the intricate system of mitochondrial gene expression. Understanding how and in what contexts this regulation is adaptive may have important implications for diseases associated with mitochondrial stress. An intriguing additional question is whether individual mitochondria are able to regulate their expression autonomously. Combined with the nuanced responses observed here, this would support and substantially extend the “co-location for redox regulation” (CoRR) hypothesis which holds that the endosymbiotically derived mitochondrial genome has been retained through evolution to enable localized regulation of mitochondrial gene expression (*60*).

A final theme emerging from our work is the flexibility of single-cell CRISPR screens compared to other functional genomic approaches. Because these screens extract rich information from each cell in a pooled format, they require only a fraction of the number of cells used by other approaches and thus are well suited to the study of iPSC-derived cells and *in vivo* samples. As technologies for single-cell, multimodal phenotyping advance, single-cell screens will continue to become more powerful. At present, the major limitation of single-cell CRISPR screens is cost. Careful experimental designs, such as multiplexed libraries or compressed sensing (*61*), together with advances in single-cell phenotyping (*62, 63*) and DNA sequencing promise to greatly increase the scale of these experiments. To this point, we concluded our work by sequencing our genome-scale K562 libraries on a lower-cost, ultra-high throughput sequencing platform developed by Ultima Genomics, generating results equivalent to those sequenced on Illumina instruments (fig. S13).

In sum, our study presents a blueprint for the construction and analysis of rich genotype-phenotype maps to serve as a driving force for the systematic exploration of genetic and cellular function.

## Materials and Methods

A complete description of our Material and Methods is found in the Supplementary Material online. This includes methods experimental methods related to Perturb-seq screens and functional experiments, as well as computational methods detailing all data analysis.

## Acknowledgements

We thank S. Vazquez, L. Gilbert, F. Urnov, C. Cotta-Ramusino, V. Sankaran, K. Loh, W. Allen, B. Do, P. Hsu, F. Diehl, C. Jan, J. Replogle, D. Phizicky, and all members of the Weissman and Norman labs for helpful discussions. We thank Jorge Dinis and the Innovative Genomics Institute for critical administrative and scientific support. We also thank: the UCSF Center for Advanced Technology, Eric Chow, and Delsy Martinez for sequencing and access to FACS machines; the Whitehead Institute Flow Cytometry Core and Kathy Daniels for access to FACS machines; the Whitehead Institute Genome Technology Core for support with bulk RNA-seq library preparation and rRNA bioanalyzers; and the Harvard Nascent Transcriptomics Core for support with PRO-seq library preparation.

## Author contributions

JMR, RAS, TMN, and JSW were responsible for the conception, design, and interpretation of the experiments and wrote the manuscript. JMR led the Perturb-seq screens. JMR and TMN led the Perturb-seq data analysis. RAS led the functional studies. ANP generated and validated the ZIM3 CRISPRi RPE1 cell line and optimized Perturb-seq protocols across cell lines. JAH assisted with data interpretation and analysis. AL designed the interactive web visualization. AG produced preliminary data for Perturb-seq across cell lines. EJW and LM performed and supervised Drosophila Integrator biochemistry. KA supervised PRO-seq library preparation. GY, NI, FO, and DL sequenced libraries on the Ultima Genomics platform. JLB and MJ cloned pJB108 and validated the ZIM3 KRAB domain. JMR and RAS helped obtain funding for experiments. All authors provided feedback on the manuscript.

## Data and materials availability

Raw sequencing data will be deposited into SRA. An interactive data browser including processed, downloadable single-cell and pseudobulk populations will be made available upon publication. Our previously published analytic framework for Perturb-seq analysis is available at https://github.com/thomasmaxwellnorman/Perturbseq_GI. Scripts for guide assignment are available at https://github.com/josephreplogle/guide_calling. Additional code related to specific analyses will be made available on github upon publication.

## Supplementary Materials

### Materials and Methods

Figs. S1 to S13

Tables S1 to S9

