## Supplementary Methods and Figures for "Mapping information-rich genotype-phenotype landscapes with genome-scale Perturb-seq"

**This file includes:**

Materials and Methods
Figs. S1 to S13

**Other Supplementary Materials for this manuscript include the following:**

Tables S1 to S9

### Materials and Methods

### EXPERIMENTAL METHODS

#### Cell culture and lentiviral production

K562 cells were grown in RPMI-1640 with 25 mM HEPES, 2.0 g/l NaHCO3, and 0.3 g/l L-glutamine supplemented with 10% FBS, 2 mM glutamine, 100 units/ml penicillin, and 100 µg/ml streptomycin. hTERT-immortalized RPE1 cells (ATCC, CRL-4000) were grown in DMEM:F12 supplemented with 10% FBS, 0.01 mg/ml hygromycin B, 100 units/ml penicillin, and 100 µg/ml streptomycin. HEK293T cells were used for generation of lentivirus, and grown in DMEM supplemented with 10% FBS, 100 units/ml penicillin and 100 µg/ml streptomycin. Lentivirus was produced by co-transfecting HEK293T cells with transfer plasmids and standard packaging vectors using TransIT-LTI Transfection Reagent (Mirus, MIR 2306).

#### Cell line generation

CRISPRi K562 cells expressing dCas9-BFP-KRAB (KOX1-derived) were obtained from Gilbert *et al.*, 2014 (*23*). CRISPRi RPE1 cells expressing dCas9-BFP-KRAB (KOX1-derived) were obtained from Jost *et al.*, 2017 (*64*) and only used for growth screens. CRISPRi RPE1 cells were generated by stably transducing RPE1 cells (ATCC, CRL-4000) with lentivirus expressing ZIM3 KRAB-dCas9-P2A-BFP from a UCOE-SFFV promoter (pJB108) and sorting for BFP+ cells stably expressing the construct using fluorescence activated cell sorting. Cell lines were verified by monitoring BFP fluorescence over several generations to confirm stable integration and confirming knockdown of select surface markers by flow cytometry.

#### Library design and cloning

A distinct set of genes was targeted for each of the three large-scale Perturb-seq experiments. For the K562 day 8 genome-scale experiment, we targeted (i) genes expressed in K562 cells (ii) transcription factors as d(Lambert et al., 2018)l., 2018 (*65*) (iii) Cancer Dependency Map common essential genes as defined in 20Q1 (iv) non-targeting control sgRNAs accounting for 5% of the total library. To define expressed genes in K562 cells, we used a combination of bulk RNA-seq data from ENCODE (https://www.encodeproject.org/files/ENCFF717EVE/) and 10x Genomics 3’ single-cell RNA-seq data (https://www.ncbi.nlm.nih.gov/geo/query/acc.cgi?acc=GSE146194), selecting a set of genes accounting for ~99% of aligned reads in both datasets. For the K562 day 6 essential-scale experiment, we targeted (i) Cancer Dependency Map common essential genes as defined in 20Q1 (ii) non-targeting control sgRNAs accounting for 5% of the total library. For the RPE1 day 7 essential-scale experiment, we targeted (i) 20Q1 Cancer Dependency Map common essential genes (https://depmap.org/portal/download/) (ii) a number of hand-selected genes with interesting phenotypes in the K562 genome-wide Perturb-seq dataset (iii) non-targeting control sgRNAs accounting for 5% of the total library. To define control perturbations, we randomly sampled non-targeting control perturbations from Horlbeck *et al.,* 2016 (*66*). A small number of genes were lost in this pipeline due to changes in gene annotation between datasets.

To minimize library size while maximizing knockdown, multiplexed CRISPRi libraries were constructed which targeted each gene with two unique sgRNAs expressed from tandem U6 expression cassettes in a single lentiviral vector, as previously described in Replogle *et al.*, 2020

(*13*). The Horlbeck *et al.*, 2016 CRISPRi sgRNA libraries were used as a source of sgRNAs targeting each gene, with the optimal sgRNA pair targeting each gene selected based on a balance of empirical data with computational predictions. For strong essential genes (defined by a p-value<0.001 and gamma<-0.2 in the Horlbeck *et al.*, 2016 CRISPRi growth screen), sgRNAs were ranked by growth. Then, for genes that produced a significant phenotype in previous CRISPRi screens, sgRNAs were ranked by a discriminant score multiplying the negative $\log_{10}$ p-value by the effect size. Finally, for genes without any empirical evidence, sgRNAs were ranked according to the Horlbeck *et al.,* 2016 hCRISPRi v2.1 algorithm. The full sgRNA content of the K562 day 8 genome-scale library, K562 day 6 essential-wide library, and RPE1 day 7 essential-wide library can be found in Tables S1, S2, and S3, respectively.

We adapted the protocol previously described in Replogle *et al.,* 2020 to clone libraries with capture sequences for 3' direct capture Perturb-seq. Briefly, an sgRNA lentiviral expression vector (pRS275/pJR101) was derived from the parental pJR85 (Addgene #140095), modified to incorporate a GFP fluorescent marker to avoid spectral overlap with BFP+ CRISPRi constructs and a UCOE element upstream of the EF1alpha promoter to prevent silencing. A two-step restriction enzyme digestion and ligation cloning of oligos into pRS275/pJR101 was performed to maintain coupling of sgRNAs targeting the same gene. Oligos encoding the targeting regions of dual-sgRNA pairs were synthesized as an oligonucleotide pool (Twist Bioscences) with the structure: 5'- PCR adapter - CCACCTTGTTG – targeting region A - gtttcagagcgagacgtgcctgcaggatacgtctcagaaacatg – targeting region B - GTTTAAGAGCTAAGCTG - PCR adapter-3'. When ordering oligos, the representation of essential genes was increased to compensate for growth phenotypes (see below). Oligo pools were amplified, digested with BstXI/BlpI, and ligated into pRS275/pJR101. To add an sgRNA constant region and U6 promoter to the vector, pJR89 (Addgene #140096) was BsmBI-digested and ligated into the intermediate library.

**K562 and RPE1 growth screens**

Pooled sgRNA growth screens in K562 cells were used to quantify growth phenotypes of sgRNA pairs targeting expressed genes. CRISPRi K562 cells expressing dCas9-BFP-KRAB were transduced with lentiviral particles encoding the dual-sgRNA library by spinfection (1000g) with polybrene (8 ug/ml) to obtain an infection rate of ~25%-35%. Screens were performed in biological replicate with the aim of maintaining 1000 cells per library element for the duration of the screen. Between day 2 and day 6 post-transduction, cells were selected for lentiviral infection using 1 ug/mL puromycin, replenished every 24 hours. On day 7 post-transduction, an aliquot of cells was harvested as an initial time point) The rest of the cell population was passaged for 10 more days and collected at final time point.

Pooled sgRNA growth screens in RPE1 cells were used to quantify growth phenotypes of sgRNA pairs targeting common essential genes. The CRISPRi RPE1 cell line expressing dCas9-BFP-KRAB was used for growth screens which took place before the publication of the next-generation ZIM3 KRAB domain. Cells were transduced in biological replicate with lentiviral particles encoding the dual-sgRNA library by replating cells into virus-laden media with polybrene (8 ug/ml) to obtain an infection rate of ~45%. Because RPE1 cells are puromycin resistant, we performed the screen without selection for sgRNA-infected cells, nonetheless maintaining an infection rate-corrected 1000 cells per library element for the duration of the screen. On day 6 post-transduction, an aliquot of cells was harvested as an final time point for direct comparison to the abundances in the plasmid library.

For both K562 and RPE1 growth screens, DNA libraries of the initial and final samples were prepared for deep sequencing by genomic DNA isolation and PCR amplification of dual-sgRNA amplicons. First, a NucleoSpin Blood XL kit (Macherey–Nagel) was used to extract genomic DNA (gDNA) from cells. Then, isolated gDNA or plasmid DNA was amplified by 22 cycles (gDNA) or 13 cycles (plasmid DNA) of PCR using NEBNext Ultra II Q5 PCR MasterMix (NEB), appending Illumina adaptors and sample indices (oJR234 forward primer: 5’-AATGATACGGCGACCACCGAGATCTACACCGCGGTCTGTATCCCTTGGAGAACCACCT-3’; index primers 5’-CAAGCAGAAGACGGCATACGAGATnnnnnGCGGCCGGCTGTTTCCAGCTTAGCTCTTAAA-3’). Amplicons were isolated by a 0.5-0.65X SPRI bead selection (SPRIselect Beckman Coulter #B23318). Sequencing was performed on a NovaSeq 6000 (Illumina) using a 19 bp read 1, 19 bp read 2, and 5 bp index read 1 with custom sequencing primers oJR326 (custom read 1, 5’- CGCGGTCTGTATCCCTTGGAGAACCACCTTGTTGG-3’), oJR328 (custom read 2, 5’- GCGGCCGGCTGTTTCCAGCTTAGCTCTTAAAC-3’), and oJR327 (custom index read 1, 5’- GTTTAAGAGCTAAGCTGGAAACAGCCGGCCGC-3’).

**Perturb-seq experiments**

To perform our K562 day 8 genome-scale Perturb-seq experiment, library lentivirus was packaged into lentivirus in 293T cells and empirically measured in K562 cells to obtain viral titers. CRISPRi K562 cells were transduced via spinfection (1000g) with polybrene (8 ug/ml) with the target of obtaining an infection rate of ~10%. Cells were maintained at a viability of >90%, a coverage of 1000 cells per library element, and a density of 250,000 to 1,000,000 cells/ml for the course of the experiment. Three days post-transduction, an infection rate of 14% was measured, and cells were sorted to near purity by FACS (FACSAria2, BD Biosciences), using GFP as a marker for sgRNA vector transduction. Eight days post infection, the cells were measured to be 97% GFP+ (LSR2, BD Biosciences), >90% viable, and at a concentration of ~800,000 cells/ml (Countess II, ThermoFisher). Cells were prepared for single-cell RNA-sequencing by resuspension in 1X PBS with 0.04% BSA as detailed in the 10x Genomics Single Cell Protocols Cell Preparation Guide (10x Genomics, CG00053 Rev C). Cells were then separated into droplet emulsions using the Chromium Controller (10x Genomics) with Chromium Single-Cell 3′ Gel Beads v3 (10x Genomics, PN-1000075 and PN-1000153) across 273 “lanes”/”GEM groups” following the 10x Genomics Chromium Single Cell 3′ Reagent Kits v3 User Guide with Feature Barcode technology for CRISPR Screening (CG000184 Rev C) with the goal of recovering ~15,000 cells per GEM group before filtering. Because the formation of droplet emulsions occurred in batches of 8 GEM groups over several hours, fresh populations of cells were obtained every hour to prevent alterations in single-cell transcriptomes.

To perform our K562 day 6 essential-scale Perturb-seq experiment, library lentivirus was packaged into lentivirus in 293T cells and empirically measured in K562 cells to obtain viral titers. CRISPRi K562 cells were transduced via spinfection (1000g) with polybrene (8 ug/ml) with the target of obtaining an infection rate of ~10% with maintenance of cells as described above. Three days post-transduction, an infection rate of 15% was measured, and cells were sorted to near purity by FACS (FACSAria2, BD Biosciences), using GFP as a marker for sgRNA vector transduction. Six days post infection, the cells were measured to be 93% GFP+ (LSR2, BD Biosciences), >90% viable, and at a concentration of ~600,000 cells/ml (Countess II, ThermoFisher). Cells were prepared for single-cell RNA-sequencing by resuspension in 1X PBS with 0.04% BSA as detailed in the 10x Genomics Single Cell Protocols Cell Preparation Guide (10x Genomics, CG00053 Rev C). Cells were then separated into droplet emulsions using the Chromium Controller (10x Genomics) with Chromium Single-Cell 3′ Gel Beads v3 (10x Genomics, PN-1000075 and PN-1000153) across 48 "lanes"/"GEM groups" following the 10x Genomics Chromium Single Cell 3′ Reagent Kits v3 User Guide with Feature Barcode technology for CRISPR Screening (CG000184 Rev C) with the goal of recovering ~15,000 cells per GEM group before filtering.

To perform our RPE1 day 7 essential-scale Perturb-seq experiment, library lentivirus was packaged into lentivirus in 293T cells and empirically measured in RPE1 cells to obtain viral titers. CRISPRi RPE1 cells expressing ZIM3 KRAB-dCas9-P2A-BFP were transduced via replating into virus-laden media with polybrene (8 ug/ml) with the target of obtaining an infection rate of ~10%. Three days post-transduction, an infection rate of 7% was measured, and cells were sorted to near purity by FACS (FACSAria2, BD Biosciences), using GFP as a marker for sgRNA vector transduction. Seven days post infection, the cells were measured to be 86% GFP+ (LSR2, BD Biosciences) and >95% viable (Countess II, ThermoFisher). After trypsin dissociation, cells were prepared for single-cell RNA-sequencing by resuspension in 1X PBS with 0.04% BSA as detailed in the 10x Genomics Single Cell Protocols Cell Preparation Guide (10x Genomics, CG00053 Rev C). Cells were then separated into droplet emulsions using the Chromium Controller (10x Genomics) with Chromium Single-Cell 3′ Gel Beads v3 (10x Genomics, PN-1000075 and PN-1000153) across 56 "lanes"/"GEM groups" following the 10x Genomics Chromium Single Cell 3′ Reagent Kits v3 User Guide with Feature Barcode technology for CRISPR Screening (CG000184 Rev C) with the goal of recovering ~15,000 cells per GEM group before filtering.

**Perturb-seq library preparation and sequencing**

For preparation of gene expression and sgRNA libraries, samples were processed according to 10x Genomics Chromium Single Cell 3′ Reagent Kits v3 User Guide with Feature Barcode technology for CRISPR Screening (CG000184 Rev C). To allow for parallel library preparation, samples were arranged in 96-well plates with magnetic selections conducted on an Alpaqua Catalyst 96 plate (#A000550). For sequencing, mRNA and sgRNA libraries were pooled to avoid index collisions at a 10:1 ratio. Libraries were sequenced on both (i) a NovaSeq 6000 (Illumina) according to the 10x Genomics User Guide and (ii) the Ultima Genomics ultra-high throughput sequencing platform.

For sequencing on the Ultima Genomics (UG) platform, final 10x libraries were converted using conversion primers that anneal to the R1 and R2 regions of the 10X library and contain a UG-specific adapter sequence overhang and sample index. 8 PCR cycles were used for conversion. Converted libraries were bead purified and quantified. After pooling libraries, pools were seeded and clonally amplified on UG sequencing beads and sequenced on a UG prototype Sequencer. Single reads were generated from the 10x 3' libraries, reading the 10X cell barcode, UMI, and 3' end of the cDNA transcript. A specific sequencing protocol including a high volume of dT nucleotides was used to accommodate the high nucleotide consumption in the poly (dT) stretch of the cDNA. Following sequencing, the single reads were quality-trimmed and split into two sub-sequences corresponding to Read1 (10X cell barcode and UMI) and Read2 (cDNA), which were used as input to Cell Ranger for alignment.

**rRNA analyses**

K562s expressing Zim3-dCas9-2A-BFP were spinfected in biological duplicate (targeting sgRNAs) or quadruplicate (non-targeting sgRNAs) with lentivirus expressing GFP and an sgRNA. Two days after spinfection, the cells were sorted for GFP+ on a BD ARIA II. Sort purity was generally >95%. After the sort, cells were maintained in media supplemented with 4 ug/ml puromycin for four days and then recovered for two days. Cells were counted, collected by centrifugation, and harvested by vigorous vortexing in Tri Reagent (ThermoFisher AM9738).

RNA was extracted with chloroform according to the manufacturer's instructions, quantified by nanodrop, and snap frozen. Small samples were diluted to 200 ng/ul and run on Bioanalyzer RNA nano chips (Agilent 5067-1511) according to the manufacturer's instructions. Runs were aligned to the 18s peak and signal intensity was normalized to total RNA area.

**Integrator co-depletion**

K562s expressing Zim3-dCas9-2A-BFP were spinfected with lentivirus expressing GFP and an sgRNA. Two days after spinfection, the cells were sorted for GFP+ on a BD ARIA II. Sort purity was generally >95%. After the sort, cells were maintained in media supplemented with 4 ug/ml puromycin for four days and then recovered for two days. Cells were counted, washed twice with DPBS, and collected as pellets. The pellets were resuspended in SDS lysis buffer (100 mM Tris pH 8.0, 1% SDS), thermomixed at 95˚/1500 RPM for thirty minutes, aliquoted, and snap-frozen.

*Quantification for western blots*. An equal amount of material was loaded, as assessed by lysate A280.

**Integrator co-immunoprecipitation**

Human expression plasmids encoding codon-optimized INTS10 or His8-INTS10 were synthesized (Twist Bioscience) and transfected into HEK 293T/17 cells (ATCC CRL-11268) with FuGene HD (Promega E2311) according to the manufacturer's protocol. Two days later, the cells were washed twice with DPBS and harvested with IP lysis buffer (25 mM Tris-HCl pH 7.4, 150 mM NaCl, 1 mM EDTA, 1% NP-40, 5% glycerol; ThermoFisher 87787) supplemented with protease inhibitors (ThermoFisher A32965). Lysates were nutated at 4˚ for 30 mins, clarified by centrifugation at 12,000xg for 10 minutes, and snap-frozen. Concentrations were measured with the BCA assay (ThermoFisher 23225).

Lysates were thawed on ice, supplemented with imidazole to 10 mM, and nutated at 4˚ for 30 minutes with cobalt magnetic beads (ThermoFisher 10103D) pre-equilibrated in IP lysis buffer + 10 mM imidazole. The beads were separated on a magnet, washed twice with lysis buffer + 10 mM imidazole, and eluted with lysis buffer + 300 mM imidazole.

*Quantification for western blots*. For input samples, an equal amount of material was loaded, as assessed by BCA. For IP samples, an equal volume of eluate was loaded.

**Integrator purification**

Human expression plasmids encoding codon-optimized HIS-INTS10, INTS13, INTS14, and C7orf26 were synthesized (Twist Bioscience) and co-transfected with ExpiFectamine 293 (ThermoFisher A14524) into Expi293 cells (ThermoFisher A14527) maintained in Expi293 medium (ThermoFisher A1435101) according to the manufacturer's instructions. The cells were harvested after four days and snap frozen.

The pellets were resuspended in CHAPS Lysis Buffer (50 mM HEPES pH 8.0, 300 mM NaCl, 0.2% CHAPS, 10% glycerol, 1 mM TCEP, 1 mM EDTA, 0.5 mM PMSF, 1x protease inhibitors, 0.002% benzonase) and stirred at 4° for 30 minutes. The lysates were clarified at 120,000xg for 30 minutes, supplemented with 15 mM imidazole, and nutated for an hour with Ni-NTA agarose beads (ThermoFisher 25215) pre-equilibrated in CHAPS lysis buffer + 15 mM imidazole. The beads were loaded into a gravity column, washed with >10 volumes of wash buffer (50 mM HEPES pH 8.0, 300 mM NaCl, 10% glycerol, 1 mM TCEP, 1 mM EDTA, 15 mM imidazole), and eluted with wash buffer supplemented with 250 mM imidazole. The eluate was concentrated and buffer exchanged into SEC buffer (50 mM HEPES pH 8.0, 150 mM KCl, 10% glycerol, 1 mM EDTA) by ultrafiltration, and snap frozen.

The eluate was thawed on ice, passed through a 0.2 μM PES filter, and loaded onto an Superdex 200 Increase 10/300 GL column pre-equilibrated with SEC buffer. Fractions were collected and flash frozen.

*Quantification for gels and western blots*. An equal volume of sample from each SEC fraction was loaded. Less Ni-NTA eluate was loaded to account for the dilution over SEC.

**Drosophila Integrator biochemistry**

*Stable cell lines and nuclear extract preparation.* Relevant *Drosophila* cDNAs were cloned into a pMT-3xFLAG-puro plasmid (*67*, *68*) following the metallothionein promotor and 3x-FLAG tag. $2x10^6$ Drosophila DL1 cells were plated in Schneider's media supplemented with 10% FBS in a 6-well plate overnight and 2 μg of plasmid was transfected using Fugene HD (Promega, Madison WI, #E2311). Plasmid DNA was mixed with 8 μL Fugene and 100 μL media and incubated at room temperature for 15 minutes before being added to cells. After 24 hours, 2.5 μg/mL puromycin was added to the media to select and maintain the cell population. Cells were transitioned to SFX media without serum for large scale growth. Protein expression for nuclear extract was induced by adding 500 mM copper sulfate for 48 hours to 1 liter of each cell line grown to approximately $1x10^7$ cells/mL.

Cells were collected and washed in cold PBS and then pelleted by centrifugation. Cells were then resuspended in five times the cell pellet volume of Buffer A (10mM Tris pH8, 1.5 mM $MgCl_2$, 10 mM KCl, 0.5mM DTT, and 0.2mM PMSF). Resuspended cells were allowed to swell during a 15-minute rotation at 4°C. After pelleting down at 1,000g for 10 minutes, two volumes of the original cell pellet of Buffer A were added and cells were homogenized with a dounce pestle B for 20 strokes on ice. Nuclear and cytosolic fractions were then separated by centrifugation at 2,000g for 10 minutes. To attain a nuclear fraction, the pellet was washed once with Buffer A before resuspending in an equal amount of the original cell pellet volume of Buffer C (20 mM Tris pH8, 420mM NaCl, 1.5 mM $MgCl_2$, 25% glycerol, 0.2 mM EDTA, 0.5 mM PMSF, and 0.5 mM DTT). The sample was then homogenized with a dounce pestle B for 20 strokes on ice and rotated for 30 minutes at 4°C before centrifuging at 15,000g for 30 minutes at 4°C. Finally, supernatants were collected and subjected to dialysis in Buffer D (20 mM HEPES, 100 mM KCl, 0.2 mM EDTA, 0.5 mM DTT, and 20% glycerol) overnight at 4°C. Prior to any downstream applications, nuclear extracts were centrifuged again at 15,000g for 3 minutes at 4°C to remove any precipitate.

*Anti-FLAG affinity purification and western blotting.* To purify FLAG-tagged Integrator complexes for mass spectrometry, generally between 8 and 10 mg of DL1 nuclear extract (approximately 1.9 mL of extract depending on the concentration) was mixed with 100 µL anti-Flag M2 affinity agarose slurry (Sigma-Aldrich, #A2220) washed with 0.1 M glycine then equilibrated in binding buffer (20 mM HEPES pH7.4, 150 mM KCl, 10% Glycerol, 0.1% NP-40). This mixture was rotated for four hours at 4°C. Following the four-hour incubation/rotation, five sequential washes were carried out in binding buffer with a 10-minute rotation at 4°C followed by a 1,000g centrifugation at 4°C. After a final wash with 20 mM HEPES buffer, the supernatant was removed using a pipette and the beads were kept cold and submitted to the mass spectrometry core where the protein complexes were eluted by digestion (described below). For immunoprecipitation samples intended for western blot, a similar protocol was used. 25 µL of bead slurry and 200 µL of extract sample were rotated for two hours at 4°C. After the fifth wash with binding buffer, protein complexes were eluted from the anti-FLAG resin by adding 50 µL of 2X SDS loading buffer and boiled at 95°C for five minutes. For western blots, input samples were generated by adding equal volume of 2X SDS loading buffer to nuclear extract and 1/10 of the immunoprecipitation was loaded as estimated by protein mass. Total protein was resolved on SDS polyacrylamide gels (Bio-Rad) with DTT, followed by transfer onto polyvinylidene difluoride (PVDF) membranes (ThermoFisher). Blots were probed as previously described using *Drosophila*-specific antibodies raised against recombinant GST fusion proteins expressed in E. coli (*68*).

*Mass spectrometry sample digestion.* The samples were prepared in a similar manner as described previously (*69*). Briefly, the agarose bead-bound proteins were washed several times with 50 mM Triethylammonium bicarbonate (TEAB) pH 7.1, before being solubilized with 40 µL of 5% SDS, 50 mM TEAB, pH 7.55 followed by a room temperature incubation for 30 minutes. The supernatant containing the proteins of interest was then transferred to a new tube, reduced by making the solution 10 mM Tris(2-carboxyethyl)phosphine (TCEP) (Thermo, #77720), and further incubated at 65°C for 10 minutes. The sample was then cooled to room temperature and 1 µL of 1M iodoacetamide acid was added and allowed to react for 20 minutes in the dark. Then, 5 µL of 12% phosphoric acid was added to the 50 µL protein solution followed by 350 µL of binding buffer (90% Methanol, 100 mM TEAB final; pH 7.1). The resulting solution was administered to an S-Trap spin column (Protifi, Farmingdale NY) and passed through the column using a bench top centrifuge (30 second spin at 4,000g). The spin column was then washed three times with 400 µL of binding buffer and centrifuged (1200 rpm, 1 min). Trypsin (Promega, #V5280) was then added to the protein mixture in a ratio of 1:25 in 50 mM TEAB, pH=8, and incubated at 37°C for 4 hours. Peptides were eluted with 80 µL of 50 mM TEAB, followed by 80 µL of 0.2% formic acid, and finally 80 µL of 50% acetonitrile, 0.2% formic acid. The combined peptide solution was then dried in a speed vacuum (room temperature, 1.5 hours) and resuspended in 2% acetonitrile, 0.1% formic acid, 97.9% water and aliquoted into an autosampler vial.

*NanoLC MS/MS Analysis.* Peptide mixtures were analyzed by nanoflow liquid chromatography-tandem mass spectrometry (nanoLC-MS/MS) using a nano-LC chromatography system (UltiMate 3000 RSLCnano, Dionex, Thermo Fisher Scientific, San Jose, CA). The nano-LC-MS/MS system was coupled on-line to a Thermo Orbitrap Fusion mass spectrometer (Thermo Fisher Scientific,

San Jose, CA) through a nanospray ion source (Thermo Scientific). A trap and elute method was used to desalt and concentrate the sample, while preserving the analytical column. The trap column (Thermo Scientific) was a C18 PepMap100 (300 µm X 5 mm, 5 µm particle size) while the analytical column was an Acclaim PepMap 100 (75 µm X 25 cm) (Thermo Scientific). After equilibrating the column in 98% solvent A (0.1% formic acid in water) and 2% solvent B (0.1% formic acid in acetonitrile (ACN)), the samples (2 µL in solvent A) were injected onto the trap column and subsequently eluted (400 nL/min) by gradient elution onto the C18 column as follows: isocratic at 2% B, 0-5 min; 2% to 32% B, 5-39 min; 32% to 70% B, 39-49 min; 70% to 90% B, 49-50 min; isocratic at 90% B, 50-54 min; 90% to 2%, 54-55 min; and isocratic at 2% B, until the 65 minute mark.

All LC-MS/MS data were acquired using XCalibur, version 2.1.0 (Thermo Fisher Scientific) in positive ion mode using a top speed data-dependent acquisition (DDA) method with a 3 second cycle time. The survey scans (m/z 350-1500) were acquired in the Orbitrap at 120,000 resolution (at m/z = 400) in profile mode, with a maximum injection time of 100 m s and an AGC target of 400,000 ions. The S-lens RF level was set to 60. Isolation was performed in the quadrupole with a 1.6 Da isolation window, and CID MS/MS acquisition was performed in profile mode using rapid scan rate with detection in the ion-trap using the following settings: parent threshold = 5,000; collision energy = 32%; maximum injection time 56 msec; AGC target 500,000 ions. Monoisotopic precursor selection (MIPS) and charge state filtering were on, with charge states 2-6 included. Dynamic exclusion was used to remove selected precursor ions, with a +/- 10 ppm mass tolerance, for 15 seconds after acquisition of one MS/MS spectrum.

*Database Searching.* Tandem mass spectra were extracted and charge state deconvoluted using Proteome Discoverer (Thermo Fisher, version 2.2.0388). Deisotoping was not performed. All MS/MS spectra were searched against the Uniprot *Drosophila* database (version 04-04-2018), using Sequest. Searches were performed with a parent ion tolerance of 5 ppm and a fragment ion tolerance of 0.60 Da. Trypsin was specified as the enzyme, allowing for two missed cleavages. Fixed modification of carbamidomethyl (C) and variable modifications of oxidation (M) and deamidation were specified in Sequest. Heat maps in Figure S9A were made using Morpheus from the Broad Institute, https://software.broadinstitute.org/morpheus. Volcano plots in Figure S9B were generated using average number of peptide counts quantified using mass spectrometry for three independent measurements of purifications, which also was the basis for adjusted p-values. These values were all calculated and plotted using GraphPad Prism software.

### Integrator PRO-seq

PRO-seq was conducted largely according to published protocols with slight modifications (*70*). K562s expressing dCas9-BFP-KRAB were spinfected with lentivirus expressing GFP and an sgRNA. Two days after spinfection, the cells were sorted for GFP+ on a BD ARIA II. Sort purity was generally >95%. After the sort, cells were maintained in media supplemented with 4 ug/ml puromycin for three days and then recovered for two days.

Cells were counted, harvested by centrifugation, and washed with cold DPBS. All subsequent steps took place at 4° or on ice. All solutions were made with RNase-free reagents and were 0.2 µm filtered and chilled before use. 12 million cells were pelleted by centrifugation, resuspended in 250ul of buffer W (10 mM Tris pH 8.0, 10 mM KCl, 250 mM sucrose, 5 mM MgCl2, 1 mM EGTA, 0.5 mM DTT, 10% glycerol, 1x protease inhibitor [ThermoFisher A32965], and 0.02% v/v

SUPERase-In RNase inhibitor [AM2694], strained, and transferred to conical tubes that had been coated with 1% BSA in PBS overnight. The cells were permeabilized by dilution in 10 ml of buffer P (buffer W + 0.1% v/v Igepal CA-630 + 0.05% v/v Tween-20) and incubated for 5 minutes. The permeabilized cells were harvested by centrifugation at 400xg for 5 minutes, resuspended in 10 mL of buffer W, harvested by centrifugation at 400xg for 5 minutes, and resuspended in 250 ul buffer F (50 mM Tris pH 8.0, 40% v/v glycerol, 5 mM MgCl2, 1.1 mM EDTA, 0.5 mM DTT, 0.02% v/v SUPERase-In RNase inhibitor). ≥97% permeabilization efficiency was confirmed on a NucleoCounter NC-202 and permeabilized cells were snap frozen.

PRO-seq libraries were generated and sequenced by the Nascent Transcriptomics Core at Harvard Medical School according to their standard protocol. PRO-seq data were aligned and quantified using STAR (version 2.7.9a) with parameters alignEndsType=Local, outFilterMultimapNmax=20, outFilterScoreMinOverLread=0.3, and outFilterMatchNminOverLread=0.3. For comparison of gene-level expression profiles, gene counts were normalized for sequencing depth (reads per million), log-transformed, and subset to well-expressed genes (n=758 genes with >3000 rpm). Then, Spearman's correlation was used to compare the similarity of expression profiles.

**SDS-PAGE and western blotting**

Samples were mixed with sample loading buffer (Licor 928-40004) supplemented with DTT and incubated at 95˚ for 5 minutes. SDS-PAGE was performed with pre-cast 4-12% gradient gels (ThermoFisher NW04127BOX) in MOPS (ThermoFisher B000102) according to the manufacturer's instructions.

For Coomassie staining, gels were washed thoroughly in water, incubated with ReadyBlue Protein Gel Stain (Sigma RSB-1L) overnight, and destained in water. For western blots, proteins were transferred to nitrocellulose membranes by semi-dry transfer (Biorad 1704158) according to the manufacturer's instructions. Membranes were rinsed in water and stained with Revert 700 Total Protein Stain according to the manufacturer's instructions. The membranes were then rinsed in TBS, rocked with Everyblot Blocking Buffer (Biorad 12010020) at room temperature for > 30 minutes, and rocked with primary antibody overnight at 4˚. The membranes were washed with TBST, and rocked with IR800CW-labeled secondary antibodies for 30-60 minutes, washed with TBST, and imaged on a Licor Odyssey CLx.

**CD11b cell surface staining**

K562s expressing dCas9-BFP-KRAB were co-spinfected with lentiviruses expressing GFP-sgKDM1A and mCherry-sgPTPN1. Eight days after spinfection, the cells were counted and harvested by centrifugation. Cells were washed with PBE buffer (DPBS + 0.5% BSA + 2 mM EDTA) and resuspended with α-CD11b-AF647 antibody diluted 1:50 in PBE. Cells were incubated at 4˚ in the dark for 30 minutes, washed twice with PBE, and analyzed on a BD LSRFortessa. The populations were gated from a single sample as sgKDM1A (GFP+,mCherry-), sgPTPN1 (GFP-, mCherry+), and sgKDM1A/sgPTPN1 (GFP+, mCherry+). Unstained K562s expressing either GFP or mCherry were used as single color compensation controls. AF647 was compensated with UltraComp eBeads Plus (Thermo 01-3333-42) labeled with α-CD11b-AF647.

**Internally controlled growth assays**

K562s expressing Zim3-dCas9-2A-BFP were co-spinfected in triplicate with lentiviruses expressing GFP-sgKDM1A and mCherry-sgPTPN1, or with lentivirus expressing GFP and a non-targeting sgRNA. Every two days, cells were analyzed for BFP, GFP, and mCherry on an Attune flow cytometer. Enrichment was calculated as sgKDM1A (BFP+, GFP+, mCherry-), sgPTPN1 (BFP+, GFP-, mCherry+), and sgKDM1A/sgPTPN1 (BFP+, GFP+, mCherry+) vs uninfected (BFP+, GFP-, mCherry-).

**Bulk RNA-seq**

K562s expressing dCas9-BFP-KRAB were spinfected in biological duplicate with lentivirus expressing GFP and an sgRNA. Two days after spinfection, the cells were sorted for GFP+ on a BD ARIA II. Sort purity was generally >95%. After the sort, cells were maintained in media supplemented with 4 ug/ml puromycin for four days and then recovered for two days. Cells were counted, collected by centrifugation, and harvested by vigorous vortexing in Qiazol (Qiagen 79306).

Total RNA was extracted with miRNeasy Mini columns (Qiagen 217004) according to the manufacturer's instructions and sequencing libraries were prepared with TruSeq Stranded Total RNA Library Prep Human/Mouse/Rat kits (Illumina 20020596) according to the manufacturer’s instructions. Libraries were sequenced 2x150 on a NovaSeq (Illumina).

Bulk RNA-seq data were aligned and quantified using STAR (version 2.7.9a) with parameters alignEndsType=Local and outFilterMultimapNmax=20. For comparison of gene-level expression profiles, gene counts were corrected for sequencing depth (reads per million), and the $\log_2$ fold-change for each gene was calculated relative to within-replicate non-targeting control expression. The two replicates for each genetic perturbation were averaged in order to produce the final data.

**Seahorse experiment**

K562s expressing dCas9-BFP-KRAB were spinfected with lentivirus expressing GFP and an sgRNA. Two days after spinfection, the cells were sorted for GFP+ on a BD ARIA II. Sort purity was generally >95%. After the sort, cells were maintained in media supplemented with 4 ug/ml puromycin for four days and then recovered for three days. On the 9th day post spinfection, seahorse assay were plates were treated with Cell-Tak (Corning 354240) according to the manufacturer’s instructions. Cells were counted, collected by centrifugation, and resuspended in supplemented Seahorse XF RPMI (Agilent 103576-100). 150,000 cells were added to the Seahorse assay plate and attached via centrifugation at 200xg for 1 minute with no brake. After 30 minutes of recovery at 37˚, the cells were subjected to a Mito Stress Test on a Seahorse XFe96 analyzer according to the manufacturer’s instructions.

### COMPUTATIONAL METHODS

**Alignment, cell calling, and guide assignment**

Cell Ranger 4.0.0 software (10x Genomics) was used for alignment of scRNA-seq reads to the transcriptome, alignment of sgRNA reads to the library, collapsing reads to UMI counts, and cell calling. The 10x Genomics GRCh38 version 2020-A genome build was used as a reference transcriptome. For specific applications discussed below, STARsolo (STAR version 2.7.9a) was used extract transcript features, including intronic and exonic alignments and alignment of reads to transposable elements.

Reads from the sgRNA libraries were mapped with Cell Ranger. To account for differences in sequencing depths across GEM groups from the same experiment, reads were downsampled to produce a more even distribution of the number of reads per cell across gemgroups, with a threshold of 1000 reads per cell in the K562 day 8 experiment, 800 reads per cell in the K562 day 6 experiment, and 3000 reads per cell in the RPE1 experiment. Guide calling was performed with a Poisson-Gaussian mixture model as previously described. For each guide, the mixture model was fit 100 times, selecting the maximum likelihood model from among the fits. After guide calling, each cell was categorized according to its guide identities as representing a single genetic perturbation or a multiplet (which may arise from lentiviral recombination or multiple cell encapsulation during droplet generation). Only cells bearing two guides targeting the same gene or a single guide were used for downstream analysis.

Downstream analyses were performed in Python, using a combination of numpy, scipy, Pandas, scikit-learn, pomegranate, infercnvpy, pygenometracks, scanpy and seaborn libraries.

**Filtering and internal normalization of gene expression measurements**

Our internal normalization approach is similar to the one described in (*9*). First, we identified "core" control sgRNAs. That is, within each experiment there are tens to hundreds of possible negative control sgRNAs that were synthesized to have similar base compositions to targeting sgRNAs (*66*). Some of these by chance induce detectable phenotypes. We constructed a minimal set of control sgRNAs that are largely indistinguishable from each other using the following procedure: (i) We take all cells bearing all possible non-targeting sgRNAs and represent them by the vector of genes with mean >1 UMI count per cell. (ii) We *z*-normalize the expression of these genes: i.e. we subtract the mean and divide by the standard deviation. (iii) For each gene, we test for equality of distribution using the Anderson-Darling test (scipy.stats.anderson_ksamp) between all possible pairs of non-targeting control sgRNAs. (E.g. In the genome-scale dataset, there are 585 possible control sgRNAs and therefore $\binom{585}{2} = 170280$ pairwise comparisons.) (iv) We adjust the resulting $p$-values for multiple hypothesis testing using the Benjamini-Hochberg procedure. (v) For each potential control sgRNA, we compute the average number of differentially expressed genes relative to all other potential control sgRNAs. (vi) We set a dataset-dependent threshold on the number of differentially expressed genes (8 in the genome-scale dataset and 30 in the "K562 essentials" and "RPE1 essentials" datasets, which were more deeply sequenced and so had more genes passing the expression threshold) and kept all potential control sgRNAs that fell below the threshold. For example, in the genome-scale dataset this resulted in 514 control sgRNAs.

Next, we filtered cells based on quality metrics. We first computed scale factors to adjust for variable sequencing depths across gemgroups: we examined all core control cells (which make up ~4% of all cells), computed factors that equalized the mean UMI counts within these cells across gemgroups, and then applied these factors to all cells in the gemgroup to produce adjusted UMI counts. We then applied two quality filters, ensuring that cells passed a minimum adjusted UMI content filter (genome-scale dataset: 2000 UMIs, K562 essentials/RPE1 essentials: 3000 UMIs) and a maximum mitochondrial RNA filter (genome-scale dataset: <25%, K562 essentials: <20%,

RPE1 essentials: <11%). (Mitochondrial RNA content is the fraction of total UMIs derived from mitochondrially-encoded genes.) These filter parameters were chosen by plotting adjusted UMI content vs. mitochondrial RNA content and manually setting thresholds that removed the low-quality cells.

Finally, we computed a normalized gene expression matrix for cells passing the quality filters via two normalization steps: (i) *UMI count normalization*: We scale expression within all cells so that their total UMI counts equal the median UMI count of core control cells within the experiment. (ii) *Relative z-normalization*: Within each gemgroup, for each gene, we compute the mean and standard deviation of expression within control cells and use these to *z*-normalize expression. In other words, if $x$ is the expression of a given gene, it is represented by the score $z = (x - \mu_{\text{control}})/\sigma_{\text{control}}$, where the mean and standard deviation are separately computed within each gemgroup.) The resulting scores should therefore be interpreted as "fraction of transcriptional effort" due to the UMI count normalization, with a scale set relative to control cells. Put simply, an expression score of +2 thus represents a gene expressed at a level 2 standard deviations above the mean in control cells.

**Examining effects of normalization on batch effects**

As described in the main text, we observed batch effects in the data (fig. S2). This variation appeared to track mostly with sets of 8 samples that went through the 10x Chromium instrument and library prep together, though the precise origin is unclear. To construct this figure, we normalized the data in two ways. *Raw data normalization:* (i) To adjust for variable sequencing depth, scale cellular UMI counts by factors chosen so that so that core control cells have the same total UMI counts across all gemgroups. (ii) Construct a gemgroup mean expression profile of all genes with mean >2 UMI counts per cell by averaging counts following normalization in previous step across all cells in the gemgroup. (iii) Scale the gemgroup mean expression profiles by dividing by their mean across all gemgroups. An expression value of 1 is then the mean across all cells across all gemgroups. *Internal z-normalization*. Normalize expression as described in previous section.

Fig. S2 compares the two normalization schemes. In both cases the ranges of the plot are chosen using seaborn's robust option (which sets the min and max to the $2^{\text{nd}}$ and $98^{\text{th}}$ percentile of the data). Genes are clustered based on the raw data normalization and are in the same order in both panels. The gemgroups are presented in order based on how samples were multiplexed while performing the experiment as indicated by the color groupings at the top.

**Energy distance test for identifying perturbations that induce altered transcriptional states**

To compare distributions of expression states, we used tests derived from energy statistics, which allow for testing of equality of distributions when data are high-dimensional. In short, each cell is represented by a vector composed of its top 20 principal component scores, and we compare whether the distribution of these 20-dimensional vectors is equal or not between unperturbed control cells and cells bearing each perturbation. When these distributions differ, we can infer that the perturbation is causing some change either in the structure or distribution of transcriptional states within the perturbed cells.

To construct the distributions to compare, we first applied a series of filtering steps: (i) we removed cells that did not pass the UMI or mitochondrial RNA filters described in *Internal normalization of gene expression measurements*; (ii) as features, we took the $z$-normalized expression of all genes with mean expression >0.5 UMIs per cell; (iii) to dampen the effects of a handful of strongly induced outlier genes, we clipped any measure with a $z$-score greater than 10 to 10 (this only affects a handful of genes); (iv) finally, we applied principal components analysis (using sklearn's PCA implementation, which will use randomized algorithms for datasets of this scale) and kept only the top 20 principal components. The test should therefore be interpreted as assessing gross changes in cellular transcriptional state.

To construct a null distribution, we randomly subsampled 5,000 control cells bearing non-targeting sgRNAs. (Subsampling was necessary for performance reasons.) For each perturbation, we then compute an estimator of the energy distance:

$$\mathcal{E}(x,y) = \frac{2}{n_1 n_2}\sum_{i=1}^{n_1}\sum_{j=1}^{n_2}\|\boldsymbol{x_i} - \boldsymbol{y_j}\| - \frac{1}{n_1^2}\sum_{i=1}^{n_1}\sum_{j=1}^{n_1}\|\boldsymbol{x_i} - \boldsymbol{x_j}\| - \frac{1}{n_2^2}\sum_{i=1}^{n_2}\sum_{j=1}^{n_2}\|\boldsymbol{y_i} - \boldsymbol{y_j}\|$$

where each $\boldsymbol{x_i}$ is one of the control cells and each $\boldsymbol{y_j}$ is one of the perturbed cells.
In the limit of infinite data, the energy distance will be 0 between identical distributions and positive between non-identical distributions. We assess statistical significance in practice using a permutation test by permuting the labels of control and perturbed cells 10,000 times and estimating how frequently a larger energy distance would be observed by chance. The specific implementation is based on the python package torch-two-sample, modified to use numba for improved performance.

**Gene-level differential expression testing using the Anderson-Darling and Mann-Whitney tests**

Because of (i) biological differences in expression characteristics across different genes, (ii) the batch effects described above, (iii) incomplete penetrance of some perturbations, and (iv) heterogeneity of some gene expression programs, we opted to use non-parametric statistical tests rather than tests based on specific distributional assumptions about gene expression. Specifically, we $z$-normalize gene expression relative to control cells as described (*Internal normalization of gene expression measurements*) and for each gene test whether the distribution of normalized expression is identical between control cells bearing non-targeting sgRNAs and cells bearing each perturbation. We used two tests implemented in scipy: the Anderson-Darling test (scipy.stats.anderson_ksamp), which is broadly sensitive to changes in distribution, and the Mann-Whitney U test (scipy.stats.mannwhitneyu), which tests whether one distribution is stochastically greater than another. For the Anderson-Darling test we extended the range of $p$ values beyond those available in scipy's implementation by computing the $p$-value for many values of the test statistic using R's kSamples package and interpolating any intermediate values using scipy.interpolate.interp1d. For the Mann-Whitney test we used the asymptotic $p$ values and excluded any perturbation with fewer than 10 cells. $p$-values in both cases were adjusted for multiple hypothesis testing using the Benjamini-Hochberg procedure to produce the final results.

**Global analysis and clustering of strong perturbations**

The analysis presented covers 1973 perturbations that met three criteria: (i) at least 50 differentially expressed genes at a significance of $p < 0.05$ by Anderson-Darling test following Benjamini-Hochberg correction; (ii) at least 25 cells that passed our quality filters; and (3) an on-target knockdown, if measured, of at least 30% (i.e. the target of perturbation was either knocked down by at least 30% or was not detected, a broad attempt to remove non-functional perturbations). As features, we used a union of two sets of genes: (i) the top 10 differentially expressed genes for all perturbations (ordered by the value of the Anderson-Darling test statistic) and (ii) all genes of mean >0.25 UMIs per cell with variance in the top 30% of the dataset. We represented perturbations by their mean normalized expression profile across these 2319 highly variable genes. To prevent the direct targets of knockdown influencing results, the target gene value was replaced by 0 for the corresponding perturbation. For example, RPS5 gene normalized expression was set to 0 in the expression profile of the RPS5 perturbation, which is equal to the mean in control cells by construction.

Because clearly related perturbations sometimes showed variable absolute phenotypic strengths, we used correlation as a metric to compare profiles, since it is scale-invariant. We conducted two global assessments of the ability of these expression profile correlations to recall known biological relationships. First, curated complexes were obtained from the 03.09.2018 CORUM3.0 database (*26*). We identified all complexes that had at least 66% of genes represented within the 1973 perturbations (based on matching gene symbols between the datasets), leading to 327 complexes. Each represented complex was then split into a series of links (e.g. if a complex contained genes A, B, and C, then it would be split into links A-B, B-C, and A-C). The figure plots the distribution of expression profile correlations of these annotated links versus the distribution of all possible links among the 1973 targeted genes. A similar analysis was then conducted using predicted protein links from the v11.5 of STRING (*27*) (9606.protein.links.v11.5.txt.gz) after mapping STRING protein IDs to gene names (using the "preferred_name" field in 9606.protein.info.v11.5.txt.gz). Among the 1973 genes in the figure there are 1,945,378 possible pairwise links between genes, 243,558 of which have scores within STRING. We binned these represented links into 6 equally spaced bins based on observed expression profile correlation. The figure shows kernel density estimates of the STRING score distribution within each bin made using seaborn's violinplot (with cut set to 0 so that density estimates do not extend past observed data).

To identify clusters of related perturbations, we manually computed correlation distances between all pairs of expression profiles, and used HDBSCAN (metric='precomputed', min_cluster_size=4, min_samples=1, cluster_selection_method='eom') to identify 63 clusters. This procedure is intrinsically conservative due to the choice of metric and clustering algorithm, so many perturbations are not assigned to any cluster—our emphasis was on identifying the strongest signals rather than the most comprehensive. We then annotated the possible function of these clusters using a combination of manual lookup of related genes and automated annotation using CORUM complexes and STRING clusters. (STRING clusters are derived from 9606.clusters.info.v11.5.txt.gz and are labeled according to the "best_described_by" field.) We only assigned automated annotations when a cluster contained 75% or more of the members of a CORUM complex or STRING cluster. Aggregated information about clusters is provided in Table S7 which has the following fields:

| | |
|---|---|
| **members** | The genes assigned to the cluster by HDBSCAN |

| | |
|---|---|
| **nearby_genes** | Perturbations that are close to the cluster center in the high-dimensional embedding (see below). These are candidate members of the cluster that may be too weak to be called by HDBSCAN, which is quite conservative. |
| **emb_variable_x** | Location of cluster center in Fig. 2D |
| **emb_variable_y** | Location of cluster center in Fig. 2D |
| **manual_annotation** | Manual annotation of cluster function. |
| **contained_string_cluster_ids** | IDs of any STRING clusters contained within this cluster (at least 75% of members must be represented). |
| **contained_string_clusters** | Descriptions of any STRING clusters contained within this cluster (at least 75% of members must be represented). |
| **contained_corum_complexes** | Names of any CORUM complexes contained within this cluster (at least 75% of members must be represented). |
| **nearby_corum_complexes** | Names of any CORUM complexes that are close to this cluster in the high-dimensional embedding (at least 75% of members must be represented). |
| **nearby_string_clusters** | Names of any STRING clusters that are close to this cluster in the high-dimensional embedding (at least 75% of members must be represented). |
| **strong_positive_gene_expr_clusters** | Gene expression programs (see Fig. 4) for which average expression in this cluster is at least 2 standard deviations above normal expression level. |
| **strong_negative_gene_expr_clusters** | Gene expression programs (see Fig. 4) for which average expression in this cluster is at least 2 standard deviations below normal expression level. |
| **best_description** | Either the manual annotation or identity as called by CORUM/STRING. Most of the labels in Fig. 2D come from this field. |
| **example_genes** | Manually curated genes indicative of cluster function. |
| **Notes** | Notes, including any known off-targets. |

**Minimum distortion embedding of strong perturbations**

The visualization in Fig. 2D is a minimum distortion embedding (MDE) of the 1973 strong perturbations created using pymde v0.1.13. pyMDE solves MDE problems based on minimizing Euclidean distances. To adapt it to correlation distances, we first *z*-normalized each expression profile. (I.e. If a perturbation is represented by the vector $\boldsymbol{x}$ of 2319 highly variable genes, we computed $\widehat{\boldsymbol{x}} = (\boldsymbol{x} - \langle \boldsymbol{x} \rangle)/\sigma_x$.) Because of the polarization identity, minimizing the Euclidean distance between these normalized profiles is equivalent to minimizing the (square root of) the correlation distance between the unnormalized profiles:

$$||\widehat{\boldsymbol{x}} - \widehat{\boldsymbol{y}}||^2 = ||\widehat{\boldsymbol{x}}||^2 + ||\widehat{\boldsymbol{y}}||^2 - 2(\widehat{\boldsymbol{x}} \cdot \widehat{\boldsymbol{y}}) = 2(1 - \mathrm{corr}(\boldsymbol{x}, \boldsymbol{y}))$$

where the second equality follows from the *z*-normalization and the scale-invariance of correlation.

We created two embeddings. First, we used pymde to embed the dataset into 20 dimensions. This "high-dimensional embedding" serves as an imputation step, as it distorts the geometry of the dataset so that clusters of related genes that may be driven by weaker overall correlations are allowed to form. Proximity within this embedding was used to identify genes, CORUM complexes, and STRING clusters that were near to the HDBSCAN clusters called on the raw data (which were well-preserved in the embedding) as described in Table S7. To construct the embedding, we initialized pymde using the spectral embedding of the dataset (using sklearn's SpectralEmbedding with n_components=20, affinity='nearest_neighbors', n_neighbors=7, eigen_solver='arpack')), and then ran pymde's "preserve_neighbors" function function with embedding_dim=20, n_neighbors=7, and repulsive_fraction=5. pymde was run until convergence with a final average distortion of 0.0979 and final residual norm of 9.4e-06.

To produce the embedding in Fig. 2D, we ran pymde with the same parameters but with the embedding dimension set to 2 (final average distortion 0.105, final residual norm 3.2e-06). The bold cluster labels in the figure correspond to the manual annotations mentioned in the previous section. A handful of changes were manually incorporated: (1) the cytochrome c-ubiquinol cluster was not detected by HDBSCAN, and was manually annotated (2) 4 clusters involving protein post-translational modifications (ubiquitination, sumoylation, acetylation, neddylation) were annotated with a single label of "post-translational modifications" (3) components of eIF3 split across two clusters that are next to each other in the embedding and are labeled as a single cluster (4) all clusters of unknown function were not labeled but are included in the supplemental tables. The complex labels come from CORUM or STRING. A complex/cluster label was placed if and only if 75% of the members of a complex or cluster were close to each other in the 20-dimensional pyMDE embedding ("close" meaning at or below the 5$^{th}$ percentile of all pairwise distances). Redundant/duplicated clusters were manually deduplicated. The locations of the labels on the figure were then adjusted for readability. Label location is a decent proxy for, but not an entirely accurate representation of, cluster and complex locations.

**Clustering of gene expression programs**

We next turned to identifying conserved gene expression programs using a similar pipeline applied to the transpose of the expression matrix from the previous sections. Initial HDBSCAN clustering on the raw data did not yield very many clusters, which we attributed to the broad range in gene expression program size and dynamic range. To attempt to equalize for these factors, we performed the clustering on a high-dimensional embedding of the data. Each gene was represented by its expression across the 1973 perturbations in Fig. 2 and we masked the targets of knockdown as described there to avoid target gene knockdown influencing clustering. We used pyMDE with the same normalization as above to encourage genes with correlated expression to be placed nearby to each other (20 dimensions, n_neighbors=7 and repulsive_fraction=5, final average distortion 0.145, final residual norm 5.1e-06). We then identified clusters using HDBSCAN applied to the embedding (metric='euclidean', min_cluster_size=10, min_samples=10, cluster_selection_method='leaf'), producing 38 clusters. We performed similar analyses as in Fig. 2 to annotate known CORUM complexes and STRING clusters. Cluster identities were then manually annotated using a combination of these automated annotations, manual gene searches, and gene set enrichment analyses conducted using Enrichr (*71*). These clusters are summarized in Table S8, which includes the fields:

| | |
|---|---|
| **members** | The genes assigned to the cluster by HDBSCAN. |
| **string_cluster_ids** | IDs of any STRING clusters contained within this cluster (at least 75% of members must be represented). |
| **string_clusters** | Descriptions of any STRING clusters contained within this cluster (at least 75% of members must be represented). |
| **corum_complexes** | Names of any CORUM complexes contained within this cluster (at least 75% of members must be represented). |
| **nearby_corum_complexes** | Names of any CORUM complexes that are close to this cluster in the high-dimensional embedding. |
| **nearby_string_clusters** | Names of any STRING clusters that are close to this cluster in the high-dimensional embedding. |
| **strong_positive_clusters** | Perturbation clusters (see Fig. 2) for which average expression of this expression program is at least 2 standard deviations above normal expression level. |
| **strong_negative_clusters** | Perturbation clusters (see Fig. 2) for which average expression of this expression program is at least 2 standard deviations below normal expression level. |
| **manual_annotation** | Manual annotation of cluster function. |
| **example_genes** | Manually curated genes indicative of cluster function. |

To produce Fig. 3B, we averaged expression within the 64 perturbation clusters from Fig. 2 across the genes within the 38 gene expression clusters. Each element in the heat map therefore represents an average over both multiple related perturbations and multiple related genes. The labels were manually selected to highlight interesting features.

**Screens of gene expression programs**

To demonstrate the ability of Perturb-seq to conduct screens on aggregate phenotypes, we conducted two analyses to identify perturbations that strongly induced interesting expression programs. In the first comparison, we compared expression of genes associated with erythroid differentiation (gene expression cluster 15) to those associated with myeloid differentiation (gene expression cluster 21) in Figure 4D. We scored expression programs by taking the mean normalized gene expression of all genes in the associated clusters. We computed scores for all perturbations in the genome-scale dataset that were detected in at least 25 cells, and then $z$-normalized these scores to make scales comparable. The figure has labels on the 15 most outlying genes across the two programs. We then conducted an identical analysis comparing expression of an unfolded protein response cluster (cluster 2) to an integrated stress response cluster (cluster 12) in Figure 4C.

**Analysis of composite phenotypes: total RNA, fraction mtRNA, fraction TE, RNA splicing**

Composite phenotypes integrate data from across the transcriptome to describe global cellular features. While some derive from simple metrics, others rely on extracting information from the transcriptome beyond gene expression levels. As an example of a simple metric, the number of UMIs aligned to the transcriptome (GRCh38 version 2020-A) of single-cells was used to represent the total cellular RNA content. To produce Fig. 4H, we averaged the total RNA content of cells for each perturbation, and z-scored the RNA content with respect to non-targeting controls. Similarly, to calculate the fraction of mitochondrial RNA per cell (fraction mtRNA), the sum of the expression levels of the 13 mitochondrial genome protein-coding genes (MT-ND6, MT-ND1, MT-ND2, MT-ATP8, MT-ND4L, MT-ND5, MT-ND3, MT-CO1, MT-CO2, MT-ND4, MT-ATP6, MT-CO3, MT-CYB) was divided by the total cellular RNA content for each individual cell.

The scTE (*72*) processing pipeline was used to quantify the expression of transposable elements in single cells. As transposable elements tend to be present in many degenerate copies throughout the genome, scTE allocates TE reads to TE metagenes rather than specific genomic positions. Reads were aligned to the genome using STARsolo (STAR version 2.7.9a) with the flags ' --outSAMattributes NH HI AS nM CR CY UR UY --soloFeatures Gene GeneFull SJ Velocyto --readFilesCommand zcat --outFilterMultimapNmax 100 --winAnchorMultimapNmax 100 --outMultimapperOrder Random --runRNGseed 777 --outSAMmultNmax 1' to allow multimapping. To avoid incompatibilities in cell calling between STARsolo and Cell Ranger, the output of Cell Ranger cell calling was used to define the STARsolo cell barcode whitelist using '--soloCBwhitelist'. Next, aligned reads were allocated to genes and TEs using scTE. The flag '-o nointron' was used to prevent the quantification of TEs in gene introns. From single-cell transcriptomes, quantification of all TEs in the classes LINE, SINE, LTR, DNA, and Retroposon based on RepeatMasker were extracted. To calculate the fraction of repetitive and transposable element RNA per cell (fraction TE), the sum of the expression level of these TEs was divided by the total cellular RNA content for each individual cell. To produce Fig. 4G, we averaged the fraction TE of cells for each perturbation.

The alignments from STARsolo described above were also be used to quantify RNA splicing. Due to (i) the sparsity of single-cell data (ii) the relationship between the fraction of spliced reads and gene expression levels, gene-wise RNA splicing was quantified at the pseudobulk level. From the STARsolo Velocyto output, the levels of spliced and unspliced reads for each gene were averaged across all cells bearing each perturbation, ignoring ambiguous reads. Then, for each gene, the fraction of unspliced reads was divided by the mean fraction of unspliced reads for that gene across all non-targeting control perturbations. The results in Fig. 3F display a common set of genes across all perturbations.

**Leverage scores for quantifying perturbation penetrance and variability**

Our use of non-parametric tests in differential expression testing is in part to accommodate perturbations that may be incompletely penetrant or heterogeneous in effect. To attempt to quantify these features, we developed a scalar single-cell score to summarize how outlying each cell's transcriptional state was relative to control cells. We used an approach based on leverage scores, which measure how outlying the rows or columns of a matrix are and which form the basis for many randomized algorithms (*73*). Specifically, we: (i) Construct an expression matrix consisting of all cells that pass the quality filters described in *Internal normalization of gene expression measurements*, and all genes with mean expression >0.25 UMI counts per cell. (ii) To dampen the effects of a handful of strongly induced outlier genes, we clipped any measure with a *z*-score greater than 10 to 10 (this only affects a handful of genes) (iii) To avoid the influence of gemgroup-level batch effects and variable sequencing depth, we then compute leverage scores separately within each gemgroup. Row leverage scores, corresponding to the cell axis of the expression matrix, are calculated as the squared norm of the top 20 left singular vectors within each gemgroup (computed via the truncated SVD routine scipy.sparse.svds with the arpack solver with k=20). We then normalize these scores so that the sum over all cells in the gemgroup is 1 (i.e. compute the leverage sampling probability distribution). (iv) Finally, to integrate leverage scores across gemgroups, we then take logs, and rescale by *z*-normalizing relative to the scores of control cells (subtracting their mean and dividing by their standard deviation). All the leverage scores presented in the figures are the leverage scores after this normalization procedure.

In fig. S10 we conducted various analyses to validate leverage scores as measures of phenotype and to use them to study penetrance of perturbations. We considered all perturbations that passed the following criteria: (i) >5 differentially expressed genes by Anderson-Darling test; (ii) detected in at least 25 cells that passed our quality filters; and (iii) the gene targeted by the perturbation was either undetectable in the expression data or knocked down by at least 30% if detected (i.e., we removed perturbations that appear non-functional). For these perturbations we compared the mean leverage scores to the number of differentially expressed genes found using the Anderson-Darling test. To assess reproducibility we then subset to perturbations that were present in both the genome-scale dataset and the K562 essentials dataset, including non-targeting controls.

In the analyses of perturbations targeting Mediator and the small subunit of the ribosome we only included perturbations that were (i) present in both the genome-scale dataset and the "K562 essentials" dataset and (ii) targeted the principal "P1" transcript identified by the FANTOM consortium. (A handful of genes also had perturbations targeting the P2 transcript that did not generally have effects.) Knockdown was computed as the ratio of mean (unnormalized) expression of the target gene within perturbed cells vs. that in cells with non-targeting sgRNAs. The plots show kernel density estimates of the distributions of the leverage scores of all cells with these perturbations constructed using seaborn's violinplot (with cut set to 0 so that estimated distributions do not extend beyond the range of the data). The gray bars represent the 10%-90% quantiles of cells with non-targeting control sgRNAs for comparison.

Finally, in Fig. 5B we used leverage scores to search for perturbations that had highly variable phenotypes. We considered all perturbations using the same criteria as in fig. S10. We used the standard deviation of the leverage scores as a metric for variability, as diagrammed in Fig. 5A. The two examples in this figure are derived from actual data. The 20 labeled genes are the most outlying from the lowess local regression between the standard deviation of leverage scores and the log of the number of differentially expressed genes detected by the Anderson-Darling test (computed using statsmodels.nonparametric.smoothers_lowess.lowess).

**Analysis of chromosomal instability and cell cycle**

The framework described in inferCNV (*43*) and implemented as infercnvpy (https://github.com/icbi-lab/infercnvpy) was used to detect evidence of chromosomal copy number changes. The raw single-cell gene expression matrix was first filtered to remove lowly expressed genes (<0.05 UMIs per cell across the population), normalized for total UMI content (using scanpy.pp.normalize_total with target_sum=1e6, exclude_highly_expressed=False, and max_fraction=0.05), and log-scaled (using scanpy.pp.log1p). Then we used infercnvpy to compute rolling average gene expression changes for windows of 100 genes with a dynamic threshold of 1.5 standard deviations for noise filtering (using infercnvpy.tl.infercnv). For each single-cell, this generated a vector of CIN values across the genome. To label cells with likely karyotypic abnormalities, unstable karyotypic cells were heuristically defined as having ≥1 chromosome with evidence of changes in chromosomal copy number (nonzero CIN values) for >80% of the chromosomal length. For Fig. 5G, the CIN score of genetic perturbations was calculated as the mean single-cell sum of squared CIN values, *z*-normalized relative to non-targeting control perturbations.

To show cell cycle effects in Fig. 5D,E, we chose to use a dimension reduction approach. Previous approaches to cell cycle analyses in Perturb-seq have largely focused on supervised classification of cells into canonical cell cycle states. However, we found that these approaches did not allow for aberrant cell cycle states sometimes generated by genetic perturbations. As a summary of single-cell cell cycle states, we performed a UMAP dimension reduction based on the expression n=199 known cell cycle genes [obtained from Seurat (*74*) and (*9*)]. From total UMI content normalized, log-scaled expression data, a neighborhood graph was computed (using scanpy.pp.neighbors with n_neighbors=30, method='umap', metric='correlation', and n_pcs=20) followed by UMAP embedding (using scanpy.tl.umap with default parameters). This UMAP revealed cells in canonical cell cycle stages when compared with other methods, but also naturally separated dying cells and putatively quiescent cells. Gates were drawn by manual inspection to approximately separate cells likely to be in S, G2/M, and G1/G0 cell cycle phases.

**Analysis of transcriptional responses to mitochondrial stress**

To compare the functional specificity of nuclear and mitochondrial genome responses to mitochondrial stress, we clustered mitochondrial perturbations based on their transcriptional phenotypes. The analysis presented in Figures 7A and 7C (K562 day 8 data) covers 268 mitochondrial perturbations that met three criteria: (i) at least 50 differentially expressed genes at a significance of $p < 0.05$ by Anderson-Darling test following Benjamini-Hochberg correction; (ii) at least 30 cells; and (iii) an on-target knockdown of at least 60%. As features, we used 1715 genes encoded in the nuclear genome expressed at >1 UMI per cell or the 13 protein-coding mitochondrial-encoded genes. Analogously, the analysis presented in Figures S12A and S12E (RPE1 day 7 data) covers 140 mitochondrial perturbations that met three criteria: (i) at least 20 differentially expressed genes at a significance of $p < 0.05$ by Anderson-Darling test following Benjamini-Hochberg correction; (ii) at least 30; and (iii) an on-target knockdown of at least 60%. As features, we used 2017 genes encoded in the nuclear genome expressed at >1 UMI per cell or the 13 protein-coding mitochondrial-encoded genes.

To cluster these data, we used correlation as a metric to compare z-normalized expression profiles, since it is scale-invariant. We clustered using HDBSCAN (metric='correlation', min_cluster_size=3, min_samples=1, cluster_selection_method='eom', alpha=1.0). As discussed above, this procedure is intrinsically conservative due to the choice of metric and clustering algorithm, so many perturbations are not assigned to any cluster. We used the hierarchical clustering output of HDBSCAN to manually identify and label groups of perturbations in the figures.

To compare the heterogeneity of mitochondrial genome expression, we used two approaches. First, we used a data-driven approach comparing the 38 gene expression programs described in Figure 4B across perturbations. For each program, we scored its expression within each genetic perturbation, and calculated a standard deviation of these scores across the different perturbations in the K562 day 8 experiment. The mitochondrial genome program discovered in this way consists of the 13 protein-coding mitochondrial genes, plus two MT-RNR2-like pseudogenes encoded in the nuclear genome which are likely to multimap with mitochondrial-encoded transcripts. In Figure S12C, we show a histogram of the variability of expression programs across perturbations. Second, we used a hypothesis-driven approach that compared the variation of mitochondrial genome responses by the protein localization of perturbations. We used data from the Human Protein Atlas to assign locations to different perturbations (https://www.proteinatlas.org/about/download table subcellular_location.tsv.zip; some locations were collapsed into supersets), excluding any dual-localized proteins. For all perturbations with the same localization and at least 50 differentially expressed genes at a significance of $p < 0.05$ by Anderson-Darling test following Benjamini-Hochberg correction, we calculated the variance of z-normalized gene expression profiles for each mitochondrially encoded gene. The data in Figure 7B (K562 day 8) and S12B (RPE1) represents the average variance across mitochondrially-encoded genes for each localization.

To quantitatively compare the specificity of the mitochondrial and nuclear transcriptional responses, we employed random forest classifiers. The analysis presented in Figure S12F (K562 day 8 data) covers perturbations that met three criteria: (i) cause at least 50 differentially expressed genes at a significance of $p < 0.05$ by Anderson-Darling test following Benjamini-Hochberg correction; (ii) at least 30 cells; and (iii) belong to either the small or large mitochondrial ribosomal subunit (identified by gene names beginning with MRP), ATP synthase (identified by gene names beginning with ATP5), proteostatic factors (including PAM16, DNAJC19, TFAM, HSPA9, TOMM40, TOMM20, PMPCB, LONP1, and HSPE1). As features, we used 2039 genes encoded in the nuclear genome expressed at >0.5 UMI per cell with variance in the upper 25% of the dataset, or the 13 protein-coding mitochondrial encoded genes. To train the random forest classifier to separate cells by perturbed complex (as described above), we used the scikit-learn implementation with extremely randomized trees with 1,000 trees in the forest and 100 features. The reported accuracy is the balanced accuracy defined as the defined as the average of recall obtained on each class. To visualize the data, we used a UMAP (metric='correlation', n_neighbors=10) generated on the nuclear or mitochondrial genome.

**Table of antibodies**

| **Target** | **Antibody** | **Dilution** |
|---|---|---|
| α-INTS10 | Abcam ab180934 (Rabbit mAb) | 1:1,000 |
| α-INTS13 | Bethyl A303-575A (Rabbit pAb) | 1:1,000 |
| α-INTS14 | Prestige HPA040651 (Rabbit pAb) | 1:1,000 |
| α-C7orf26 | Prestige HPA052175 (Rabbit pAb) | 1:1,000 |
| α-His | Cell Signaling Technologies 2366 (Mouse mAb) | 1:2,000 |
| α-Mouse | Licor 926-32210 (Goat pAb) | 1:10,000 |
| α-Rabbit | Licor 926-32213 (Donkey pAb) | 1:10,000 |
| α-CD11b | Biolegend 101220 (M1/70, AF647) | 1:50 |

**Table of cell lines**

| **Cell line** | **Source** |
|---|---|
| K562 dCas9-BFP-KRAB | Gilbert, 2014 |
| RPE1 Zim3-dCas9-2A-BFP | This study |
| K562 Zim3-dCas9-2A-BFP | This study |

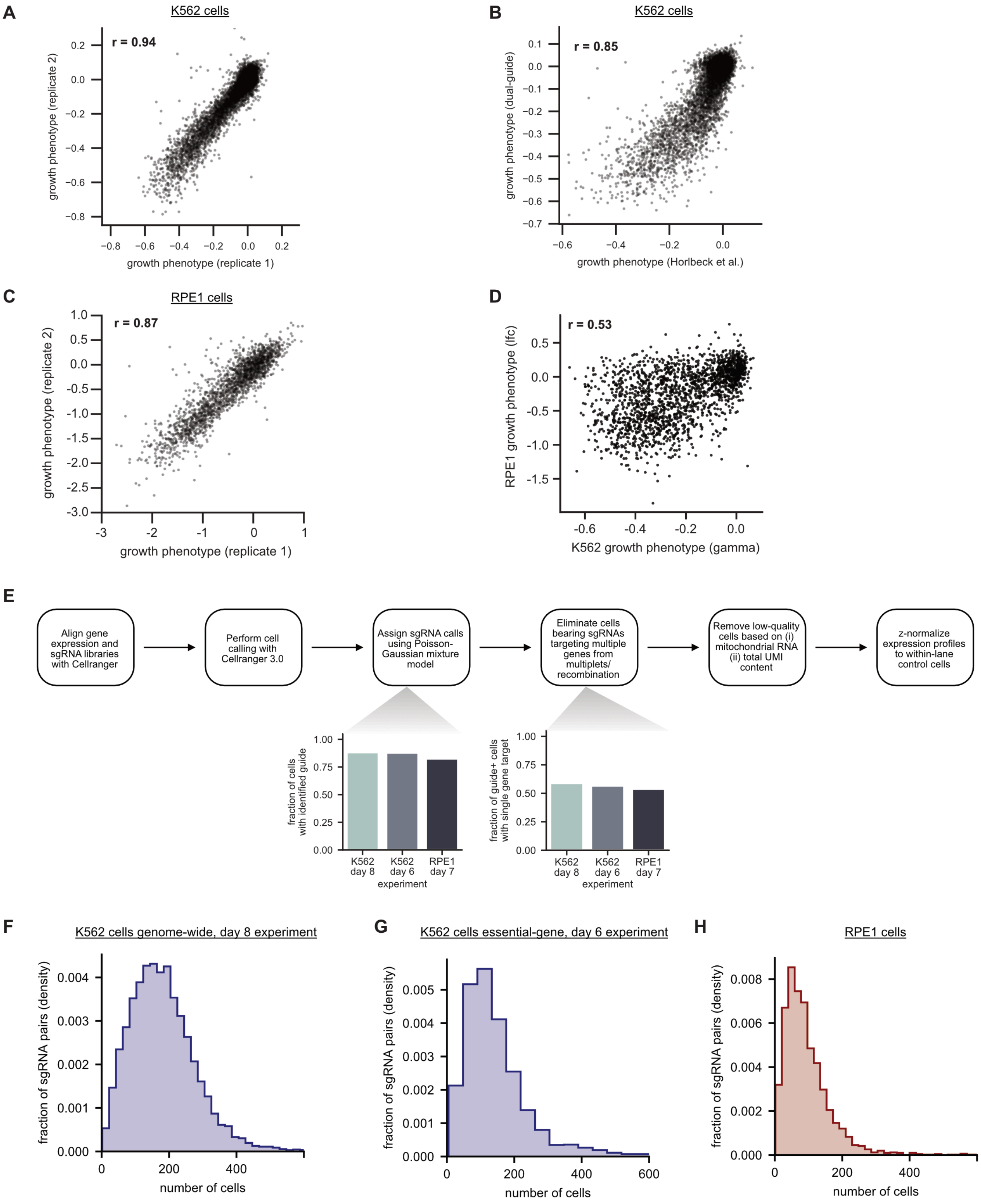


**Figure S1. (legend on next page)**

**Figure S1: Growth screens, filtering, and coverage.**

A) Comparing the growth phenotypes of dual-sgRNA constructs between growth screen replicates in K562 cells. Growth phenotypes are reported as the $\log_2$ guide enrichment per cell doubling (gamma) between day 6 and day 16 post library transduction. Replicates are strongly correlated (n=11,056 dual sgRNA constructs; r = 0.94). For 50 outlier genes (where the residual from a regression comparing replicates was >0.2), the growth phenotype was set to missing.

B) Benchmarking the growth phenotypes of dual-sgRNA constructs to single-sgRNA screens. Growth phenotypes (gammas) are compared between the dual-sgRNA library compared to the mean of the best three sgRNAs from Horlbeck et al. The screens are strongly correlated (n=9386 genes after excluding constructs mapping to secondary TSSs; r = 0.85) but with stronger growth phenotypes observed for the dual-sgRNA library.

C) Comparing the growth phenotypes of dual-sgRNA constructs between growth screen replicates in RPE1 cells. Growth phenotypes are reported as the $\log_2$ guide enrichment between the plasmid library and day 7 post library transduction. Replicates are strongly correlated (n=2203 constructs targeting common essential genes; r = 0.87). For 19 outlier genes (where the residual from a regression comparing replicates was >1), the growth phenotype was set to missing.

D) Comparing growth phenotypes between K562 and RPE1 cells. Growth phenotypes are correlated (n=1951 constructs; r=0.53) despite substantial differences in screen timepoint (day 6 to day 16 for K562 cells versus day 0 to day 7 in RPE1 cells).

E) Schematic overview of data alignment, cell calling, sgRNA assignment, and filtering.

F) Histogram of the number of cells per genetic perturbation in the K562 day 8 genome-wide Perturb-seq experiment. The number of detected genetic perturbations (expected sgRNA pairs) was n=11,258, with a mean coverage 183 cells per perturbation and a median coverage of 171 cells per perturbation after filtering.

G) Histogram of the number of cells per genetic perturbation in the K562 day 6 essential-wide Perturb-seq experiment. The number of detected genetic perturbations (expected sgRNA pairs) was n=2,285, with a mean coverage 148 cells per perturbation and a median coverage of 124 cells per perturbation after filtering.

H) Histogram of the number of cells per genetic perturbation in the RPE1 cell day 7 essential-wide Perturb-seq experiment. The number of detected genetic perturbations (expected sgRNA pairs) was n=2,679, with a mean coverage 101 cells per perturbation and a median coverage of 79 cells per perturbation after filtering.

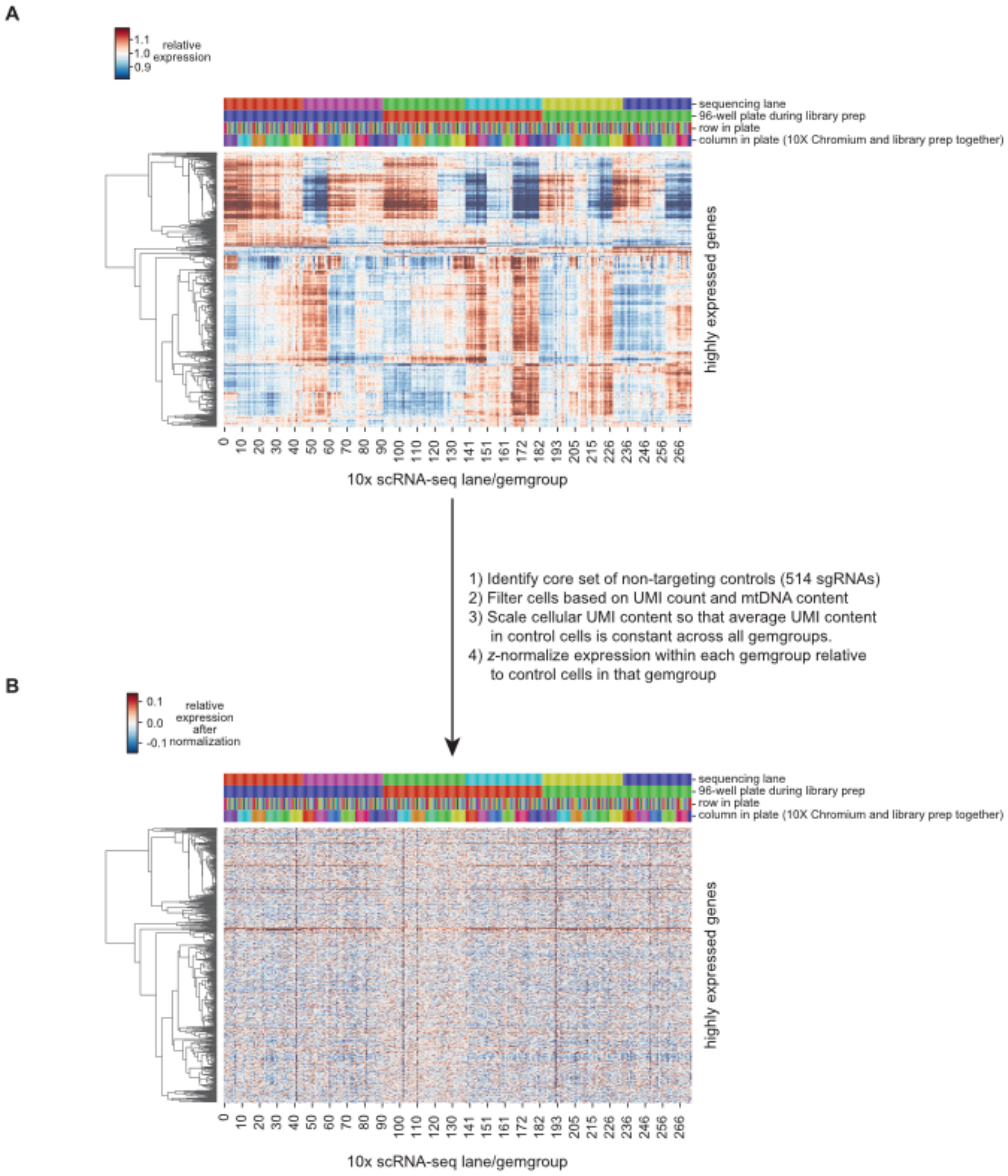


**Figure S2: Schematic and performance of internal normalization of gene expression measurements**

A) Batch effects in raw data. The K562 day 8 genome-wide experiment was conducted across 273 separate lanes of 10x Genomics droplet single-cell RNA sequencing ("gemgroups"). The plot shows mean expression profiles of highly expressed genes (>2 UMI per cell) within all cells in each gemgroup. The data is normalized (i) for sequencing depth of each gemgroup and (ii) so that the mean expression of each gene is 1 across all gemgroups. Colors at the top indicate different levels of multiplexing that were present within the experiment: including groups of samples (generally in sets of 8) that went through scRNA-seq together, 96 well plates used for library preparation, and separate lanes used during Illumina sequencing. The range of the heatmap is set according to the 2%-98% quantiles of the data.

B) Expression following internal normalization. We rescale gene expression by *z*-normalizing relative to the control cells (containing non-targeting sgRNAs, ~4% of all cells) within the same gemgroup. The plot shows the average normalized expression of all cells within each gemgroup following this procedure, with the range of the heatmap set according to the 2%-98% quantiles of the data. By construction control cells have mean expression 0 and standard deviation 1 in this scale. Genes are presented in the same order as in panel (A).

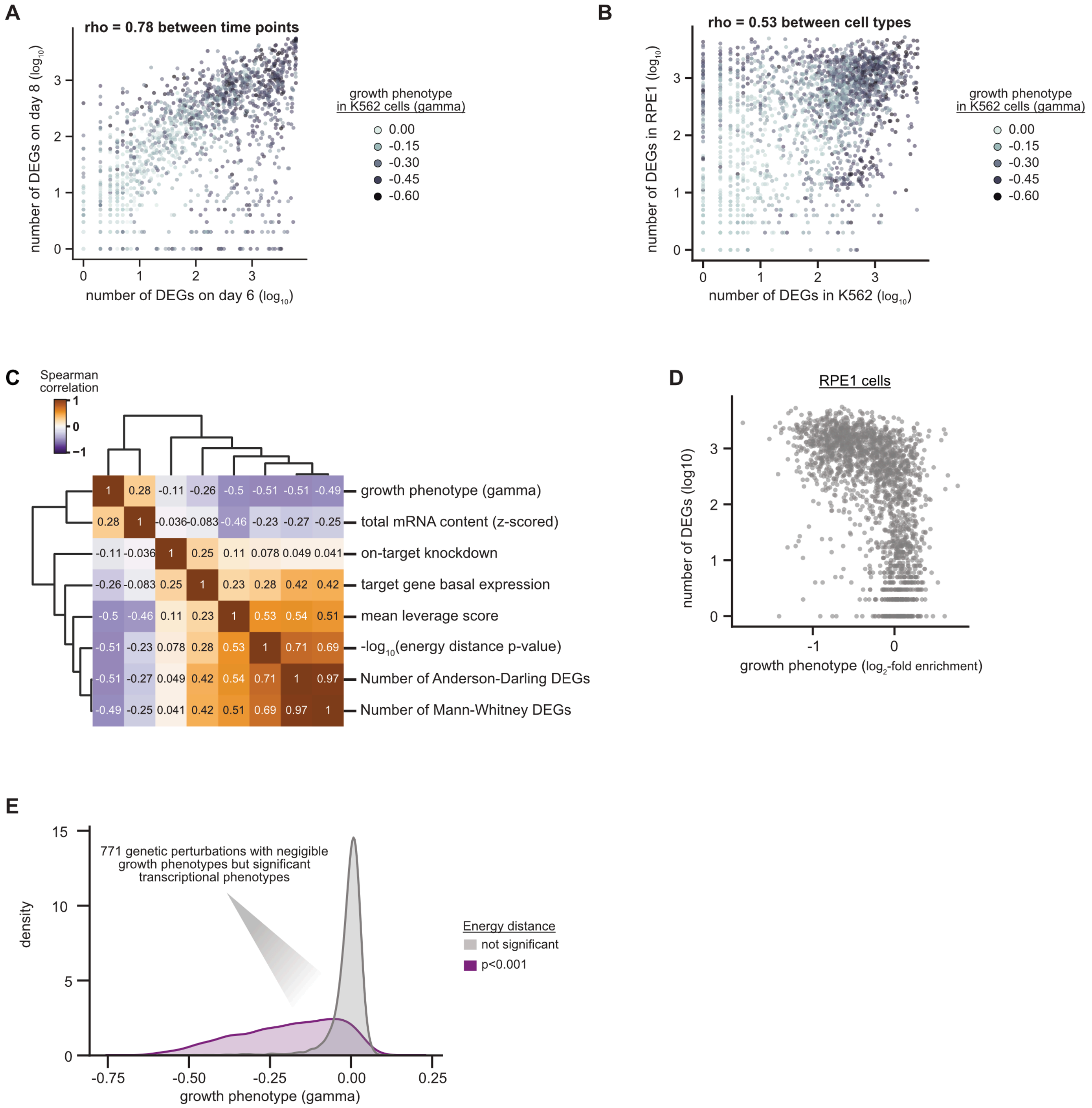


**Figure S3: Growth screens, filtering, and coverage.**

A) Relationship between the number of differentially expressed genes (DEGs) for a genetic perturbation in K562 cells at day 8 versus day 6 post-transduction. DEGs were determined using a two-sample Anderson-Darling test comparing against non-targeting guides (n=2276 common genetic perturbations, Spearman's rho=0.78).

B) Relationship between the number of DEGs for a genetic perturbation in K562 cells (day 8 genome-wide dataset) versus RPE1 cells. DEGs were determined using a two-sample Anderson-Darling test comparing against non-targeting guides (n=2636 common genetic perturbations, Spearman's rho=0.53).

C) Relationship between features of genetic perturbations in K562 cells genome-wide day 8 Perturb-seq. The features were calculated as detailed in Methods. The heatmap displays Spearman correlations between features.
D) Comparing the growth phenotype versus the number of DEGs for each multiplexed guide pairs in RPE1 cells. Growth phenotypes are reported as the $log_2$ guide enrichment between day 0 and day 7 post-lentiviral transduction. DEGs were determined using a two-sample Anderson-Darling test comparing against non-targeting guides.
E) The distribution of growth phenotypes in genetic perturbations with a transcriptional phenotypes in K562 cells genome-wide day 8 Perturb-seq. Histogram (kernel density estimate) comparing the growth phenotype in K562 cells (gamma) of genetic perturbations to the permuted energy distance test. 771 genetic perturbations had a gamma > -0.1 (considered a negligible effect on cellular growth) but a significant transcriptional phenotype.

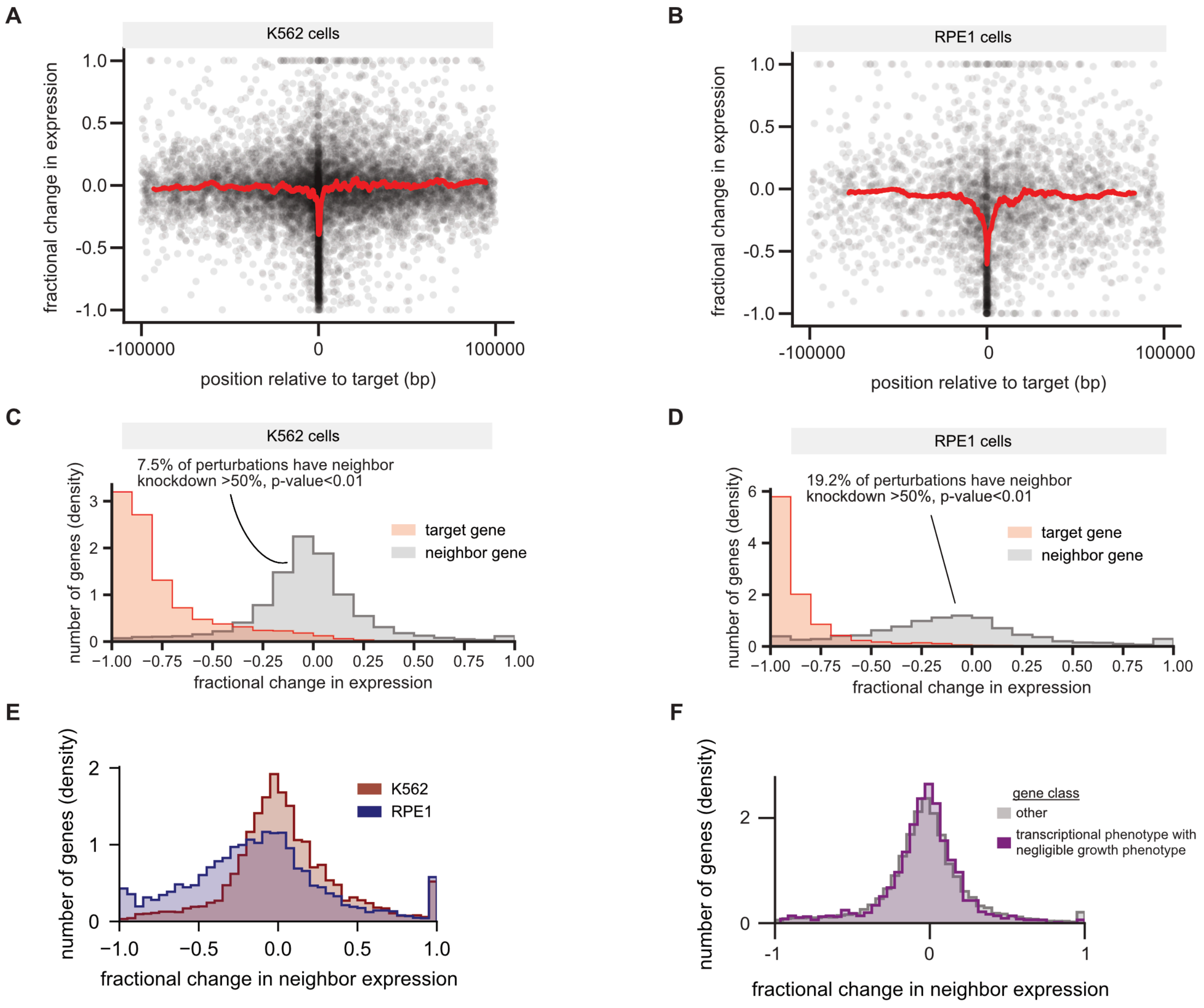


**Figure S4: Assessing neighbor gene off-target knockdown in Perturb-seq data.**

A) and B) Relationship between neighbor gene off-target knockdown and position relative to the target gene in K562 cells (day 8) (A) and RPE1 cells (B). For each target genes, the two neighbor genes are defined as the gene immediately upstream and downstream (at an expression >0.1 UMI per cell). The position relative to the target is the distance of either the start or end of the neighbor gene (whichever is closer) to the start of the target gene. The fractional change in expression is defined as the expression in the targeted cells minus the expression in non-targeting cells, relative to the expression in the non-targeting cell population (-1 implies 100% knockdown).

C) and D) Comparison between target gene and neighbor gene knockdown in K562 cells (day 8) (C) and RPE1 cells (D). P-values are assigned by comparing a bootstrap test.

E) Comparison of neighbor gene knockdown in K562 cells (day 8) versus RPE1 cells.

F) Comparison of neighbor gene knockdown based on transcriptional phenotype in K562 cells (day 8). Perturbations with "transcriptional phenotype with negligible growth phenotype" are those perturbations where gamma > -0.1 that had a significant transcriptional phenotype by the permuted energy distance test.

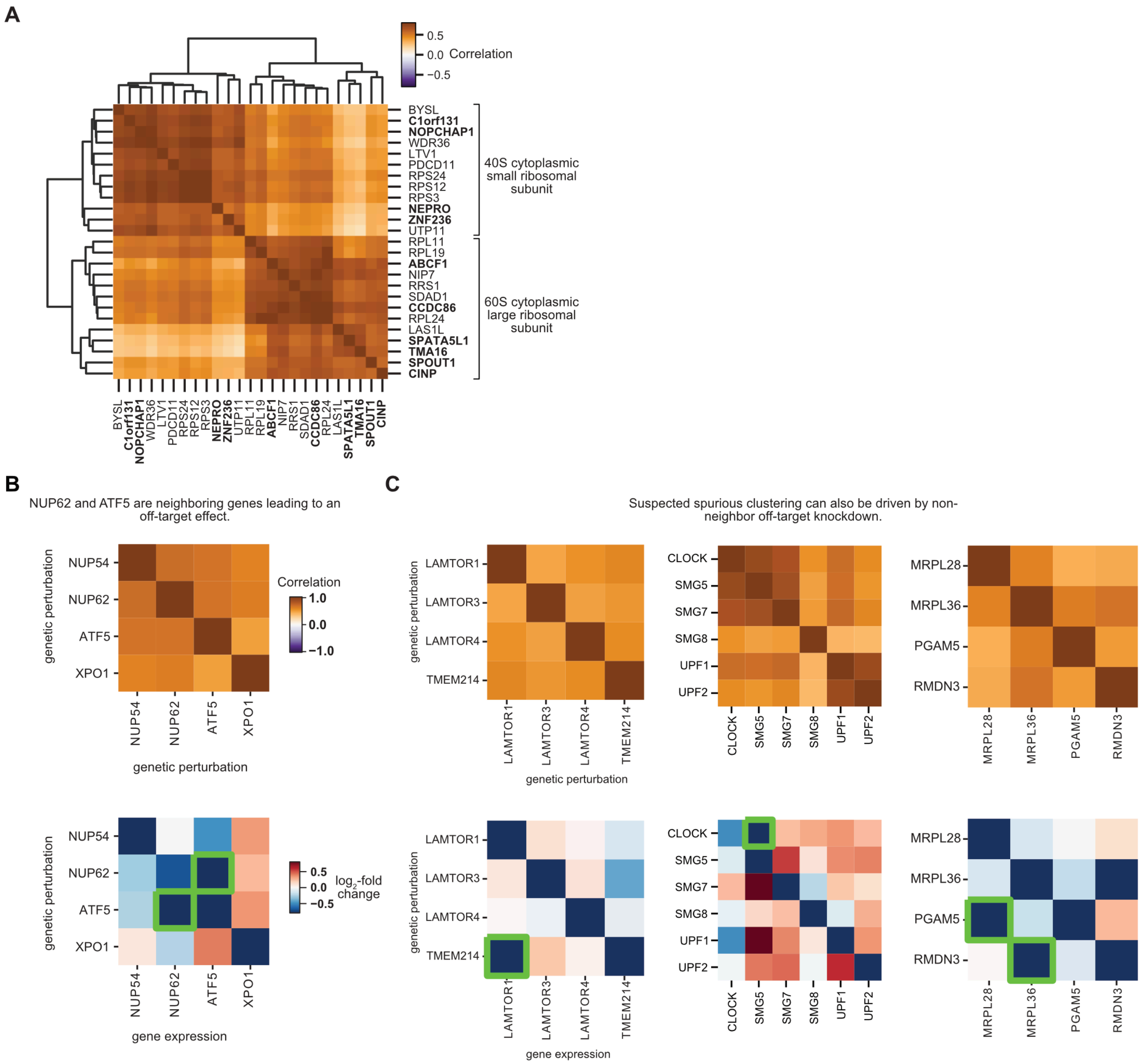


**Figure S5: Supplementary data related to defining gene function with Perturb-seq.**

A) Relationship between members of ribosomal subunits, biogenesis factors, and poorly characterized genes. The heatmap displays the Pearson correlation between pseudobulk $z$-normalized gene expression profiles of select genes. Genetic perturbations are ordered by average linkage hierarchical clustering based on Euclidean distance in K562 cells (day 8). Poorly characterized genes are shown in bold.

B) Investigation of *ATF5* phenotype in K562 cells (day 8). The upper heatmap shows the correlation of the expression profile of *ATF5* knockdown with similar genetic perturbations. The sgRNA pair targeting *ATF5* leads to a phenotype strongly correlated with knockdown of subunits of the nuclear pore complex (*NUP54*, *NUP62*) and nuclear export proteins (*XPO1*). The sgRNA pair targeting *ATF5* leads to strong downregulation of NUP62 (bottom heatmap, shown as the $\log_2$-fold change compared to control cells). *ATF5* is bidirectionally expressed with *NUP62* on chromosome 19, explaining this neighbor gene off-target knockdown and similar phenotypes.

C) Investigation of other surprising phenotypes in K562 cells (day 8). We investigated three different surprising relationships between genetic perturbations: clustering of *TMEM215* with LAMTOR subunits, clustering of *CLOCK* with NMD machinery, and clustering of *PGAM5* and *RMDN3* with the mitochondrial large ribosomal subunit (upper heatmaps). In all three cases, the relationship could be explained by suspected off-target knockdown of a component of the complex, directly detected in Perturb-seq (bottom heatmaps).

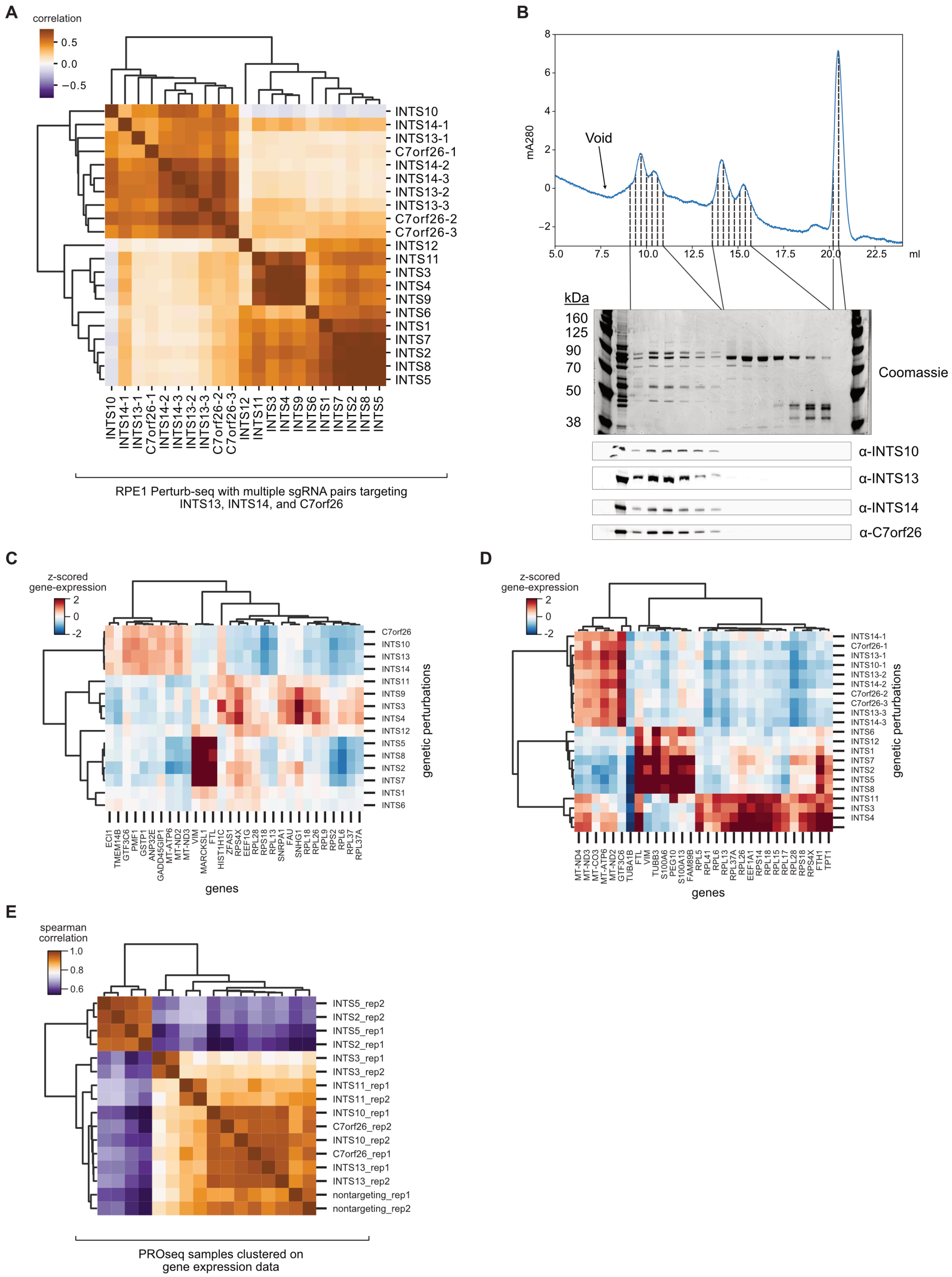


**Figure S6. (legend on next page)**

**Figure S6: Supplementary data related to the functional modules of the Integrator complex.**

A) Relationship between Integrator complex members and *C7orf26* in RPE1 cells. Multiple independent sgRNA pairs were used to target *INTS13*, *INTS14*, and *C7orf26*, with independent guides indicated by numbers next to gene names (e.g. C7orf26-1, C7orf26-2, etc.). The heatmap displays the Pearson correlation between pseudobulk *z*-normalized gene expression profiles of Integrator complex members. Genetic perturbations are ordered by average linkage hierarchical clustering based on correlation.

B) SEC trace and full Western blots for purification of a INTS10-INTS13-INTS14-C7orf26 complex. His-INTS10, INTS13, INTS14, and C7orf26 were overexpressed in Expi293 cells, affinity purified, and separated via SEC. The INTS10-INTS13-INTS14-C7orf26 proteins co-fractionated as a higher molecular weight species as visualized by Western blotting.

C) and D) Identification of differentially expressed genes between Integrator complex modules in K562 cells (C) and RPE1 cells (D). Random forest classifiers were trained on gene expression profiles to classify cells as having perturbation to one of three Integrator complex modules ("endonuclease", "shoulder and backbone" or "10-13-14-C7orf26"). The top 30 gene features led to an accuracy of 97% in K562 cells and 89% in RPE1 cells. Heatmap displays the z-scored gene expression of the top 30 features in each cell type.

E) Comparison of PRO-seq active RNA polymerase II gene expression profiles across genetic perturbations. PRO-seq reads were aligned to the transcriptome and depth normalized. Heatmap shows the Spearman correlation of expression profiles with biological replicates indicated (e.g. INTS5_rep1 and INTS5_rep2).

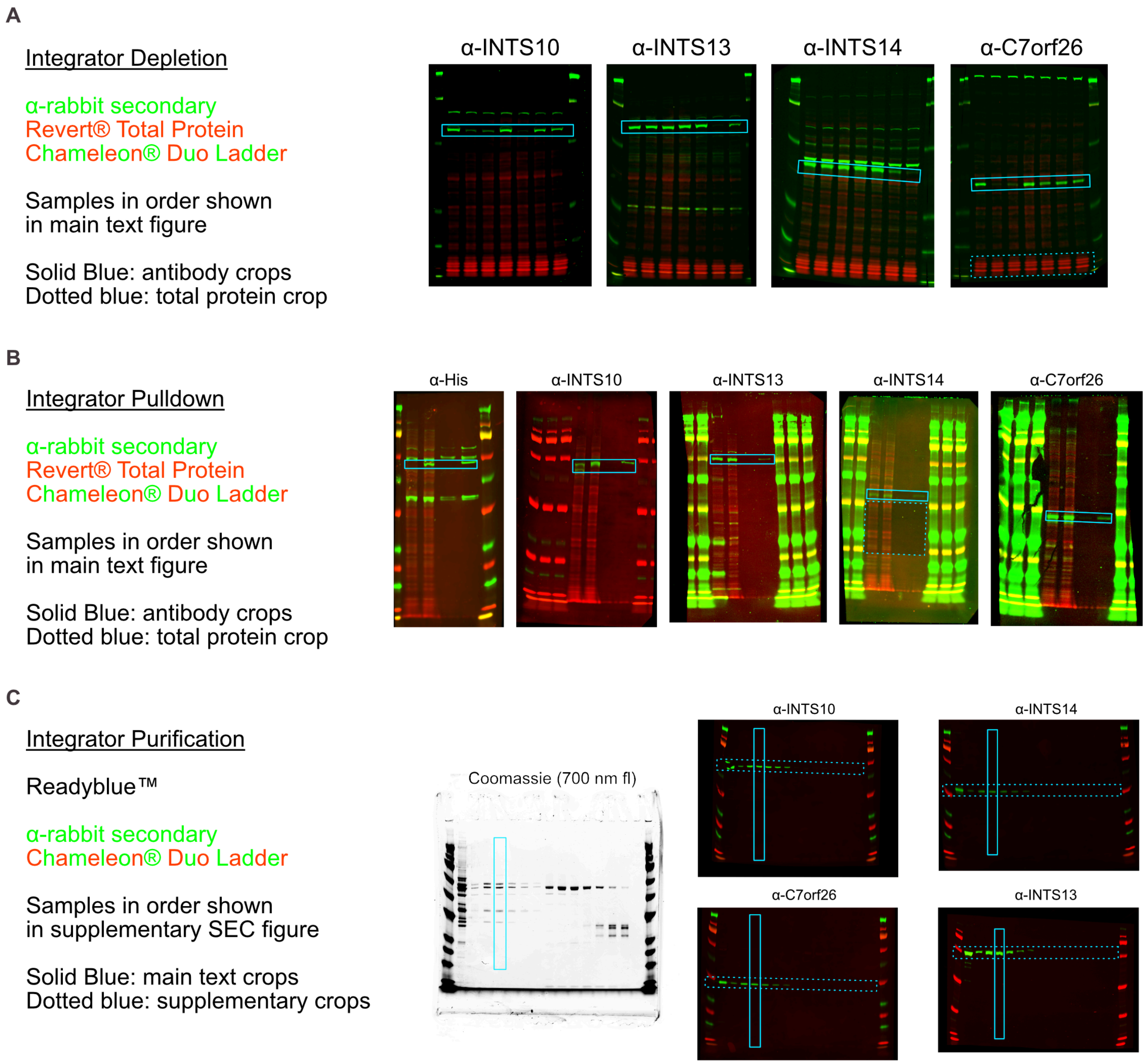


**Figure S7: Supplementary data related to Integrator biochemistry.**

A) Full blots visualizing the effects of CRISPRi-based depletion of Integrator subunits with different probes.
B) Full blots visualizing INTS10 pulldown with different probes.
C) Full blots visualizing Integrator SEC purification with different probes.

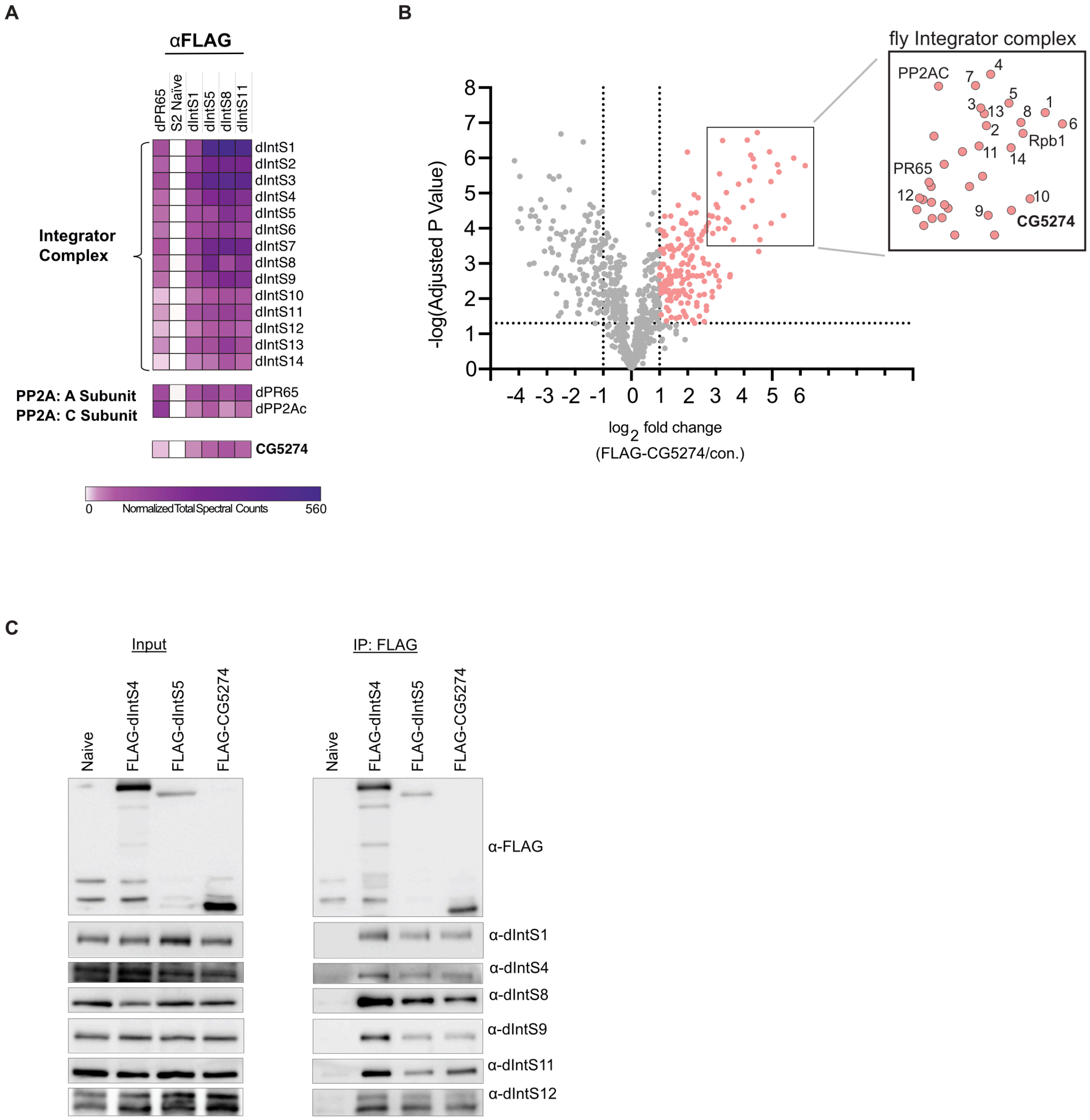


**Figure S8: Supplementary data related to Integrator biochemistry in Drosophila.**

A) Heatmap of results from co-immunoprecipitation of Integrator complex components in Drosophila. Heatmaps were generated using average spectral counts resulting from triplicate pulldowns that were each normalized to total spectra.

B) Volcano plot of enrichments from co-immunoprecipitation using nuclear extract derived from S2 cells stably expressing FLAG-tagged CG5274, which is the Drosophila orthologue of *C7orf26*. Enrichment was calculated using spectral counts and significance was determined from three independent purifications.

C) Co-immunoprecipitation of Integrator complex components in Drosophila. Cell lysates were affinity purified and select Integrator proteins were probed by western blot.

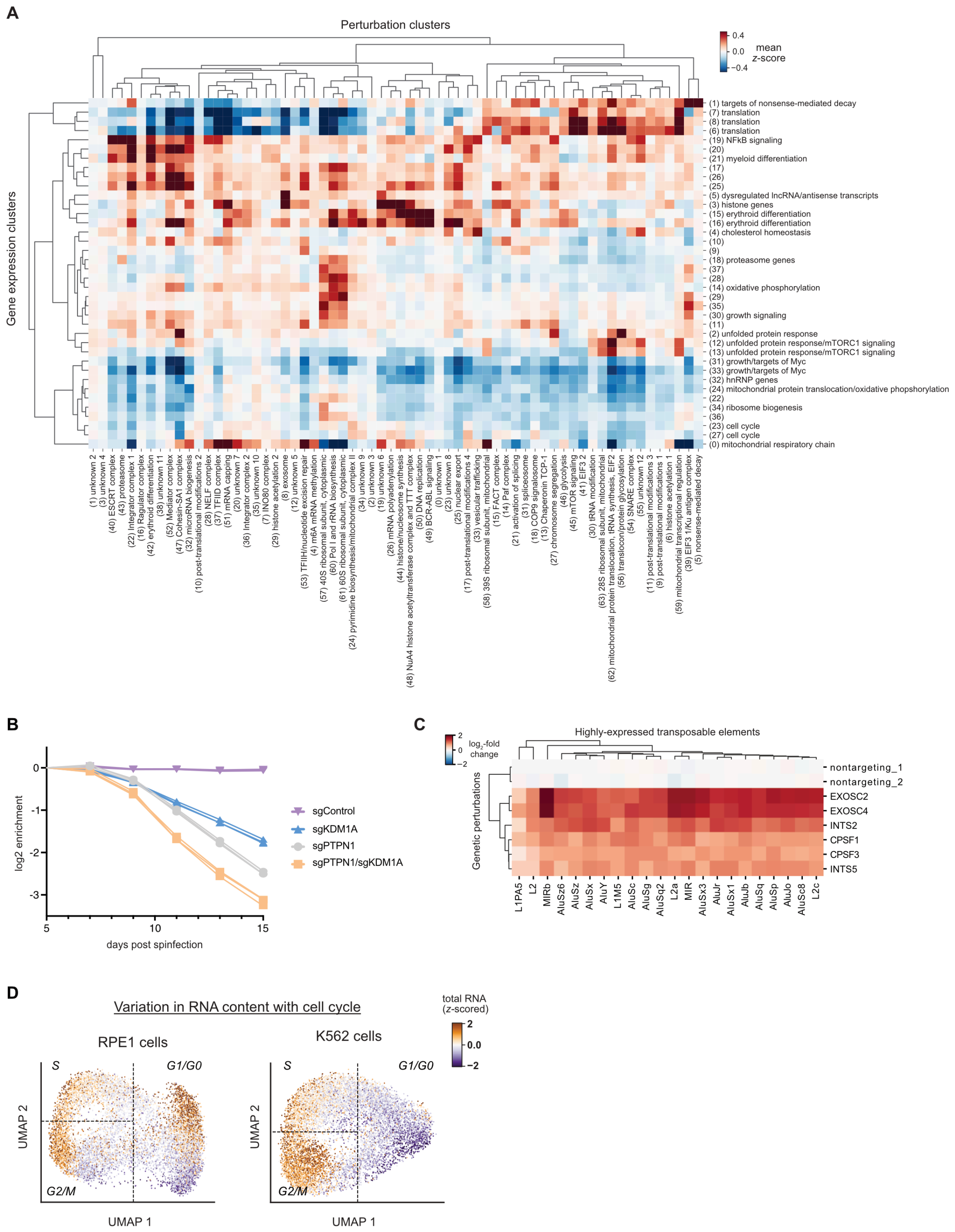


**Figure S9: Supplementary data related to phenotype relationships.**

A) Heatmap of the high-level genotype-phenotype map (identical to Figure 4B with full labels). The heatmap represents the mean z-scored expression for gene expression and perturbation clusters. For a subset of clusters, clustered are labelled with manual annotations (black labels) of cluster function.

B) Growth effect of *PTPN1* or *KDM1A* knockdown in K562 cells. Cells were co-transduced with fluorescently labelled sgKDM1A, sgPTPN1, or a non-targeting control guide. Enrichment was determined by flow cytometry relative to uninfected cells in biological triplicate.

C) Comparison of transposable element expression profiles between top regulators. Heatmap displays $\log_2$-fold changes in expression of highly-expressed transposable element metagenes (columns) for genetic perturbations (rows) in K562 cells (day 8) Perturb-seq. Genetic perturbations and genes are ordered by average linkage hierarchical clustering with a Euclidean distance metric.

D) Comparison of total RNA content with cell cycle state. For single-cells, cell-cycle positioning was inferred by UMAP dimension reduction on differential expression profiles of 199 selected cell-cycle regulated genes. The dimension reduction was performed independently for RPE1 cells (left) and K562 cells (right). Cell cycle occupancy is shown as a scatterplot of UMAP positions of a random subset of 10,000 cells per cell type. Approximate gates between cell cycle phases (G1 or G0; S; G2 or M) are shown as dotted lines. The total RNA content per cell was calculated from the total number of UMIs detected per cell which were z-scored with respect to gemgroup/lane control cells.

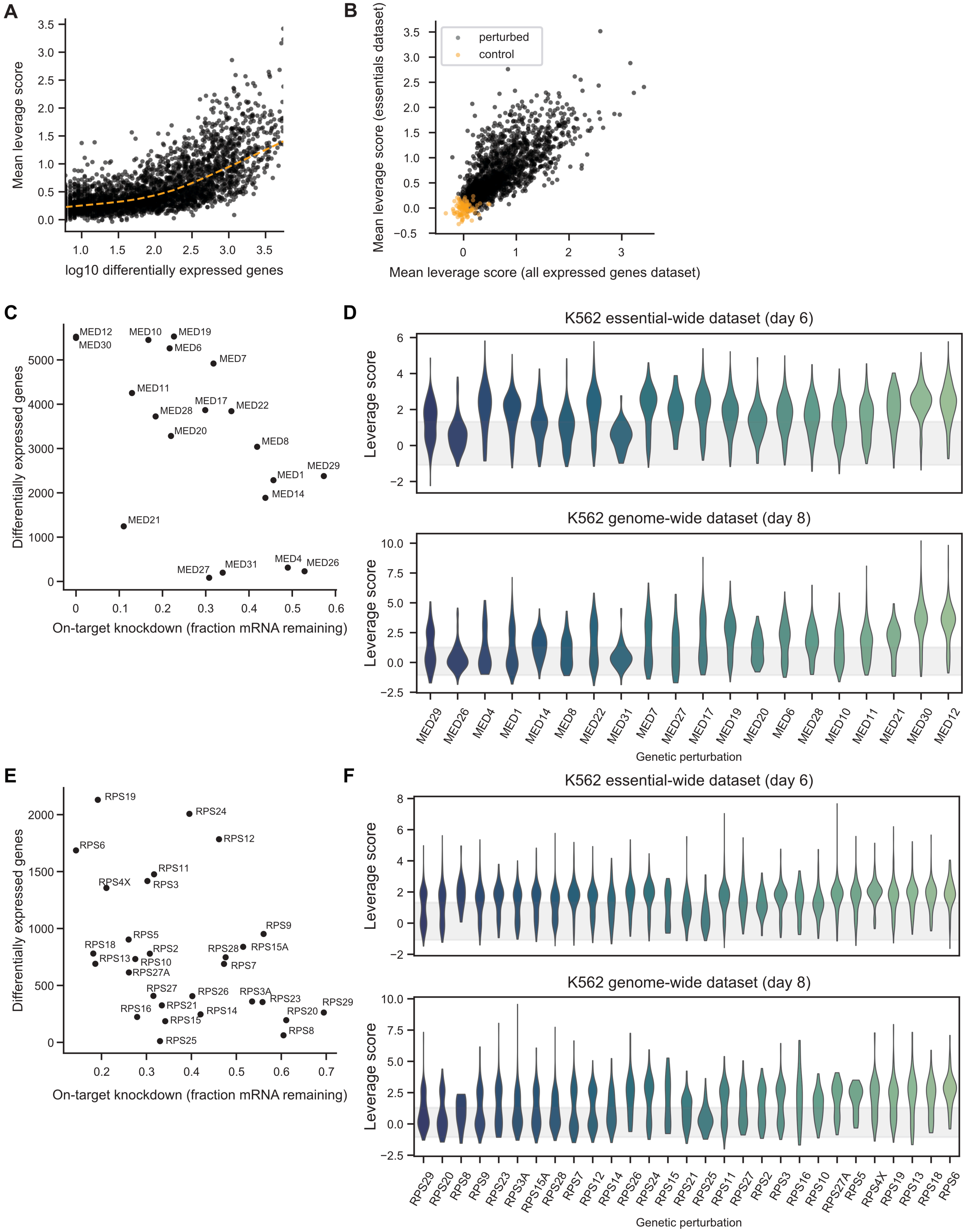


**Figure S10. (legend on next page)**

**Figure S10: Assessing the penetrance and heterogeneity of response to genetic perturbations.**

A) We scored how outlying each perturbed cell was relative to non-targeting control cells using leverage scores. The plot compares the mean leverage score for each genetic perturbation to the number of differentially expressed genes detected by the Anderson-Darling test (Spearman's rho = 0.71).

B) To assess reproducibility of leverage scores, plot compares mean leverage scores of perturbations (black dots) in K562 cells between the essentials dataset (taken at day 6 post-infection) and the dataset targeting all expressed genes (taken at day 8 post-infection). Non-targeting control sgRNAs are in orange (Spearman's rho = 0.79).

C) Relationship between knockdown of target gene (relative to expression in control cells bearing non-targeting sgRNAs) and number of differentially expressed genes detected by Anderson-Darling test for perturbations targeting subunits of the Mediator complex.

D) Leverage scores distributions of perturbations targeting subunits of the Mediator complex. Plot shows kernel density estimates for each perturbation ordered from least knocked down (left) to most (right). Top panel is within essentials dataset and bottom panel is within the all expressed genes dataset. The gray bar shows the 10%-90% range of leverage scores within control cells bearing non-targeting sgRNAs.

E) and F) As in C) and D) but for perturbations targeting the small subunit of the ribosome.

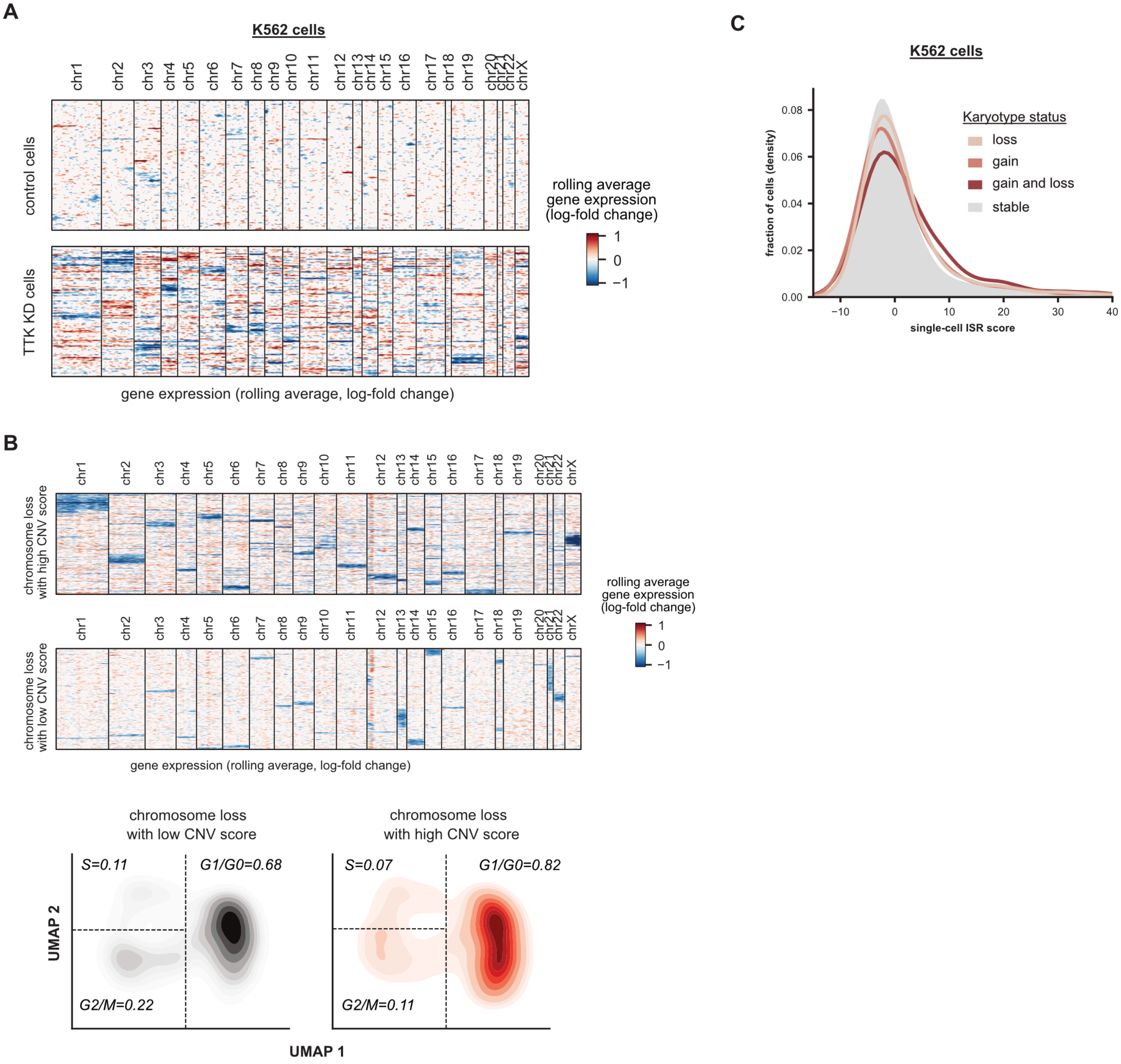


**Figure S11: Supplementary data related to chromosomal instability.**

A) Heatmap of chromosomal copy number inference from Perturb-seq data. For all genes (expressed >0.05 UMI per cell), the log-fold change in expression is calculated with respect to the average of non-targeting control cells, and genes are ordered along the genome. A weighted moving average of 100 genes is used infer copy number changes (columns) in single-cells (rows) with noise and median filtering. 199 *TTK* knockdown K562 cells and 199 randomly sampled non-targeting control K562 cells are shown (data from K562 essential-wide day 6 dataset). Cells are ordered by average linkage hierarchical clustering based on correlation of chromosomal copy number profiles.

B) Comparison of cell cycle effects by magnitude of karyotypic abnormality in RPE1 cells. RPE1 cells with at least one chromosomal loss (defined as evidence of chromosomal loss for >80% of the chromosomal length) were stratified into high, medium, and low degree of karyotypic abnormality based on their CNV score. 500 randomly-sampled high and low CNV score cells were visualized in a heatmap of chromosomal copy number inference. Below, the cell cycle occupancy of high and low CNV cells is shown for 1000 randomly-sampled cells. For single-cells, cell-cycle positioning was inferred by UMAP dimension reduction on differential expression profiles of 199 selected cell-cycle regulated genes. Cell cycle occupancy is shown as a 2D kernel density estimate of a random subset of 1000 cells per karyotypic status. Approximate gates between cell cycle phases (G1 or G0; S; G2 or M) are shown as dotted lines, and the fraction of cells in each cell cycle phase are indicated.

C) Effect of chromosomal instability (CIN) on activation of the Integrated Stress Response (ISR). Histogram (kernel density estimate) compares the ISR score versus CIN status in K562 cells (day 6). CIN status is defined as evidence of gain or loss of chromosomal copy number for >80% of the chromosomal length, with 290,432 stable cells, 11,100 cells bearing chromosomal loss, 5,852 cells bearing chromosomal gain, and 4,541 cells bearing gain and loss of chromosomes. ISR score is defined as the sum of z-normalized expression of ISR marker genes where increased values indicate stronger ISR activation.

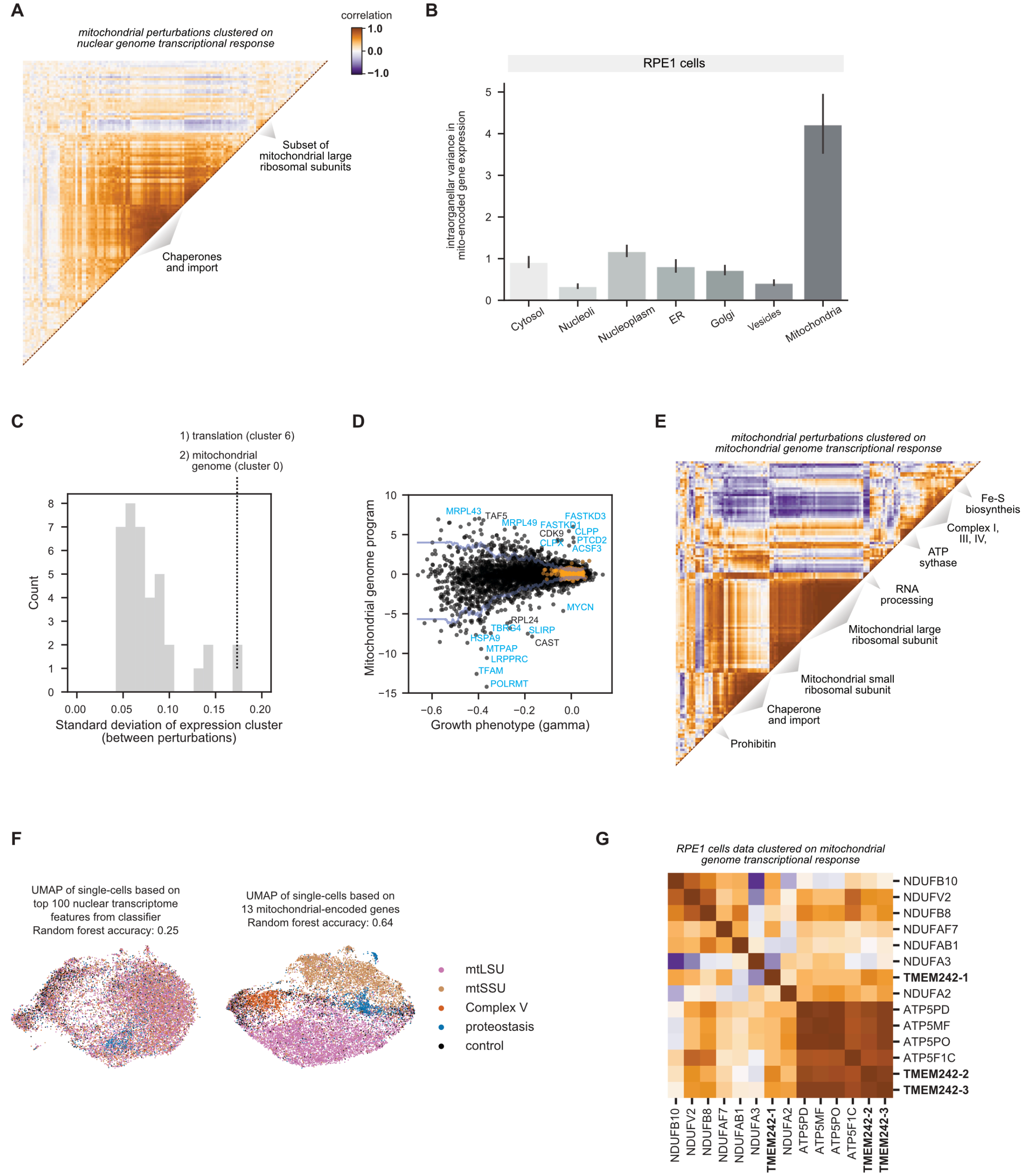


**Figure S12: Supplementary data related to mitochondrial genome regulation.**

A) Clustering mitochondrial perturbations by nuclear transcriptional response. CRISPRi enables knockdown of nuclear-encoded genes whose protein products are targeted to mitochondria (mitochondrial perturbations). Mitochondrial perturbations were annotated by MitoCarta3.0 and subset to those with a strong transcriptional phenotype (n=140 mitochondrial perturbations). Gene expression profiles were restricted to nuclear encoded genes (including 99% of mitochondrial proteins). The heatmap displays the Pearson correlation between pseudobulk *z*-normalized gene expression profiles of mitochondrial perturbations in RPE1 cells. Genetic perturbations are ordered by HDBSCAN with a correlation metric.

B) Comparing variability in the mitochondrial transcriptome by perturbation localization. The mitochondrial genome encodes 13 protein-coding genes. Genetic perturbations were grouped based on localization of their protein products as determined by the Human Protein Atlas. For each of these 13 mitochondrially encoded genes, the variance in pseudobulk z-normalized expression profiles was calculated between all perturbations with the same localization. Barplots represent the average across genes with 95% confidence interval obtained by bootstrapping.

C) Variability of gene expression programs from Figure 4B across perturbations. 38 clusters of co-regulated genes were defined via HDBSCAN clustering, and scored within each perturbation. The histogram shows the standard deviation of these scores across the different perturbations in the K562 day 8 experiment.

D) Activity of mitochondrial genome program in different perturbations. The plot compares growth phenotypes of each perturbation (black dots) to the scores of the mitochondrial genome program in the K562 day 8 experiment. The mitochondrial genome program consists of the 13 protein-coding mitochondrial genes, plus two MT-RNR2-like pseudogenes encoded in the nuclear genome. Negative/positive scores indicate up-/downregulation of the program relative to cells with non-targeting sgRNAs (orange dots). Blue lines indicate local estimate of the +/- 2 standard deviations range of the dataset. Labels indicate the most outlying perturbations, with cyan labels indicating genes with known functions in mitochondria.

E) Clustering mitochondrial perturbations by mitochondrial transcriptional response. Mitochondrial perturbations were annotated by MitoCarta3.0 and subset to those with a strong transcriptional phenotype as above (n=140 mitochondrial perturbations). Gene expression profiles were restricted to the 13 mitochondrial-encoded genes. The heatmap displays the Pearson correlation between pseudobulk *z*-normalized gene expression profiles of mitochondrial perturbations in RPE1 cells. Genetic perturbations are ordered by HDBSCAN with a correlation metric.

F) Comparison of predictive accuracy of nuclear versus mitochondrial genome response. To assess the specificity of the nuclear and mitochondrial genome regulation, random forest classifiers were trained on gene expression profiles to classify cells as having perturbation to one of four mitochondrial complexes ("mtLSU" which corresponds to components of the mitochondrial large ribosomal subunit; "mtSSU" which corresponds to components of the mitochondrial small ribosomal subunit; "Complex V" which corresponds to components of ATP synthase; "proteostasis" which corresponds to essential proteostatic machinery; "control" which corresponds to non-targeting control cells). In K562 cells (day 8), the top 100 gene features led to an accuracy of 25% for the nuclear-encoded genes, versus 64% for the 13 mitochondrial-encoded genes. As a visual guide, the plot displays the UMAP embedding of single cells colored by perturbed mitochondrial complex based on sgRNA assignment.

G) Clustering of *TMEM242* genetic perturbation based on the mitochondrial transcriptome. Genetic perturbations to members of ATP synthase and Complex I of the respiratory chain were compared to knockdown of *TMEM242*, a mitochondrial gene of unknown function. Gene expression profiles were restricted to the 13 mitochondrially encoded genes. The heatmap displays the Pearson correlation between pseudobulk *z*-normalized gene expression profiles of mitochondrial perturbations in RPE1 cells. Multiple independent sgRNA pairs targeting *TMEM242* were used. Genetic perturbations are ordered by HDBSCAN with a correlation metric.

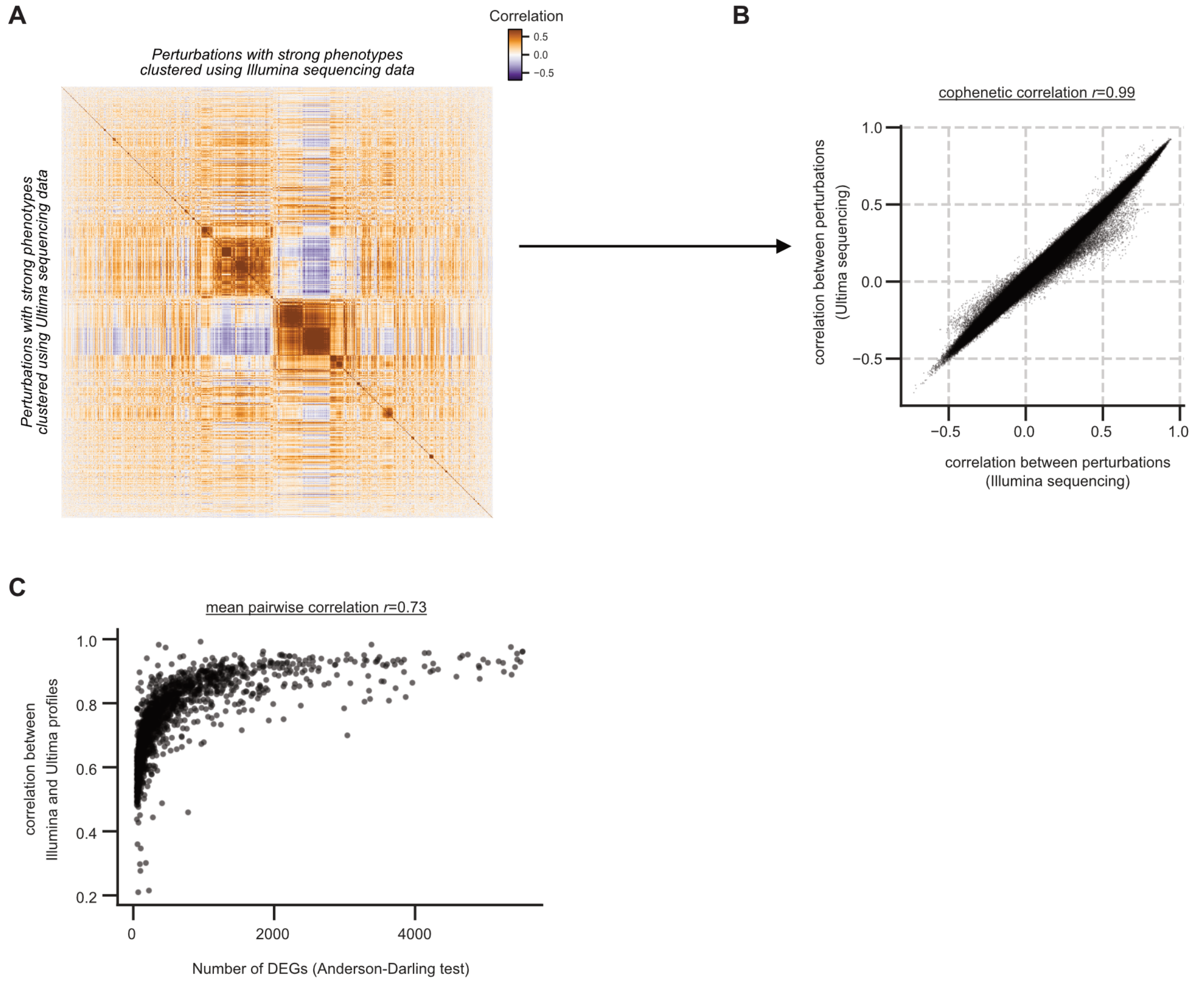


**Figure S13: Validation of Ultima Genomics ultra-high throughput sequencing.**

A) Heatmap of perturbation relationships as derived from independent Ultima and Illumina sequencing data. We analyzed 2,017 genetic perturbations that elicited strong phenotypes. Pearson correlations were used to summarize perturbation-perturbation relationships calculated on $z$-normalized gene expression profiles across 2,059 highly variable genes. Genetic perturbations were ordered by HDBSCAN with a correlation metric calculated on the Illumina data (upper triangle) and the Ultima data (lower triangle) were ordered identically for comparison.

B) Comparison of perturbation relationships as derived from Ultima and Illumina sequencing data. Perturbation-perturbation relationships were calculated as described in fig. S13A. The cophenetic correlation between the Illumina and Ultima datasets is $r$=0.99.

C) Comparison of perturbation profiles derived from Ultima and Illumina sequencing. We computed the pairwise correlation of gene expression profiles of 2,017 genetic perturbations that elicited strong phenotypes. The mean correlation of profiles was $r$=0.73. Perturbations that elicited a higher number of differentially expressed genes (DEGs) (determined using a two-sample Anderson-Darling test compared against non-targeting guides) were better correlated across sequencing platforms (Spearman's rho=0.90).

**List of Supplementary Tables**

**Table S1.** K562 day 8 Perturb-seq sgRNA library table.

**Table S2.** K562 day 6 Perturb-seq sgRNA library table.

**Table S3.** RPE1 day 7 Perturb-seq sgRNA library table.

**Table S4.** K562 day 8 Perturb-seq pseudobulk summary statistics.

**Table S5.** K562 day 6 Perturb-seq pseudobulk summary statistics.

**Table S6.** RPE1 day 7 Perturb-seq pseudobulk summary statistics.

**Table S7.** Description of genetic perturbation clusters.

**Table S8.** Description of gene expression clusters.

**Table S9.** Description of plasmids.
